# Time-Resolved Tracking and Secretome Mapping of Intracellular, Cell Surface, and Extracellular Glycoconjugates with a Mini-Tetrazine *N*-Acetylglucosamine

**DOI:** 10.1101/2025.10.16.682811

**Authors:** Ashlyn S. Hillman, Mary W. N. Burns, Sarah L. Janney, Ashley M. Stark, Feng Cai, Timothy Chaya, Catherine L. Grimes, Jennifer J. Kohler, Joseph M. Fox

**Author notes:** Correspondence to: The University of Delaware, Department of Chemistry and Biochemistry, Newark, DE 19716,. Correspondence to: The University of Texas Southwestern Medical Center, Department of Biochemistry, Dallas, TX 75390,. Contributed equally to this work (Ashlyn S. Hillman, Mary W. N. Burns).

## Abstract

Glycosylation is instrumental in governing essential biological processes that dictate health outcomes. Herein we describe a mini-tetrazine *N*-acetylglucosamine (HTz-GlcNAc) that is incorporated into *N-*linked glycans and through rapid live-cell labeling with *trans*-cyclooctene (TCO) reagents enables real-time visualization and evaluation of glycoconjugate trafficking in living cells. We utilize cell-permeable and impermeable TCO-fluorophores to perform time-resolved, multicolor live labeling of cell-surface glycoconjugates, as well as intracellular, and internalized glycoconjugates. We observe intracellular glycoconjugate trafficking throughout the secretory pathway of live cells in real-time, as well as cell surface glycan turnover and extracellular vesicle secretion across three days. In the toolkit of probes for metabolic oligosaccharide engineering, HTz-GlcNAc brings the capability of precise interrogation of intracellular and extracellular glycoconjugate trafficking, and selective tagging of intra-and-extracellular vesicles, while providing a handle for the purification and enrichment of glycans for immunoblot and proteomic analyses.

TOC image

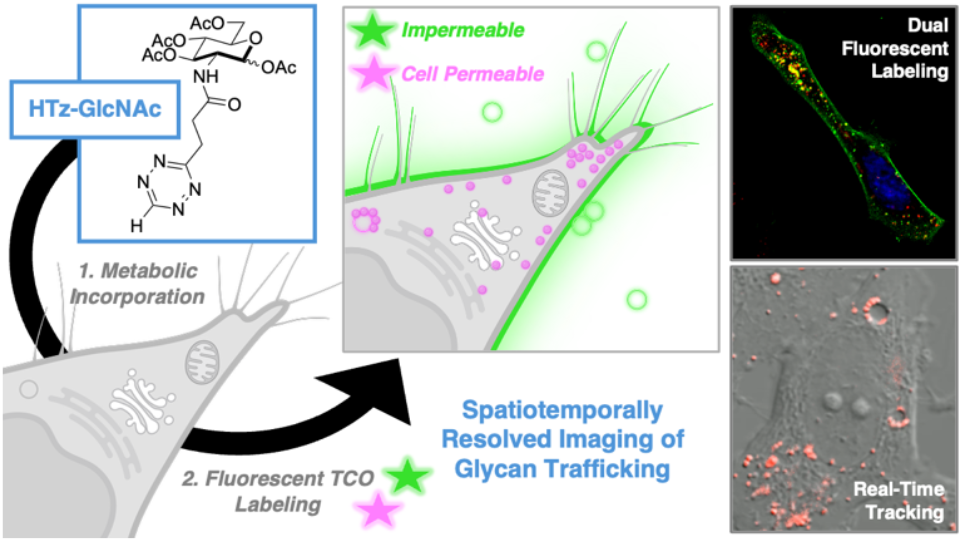

## Main

As one of the most abundant post-translational modifications, glycosylation is an essential regulator of human health.^1, 2^ Glycans are critical in facilitating many cellular processes, serving as molecular recognition sites for the immune system and in regulating cell signaling, growth, division, and adhesion.^3-4^ Dysregulation of glycan-mediated processes has been implicated in significant adverse health outcomes including autoimmune and inflammatory conditions, neurodegenerative diseases, and cancer.^1,3,5^ Furthermore, congenital disorders of glycosylation result in multi-system diseases.^6^ Therefore, tools are needed to investigate aberrant glycosylation as well as to characterize localization, structure, and function of glycans in cell biology to enable development of effective therapeutic treatments.

Metabolic oligosaccharide engineering (MOE) has enabled groundbreaking explorations in glycobiology through the incorporation of tagged glycan building blocks into cellular glycoconjugates including glycoproteins, glycolipids, and glycoRNA.^7-12^ In mammalian cells, MOE utilizes endogenous carbohydrate metabolic pathways for *N*-acetyl-glucosamine (GlcNAc), *N*-acetyl-galactosamine (GalNAc), and sialic acid to produce the corresponding modified activated nucleotide sugars: uridine-diphosphate-GlcNAc (UDP-GlcNAc), uridine-diphosphate-GalNAc (UDP-GalNAc), and cytidine-monophosphate-sialic acid (CMP-sialic acid) analogs.^13, 14^ Modified nucleotide-sugars serve as donor substrates for glycosyltransferases, the majority of which reside in the secretory pathway and decorate biomolecules through *O*-linked and *N*-linked glycosylation.^1, 5^ In this way, MOE enables production of modified glycoconjugates whose ultimate fate is to reside in the secretory pathway, be displayed on the cell surface, or be secreted extracellularly.

Bioorthogonal chemistry has emerged as a revolutionary method for glycobiology, as it encompasses a class of reactions that can proceed rapidly and selectively without perturbing endogenous functional groups and processes. ^15, 16^ Azide- and alkyne-tagged sugars have become staples in the field, as their small size enables enzyme permissibility, resulting in high metabolic incorporation.^13, 17, 18^ Azide- and alkyne-modified sugars allow for copper-catalyzed or strain-promoted cycloadditions (CuAAC or SPAAC) for fixed and live-cell imaging and glycoproteomic analysis of cell-surface glycans.^17,19-21^ Sugars have also been modified with a variety of functional group reporters to be used as MOE tools with applications extending to higher organisms including zebrafish and mice.^22-24^ However, due to modest labeling kinetics, the ability to visualize spatiotemporal changes in glycoconjugate movement throughout the secretory pathway remains elusive.

Inverse electron demand Diels-Alder reactions (IEDDA) with tetrazines (Tz) have figured prominently in the MOE toolkit with glycan reporters bearing cyclopropene,^25-27^ isonitrile,^28^ a-olefin,^29^ norbornene,^30^ bicyclononyne^31^ and *trans-*cyclooctene (TCO)^32^ tags that can be labelled with Tz-fluorophores with applications that extend to multicolor labeling based on orthogonal bioorthogonal tagging.^33, 34^ IEDDA reactions proceed rapidly with second-order rate constants exceeding 10^5^ M^-1^ s^-1^ when *trans-*cyclooctene (TCO) is the dienophile, ^16, 35, 36^ Recently, 3-methyl-6-aryltetrazine conjugates of tetraacetylmannosamine, tetraacetylgalactosamine, and tetraacetylsialic acid were used in cell surface labeling of MCF-7 cells, whereas noncancerous L929 fibroblasts did not accept these probes.^37^ Phospholipid liposomes were used to deliver a 3-methyl-6-aryltetrazine sialic acid conjugate to a broader range of cell lines and to cancer cells in vivo.^38^ While the history of MOE has been rich, these tools have been largely applied to cell-surface visualization of glycans. Elegantly, fluorescent fusion proteins have been used for the intracellular visualization of glycoproteins through Förster resonance energy transfer-fluorescence lifetime imaging microscopy (FRET-FLIM).^39^ Still, new tools are needed for the real-time visualization of glycoconjugates that enable study of spatiotemporal changes throughout the secretory pathway.

The efficiency of MOE is often sensitive to steric effects, with precursors bearing small bioorthogonal handles (e.g. alkynes, alkenes, cyclopropenes) serving as the most general tools. Bioorthogonal reporters bearing bulkier tetrazine, cyclooctyne or TCO handles can provide higher reactivity but at the expense of incorporation efficiency.^40^ Structure-guided mutagenesis of GlcNAc-1-phosphate uridyltransferase, AGX1, has been used to design a mutant AGX1 for efficient production of a diazirine-tagged UDP-GlcNAc.^41^ Schumann and colleagues have further developed a system that circumvents the steric bottlenecks of endogenous sugar salvage pathways via transfection of an artificial pathway for producing UDP-sugars from bioorthogonally tagged sugars, utilizing an AGX1 mutant and a promiscuous bacterial GlcNAc kinase (NahK). ^42, 43^ This system was coined Bio-Orthogonal Cell line-specific Tagging of Glycoproteins (BOCTAG) and has proven to be more tolerant to bulkier bioorthogonal tags appended to the *N*-acetyl group of 2-aminosugars.^44^

We have recently shown that a “mini” Tz *N*-acetyl-muramic acid can traverse peptidoglycan recycling and synthetic enzymes to be metabolically incorporated into the peptidoglycan of bacteria.^45, 46^ A hydrophilic axial-hydroxyl-TCO (aTCO) conjugated to a cell permeable fluorogenic Si-rhodamine (SiR) dye allowed for rapid, real-time, live-cell labeling of Tz-remodeled bacterial peptidoglycan during macrophage mediated phagocytosis.^45, 47, 48^

Here we report a mini-Tz-GlcNAc (HTz-GlcNAc, **Figure 1A**) that can be selectively metabolically incorporated into *N*-linked glycoproteins of mammalian BOCTAG cells. The two-enzyme BOCTAG platform, NahK and AGX1^F383A^, permits production of UDP-HTz-GlcNAc, which is utilized by secretory glycotransferases to incorporate HTz-GlcNAc into *N*-linked glycans. Selection of suitable TCO-fluorophores allows for differential labeling of extracellular and cell-surface glycans with impermeable TCO-AF488, and intracellular glycoconjugates with cell-permeable aTCO-SiR. The rapid labeling kinetics with TCO afford visualization and enrichment of glycoconjugates in immunoblot and proteomic analyses. We capture cell-surface glycan turnover and extracellular vesicle production across 3 days by utilizing a panel of impermeable TCO-fluorophores to selectively label extracellular and cell surface glycans at specific timepoints and monitor the fluorophores’ subcellular locations with live-cell confocal microscopy. Labeling with cell-permeable aTCO-SiR allows for visualization of intracellular HTz-GlcNAc metabolite trafficking in the secretory pathway (**Figure 1B**). Live-cell, real-time time-course imaging reveals labeling localized to ∼100 nm vesicles traveling toward the poles of the cell. Additionally, we see recruitment and organization of these small secretory vesicles around the perimeter of larger micron-sized structures. Experiments varying the introduction time of brefeldin A, an inhibitor that stops protein transport from the ER to the Golgi, resulted in exposure-dependent alterations in HTz-GlcNAc-tagged secretory vesicle formation and movement in the cell. This finding demonstrates HTz-GlcNAc’s utility in analyzing spatiotemporal alterations in glycan trafficking and synthesis that result from the introduction of stimuli. Overall, HTz-GlcNAc exhibits promise to serve as a live-cell tool for advancing glycoscience research and our understanding of how aberrant glycosylation results in the etiology of different diseases and disease pathogenesis. More broadly, HTz-GlcNAc is a versatile microscopy and enrichment tool for studying the secretory pathway, vesicular formation and transport, extracellular vesicles, and cell-cell interactions.

**Figure 1.**
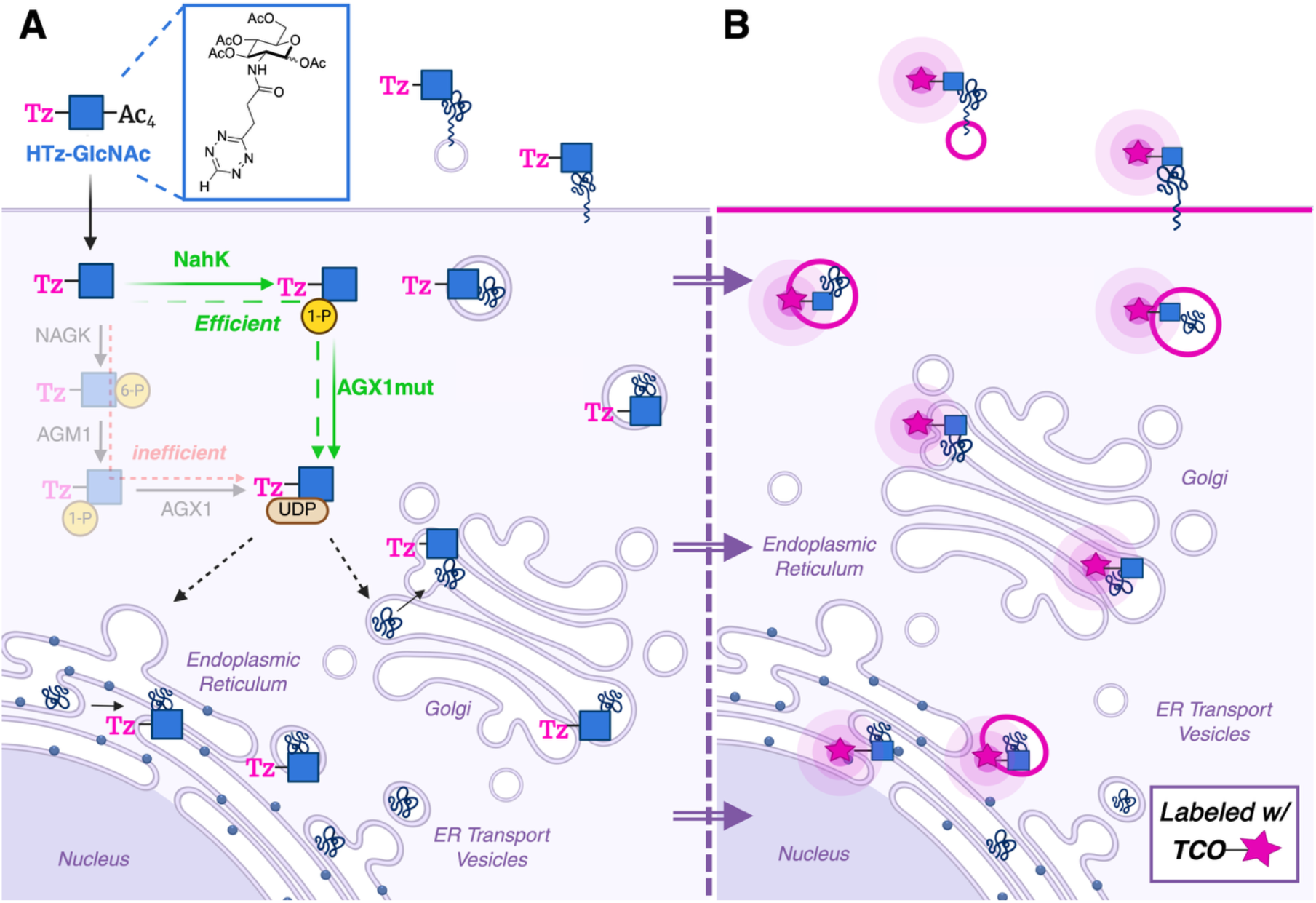
Metabolic Oligosaccharide Engineering with HTz-GlcNAc. A) Peracetylated HTz-GlcNAc passively diffuses across the plasma membrane and endogenous esterases remove the acetyl groups. HT-GlcNAc is too sterically encumbered to be an effective substrate for endogenous GlcNAc salvage pathway enzymes. Alternatively, the artificial two-enzyme salvage pathway (BOCTAG) utilizing NahK and AGX1^F383A^ can convert HTz-GlcNAc to UDP-HTz-GlcNAc, which is utilized by glycosyltransferases in glycoconjugate biosynthesis in the ER and Golgi. Mature HTz-GlcNAc-modified glycoconjugates are displayed on the cell surface and secreted into the extracellular environment. B) Rapid live-cell labeling with cell-permeable aTCO-SiR allows for tracking of intracellular secretory pathway glycoconjugate trafficking in real-time.

## Results

### Mini-Tetrazine Carbohydrates as Substrates for Metabolic Oligosaccharide Engineering

Multiple synthetic routes were successful in accessing mini Tz-*N*-acetylhexosamine (HexNAc) analogs, HTz-GlcNAc (SI Fig 1) and HTz-GalNAc (SI Fig 2), in one to two steps from the corresponding aminosugar tetraacetate hydrochloride salts (**Figure 2A**). HTz-HexNAc analogs could be prepared in one step from the 2-aminosugars via direct amide coupling with an HTz acid or pentafluorophenyl ester, which were prepared as previously described.^45^ Alternatively, to avoid volatile Tz precursors, we utilized the streamlined one-pot synthesis of thiomethyl-Tz (SMe-Tz) from carboxylic ester precursors to prepare *t-*butyl 3-(6-methylsulfanyl-1,2,4,5-tetrazin-3-yl)propanoate.^49^ This was hydrolyzed on gram-scale to the bench-stable 3-(6-methylsulfanyl-1,2,4,5-tetrazin-3-yl)propanoic acid. This acid or its corresponding activated ester was then coupled to the 2-aminosugars to access SMe-Tz-GlcNAc (SI Fig 1) and SMe-Tz-GalNAc (SI Fig 2). Then, the corresponding HTz-HexNAc analogs were achieved through late-stage palladium-catalyzed reduction.

**Figure 2.**
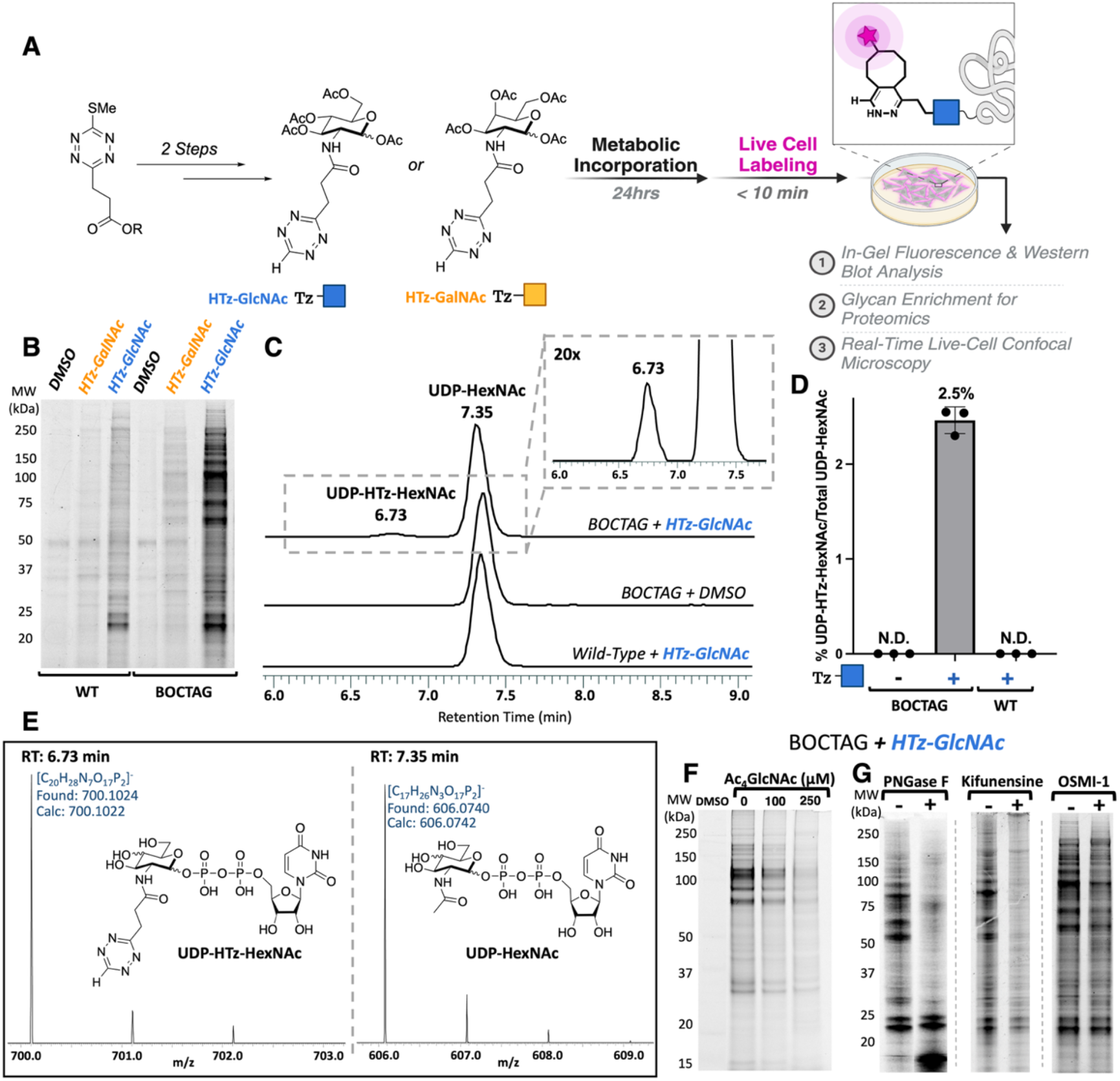
Development of Mini-Tetrazine Carbohydrates for Metabolic Glycome Tagging. A) Synthesis of HTz-GlcNAc & HTz-GalNAc in two steps from the glucosamine and galactosamine tetraacetate hydrogen chloride salts. SI Fig 1 & 2 for full synthetic schemes. Metabolic Incorporation Workflow: BOCTAG and WT SaOS2 osteosarcoma cells treated with HTz-Sugar for 24 hr, washed, followed by rapid live-cell TCO labeling; aTCO-SiR was used for in-gel fluorescence analysis. B) In-gel fluorescence analysis of HTz-GalNAc and HTz-GlcNAc metabolic incorporation in SaOS2 WT & BOCTAG cells. C) Targeted metabolomics analysis of UDP-HTz-HexNAc production on a Thermo Fisher Scientific Orbitrap Fusion Lumos Mass Spectrometer D) Estimated percentage of UDP-HTz-HexNAc present in BOCTAG and WT cell lysates normalized to endogenous UDP-HexNAc after 20 hr incubation. E) Orbitrap high-resolution mass spectrum of BOCTAG + HTz-GlcNAc chromatogram peaks: 6.73 min correlates to mass of UDP-HTz-HexNAc and 7.35 min correlates to mass of UDP-HexNAc F) HTz-GlcNAc competition assay with Ac_4_GlcNAc in SaOS2 BOCTAG cells. Vehicle Control (**-**) was treated with an equivalent volume of DMSO. G) Assessment of HTz-GlcNAc incorporation into glycans in the secretory pathway of SaOS2 BOCTAG cells via in-gel fluorescence after labeling with TCO-fluorophore, cell-impermeable TCO-647 for PNGaseF gel and cell-permeable aTCO-SiR for OSMI-1 and kifunensine gel. PNGaseF treatment was performed on cell lysate. OSMI-1 or kifunensine was included along with HTz-GlcNAc in culture media during metabolic incorporation.

With HTz-GlcNAc and HTz-GalNAc in hand, we assessed their utility for labeling mammalian glycans in human cells by culturing wild type (WT) osteosarcoma SaOS-2 cells in media containing the compounds. After 24 hours, live-cell labeling was performed using cell-permeable aTCO-SiR and labeled proteins were analyzed by in-gel fluorescence (**Figure 2A**). We did not observe strong fluorescent signal, suggesting that WT cells do not efficiently metabolically incorporate HTz-GlcNAc and HTz-GalNAc (**Figure 2B**). Endogenous HexNAc salvage machinery, comprised of GlcNAc kinase (NAGK), GlcNAc-6-phosphate mutase (AGM1), and AGX1, is a known bottleneck in the metabolic incorporation of bulkier HexNAc analogs. ^41, 43^

To circumvent the steric intolerances observed with the endogenous HexNAc salvage pathway, we used the BOCTAG system to introduce an artificial salvage pathway amenable to larger substrates.^43^ The BOCTAG system bypasses endogenous salvage pathway enzymes by using a promiscuous bacterial kinase, NahK, to phosphorylate the anomeric hydroxyl of HexNAc analogs. Then a mutant AGX, AGX1^F383A^, which harbors a mutation that enlarges the active site to permit bulkier HexNAc substrates, facilitates the transfer of a uridinyl phosphate group to produce the UDP-HexNAc analog (**Figure 1A**). Using SaOS-2 cells stably expressing the BOCTAG system, we performed live-cell labeling using aTCO-SiR. Labeling was evaluated by in-gel fluorescence, and we observed BOCTAG-dependent labeling for both HTz-GlcNAc and HTz-GalNAc (**Figure 2B**). At this point, we prioritized HTz-GlcNAc due to the stronger labeling profile compared to HTz-GalNAc; the lower HTz-GalNAc labeling efficiency suggested the possibility of an additional barrier to HTz-GalNAc incorporation.

Using an in-gel fluorescence assay, we observed dose-dependent labeling by HTz-GlcNAc (SI Fig 3). We observed labeling with as low as 1 µM HTz-GlcNAc, and the fluorescence signal increased with higher concentrations, plateauing in intensity at around 50 µM, the concentration we chose to use in subsequent experiments. HTz-GlcNAc showed no overt cytotoxicity; we quantified the effect of HTz-GlcNAc on WT and BOCTAG viability and found that it had no impact on cell viability even at the highest tested concentration (100 µM) (SI Fig 4).

### UDP-HTz-GlcNAc Production in BOCTAG Cells

Although the BOCTAG-dependence of efficient HTz-GlcNAc labeling suggested that the incorporation is dependent on UDP-HTz-GlcNAc generation, we sought additional evidence that the labeling was the result of enzymatic processes. This is particularly important given that peracetylated *N*-acetyl sugar analogs can non-enzymatically modify cysteine residues, via a process that does not require UDP-HexNAc generation.^50^

Initially, we used untargeted metabolomics to test whether culturing BOCTAG cells with HTz-GlcNAc led to production of UDP-HTz-HexNAc. Positive and negative ions corresponding to UDP-HTz-HexNAc fragments were detected in LC/MS and LC/MS/MS, and a third fragmentation was performed in positive ion mode to generate LC/MS/MS/MS data, further confirming the existence of UDP-HTz-HexNAc (SI Fig 5 & 6). Similar analysis was performed for UDP-HexNAc (SI Fig 7 & 8). We performed targeted metabolomics in triplicate using BOCTAG cells cultured with HTz-GlcNAc or DMSO and WT cells cultured with HTz-GlcNAc (**Figure 2C**). We detected UDP-HexNAc-modified with HTz in BOCTAG cells cultured with HTz-GlcNAc, while UDP-HTz-HexNAc was not detected in BOCTAG cells cultured with DMSO or in WT cells cultured with HTz-GlcNAc (**Figure 2C & 2E**). We could not measure the response factor of UDP-HTz-GlcNAc because we did not have a synthetic standard; to estimate UDP-HTz-GlcNAc concentration, we assumed that the response factors of UDP-GlcNAc and UDP-HTz-GlcNAc were equivalent. We quantified the peak intensities acquired in negative mode and estimated that 2.5% of cellular UDP-HexNAc is HTz-modified (**Figure 2D**). Positive mode quantification yielded a similar estimate (2.1% of HTz-modified cellular UDP-HexNAc). The BOCTAG-dependent labeling, as well as the BOCTAG-dependent UDP-HTz-HexNAc generation, suggests that HTz-GlcNAc is enzymatically incorporated into glycans and that the observed labeling is not a result of non-enzymatic cysteine labeling. The metabolomic results also confirmed that WT cells cultured with HTz-GlcNAc can serve as an appropriate control to account for non-specific and non-enzymatic HTz-GlcNAc labeling, since UDP-HTz-HexNAc production was not detected.

### Incorporation of HTz-GlcNAc into N-linked Glycans

We further characterized the glycoconjugates into which HTz-GlcNAc is incorporated. To probe the specificity of HTz-GlcNAc incorporation, we cultured cells with HTz-GlcNAc along with Ac_4_GlcNAc. Increasing the concentration of Ac_4_GlcNAc diminished labeling by HTz-GlcNAc in a dose-dependent manner, suggesting that BOCTAG cells were incorporating HTz-GlcNAc in a similar manner to salvaged, unmodified GlcNAc (**Figure 2F**).

HTz-GlcNAc-treated BOCTAG cell lysates were treated with PNGase F to selectively cleave N-glycans.^51^ PNGase F digestion resulted in a dramatic reduction in fluorescent labeling, indicating that HTz-GlcNAc is incorporated into N-glycans (**Figure 2G**). Interestingly, HTz-GalNAc-labeling was also reduced with PNGase F digestion, even though GalNAc is mainly present in O-glycans (SI Fig 9). This result suggests that HTz-GalNAc might be epimerized to HTz-GlcNAc by GALE and incorporated as HTz-GlcNAc.

We also cultured BOCTAG cells with kifunensine, an inhibitor of complex and hybrid N-glycan maturation, and we saw a dramatic reduction in HTz-GlcNAc labeling, further supporting that HTz-GlcNAc is incorporated into N-linked glycans (**Figure 2G &** SI Fig 10).^52^ The effect of kifunensine treatment on labeling also suggests that the majority of HTz-GlcNAc is incorporated into the antennae of mature N-glycans and not in the chitobiose core. We used lectin blots to assess the impact of HTz-GlcNAc on terminal fucose and sialic acid modifications (SI Fig 11). While no statistically significant impact on fucosylation or sialylation was detected in this assay, it remains possible that HTz-GlcNAc incorporation may impact further maturation of N-linked glycans.

Certain HexNAc analogs have been shown to be metabolically incorporated in the form of the O-GlcNAc modification on cytosolic and nuclear proteins through the action of O-GlcNAc-transferase (OGT).^53-55^ To investigate whether HTz-GlcNAc is metabolized through this pathway, BOCTAG SaOS-2 cells were cultured with HTz-GlcNAc in the presence of a small molecule OGT inhibitor, OSMI-1.^56^ An immunoblot with O-GlcNAc-recognizing antibody confirmed that samples treated with OSMI-1 displayed a statistically significant reduction in O-GlcNAc levels indicating successful OGT inhibition (SI Fig 12). Fluorescent labeling with aTCO-SiR and in-gel fluorescence analysis showed a no significant reduction in labeling in OSMI-1 treated cells (**Figure 2G &** SI Figure 12) suggesting that little to no HTz-GlcNAc is incorporated into O-GlcNAc modifications.

### Adaptable Workflows for the Enrichment and Proteomic Detection of Low Abundance Secreted Glycoproteins

To learn which proteins are decorated with HTz-GlcNAc-modified glycans we used TCO-biotin to label whole cell lysates prepared from WT and BOCTAG cells cultured with HTz-GlcNAc. Biotin-labeled proteins were enriched with neutravidin resin, then subjected to on-bead trypsin digestion, followed by LC/MS to identify and quantify unmodified peptides (**Figure 3A**). We performed label-free quantitation based on precursor ion intensity, and calculated fold-enrichment of proteins identified from BOCTAG cells as compared to proteins identified from WT cells. The majority of significantly enriched hits were cell-surface glycoproteins with known N-glycosylation sites, such as LAMP1 and ITGB1, further supporting that HTz-GlcNAc is incorporated into N-glycans and selectively labels proteins that traffic through the secretory pathway (**Figure 3B**). To further validate the proteomics results as well as determine if glycoproteins could be purified through HTz-GlcNAc, we performed TCO-biotin labeling and neutravidin enrichment followed by immunoblotting for representative glycoproteins LAMP1 and ITGB1 and found that they were highly enriched (**Figure 3C**). The enrichment was BOCTAG-dependent, demonstrating that UDP-HTz-GlcNAc generation is necessary for efficient glycoprotein labeling using HTz-GlcNAc.

**Figure 3.**
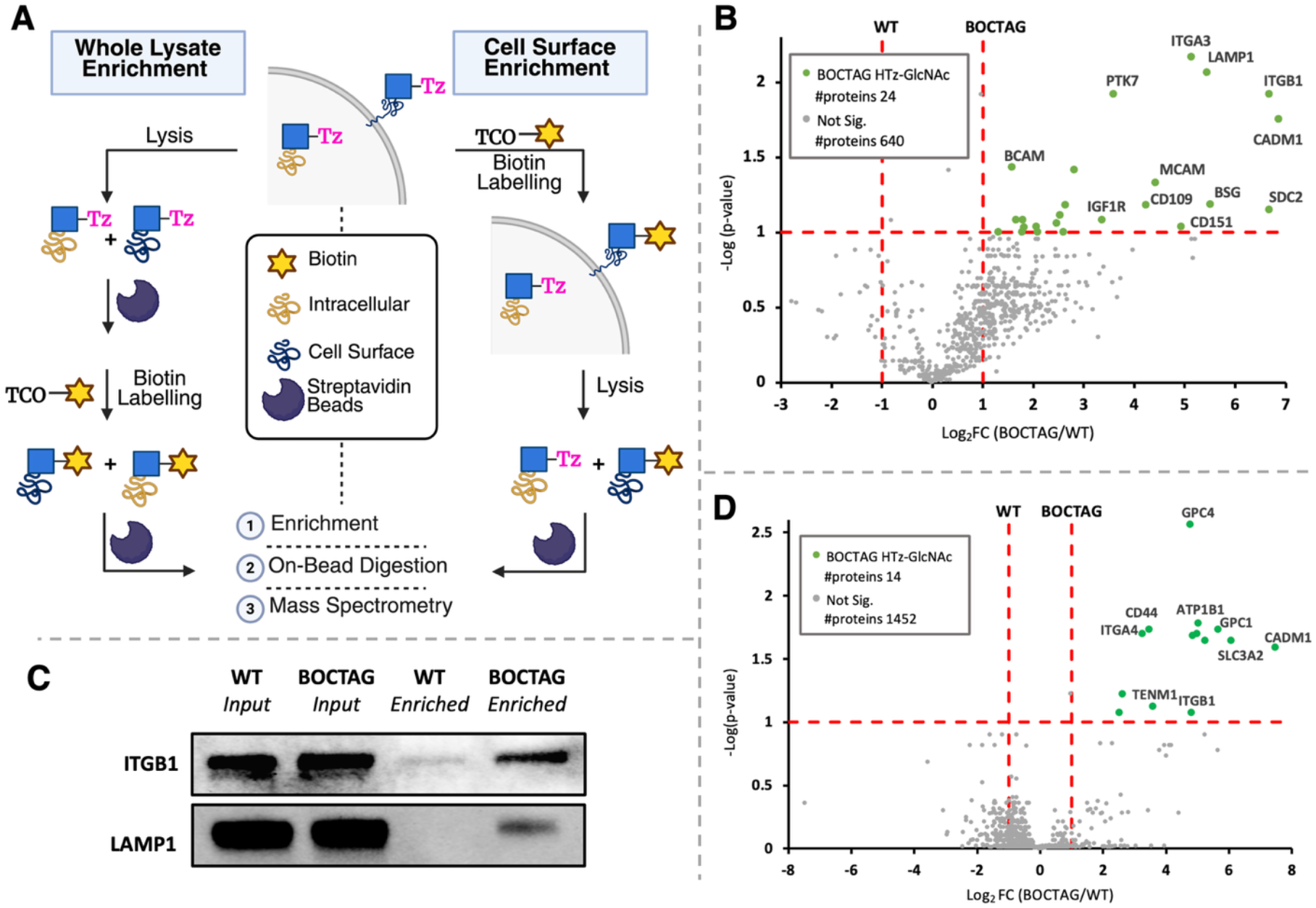
Enrichment of Secretory Glycoproteins with HTz-GlcNAc for MS Analysis. A) Labeling workflow for whole-lysate enrichment vs. cell surface enrichment with HTz-GlcNAc B) Volcano plot of protein enrichment achieved through whole lysate labeling of BOCTAG SaOS-2 cells cultured with 50 µM HTz-GlcNAc compared to WT SaOS-2 cells cultured with 50 µM HTz-GlcNAc. C) Immunoblot analysis of enrichment of ITGB1 and LAMP1 via HTz-GlcNAc labeling in SaOS-2 BOCTAG Cells D) Volcano plot of protein enrichment achieved via cell surface labeling of BOCTAG SaOS-2 cells cultured with 50 µM HTz-GlcNAc compared to WT SaOS-2 cells cultured with 50 µM HTz-GlcNAc. See SI Fig 13 for volcano plots comparing HTz-GlcNAc labeling to the DMSO controls for both WT and BOCTAG cells.

We also performed labeling on intact cells with an impermeable TCO-biotin to selectively label cell-surface glycoconjugates. Samples were enriched and analyzed by proteomics as above (**Figure 3A**). All significantly enriched hits were annotated to be glycoproteins carrying N-glycans, and many of the hits were present in the whole lysate proteomics results as well. By selectively labeling the cell surface, we captured GPI-anchored proteoglycans, such as GPC1 and GPC4, that are also modified with N-linked glycans. (**Figure 3D**). These labeling workflows demonstrate that N-linked glycoproteins of interest can be selectively targeted with TCO-reagents to enhance sensitivity of glycoprotein detection by proteomics.

### Differential Labeling of Intracellular and Extracellular Glycoconjugates

After characterizing HTz-GlcNAc labeling by biochemical methods and confirming enzymatic incorporation into N-glycans, we tested the utility of HTz-GlcNAc for live-cell imaging. First, we employed an impermeable TCO-dye (TCO-AF488) to selectively label HTz-GlcNAc-containing cell-surface glycans (**Figure 4A**). We observed BOCTAG-dependent extracellular labeling with the cell-impermeable fluorophore (**Figure 4B**), which is consistent with the metabolomics results indicating that BOCTAG salvage enzymes are required for UDP-HTz-GlcNAc production and incorporation into cell-surface glycans. Additionally, we performed flow cytometry to analyze cell-surface labeling and observed strong BOCTAG-dependent labeling with both HTz-GlcNAc (>50-fold above WT) and HTz-GalNAc (>10-fold above WT) (SI Fig 14). Subsequent labeling of BOCTAG cells with a cell permeable TCO-dye (aTCO-SiR) allows for differential dual labeling of cell-surface and intracellular glycoconjugates (**Figure 2C** and SI Video 1). We further confirmed BOCTAG-dependence of both cell surface and intracellular HTz-GlcNAc labeling by labeling cells with either cell-permeable or impermeable dyes and analyzing by in-gel fluorescence (SI Fig 9 & 12).

**Figure 4.**
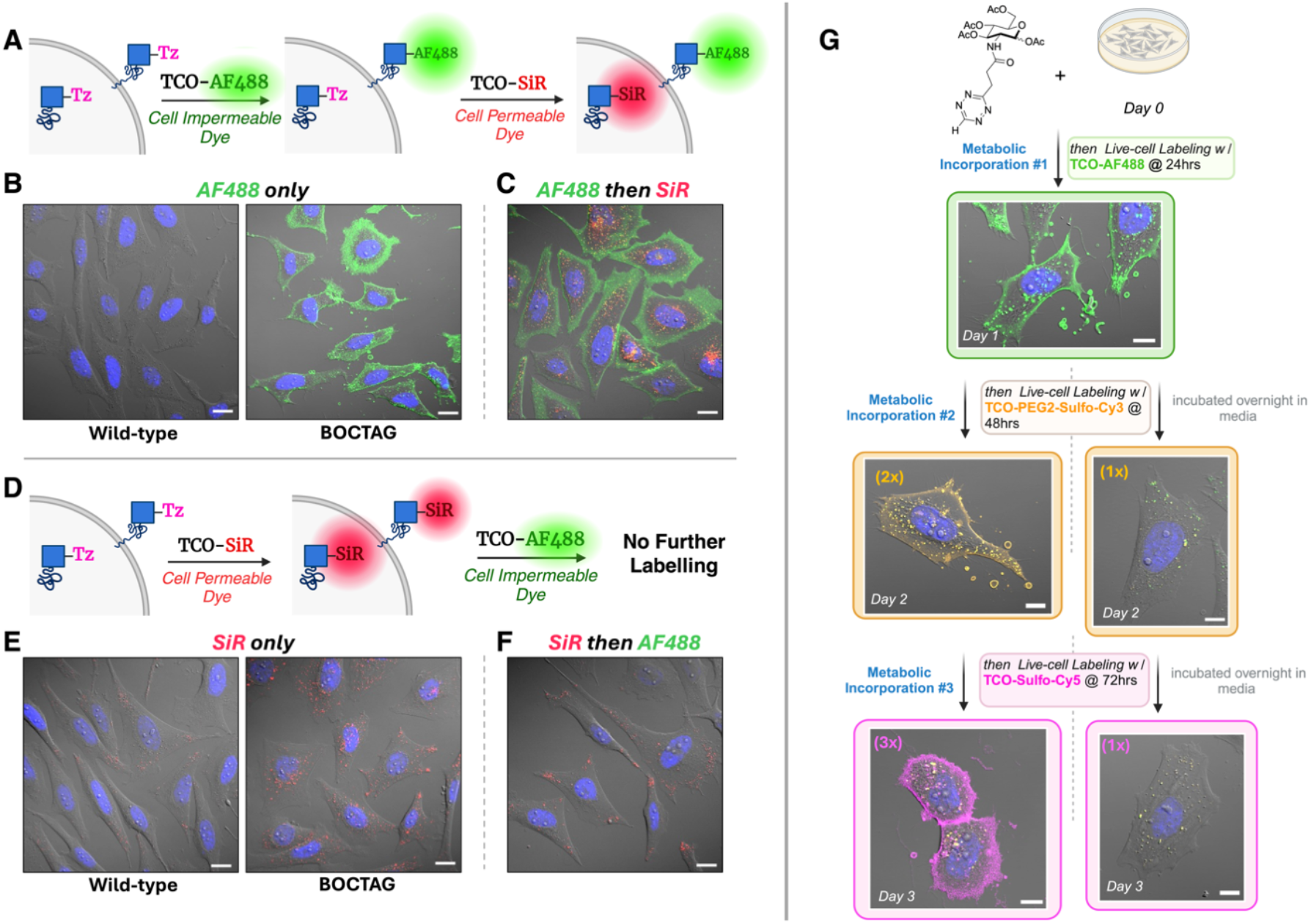
Live-cell differential visualization of cell surface glycan turnover, intracellular glycoconjugates, and extracellular vesicle secretion. A) Dual Differential Labeling Workflow: Following 24 hr MOE with HTz-GlcNAc, cells are live-labeled with a cell-impermeable dye TCO-AF488; (green), washed, and live-labeled with cell-permeable aTCO-SiR (red). B) Left: Wild-type SaOS2 cells cultured with 50 µM HTz-GlcNAc, then labeled with TCO-AF488. Right: BOCTAG SaOS2 cells cultured with 50 µM HTz-GlcNAc, then labeled with TCO-AF488. C) BOCTAG SaOS2 cells cultured with 50 µM HTz-GlcNAc, then labeled with TCO-AF488 followed by aTCO-SiR. D) Intracellular Labeling Workflow: Following 24 hr MOE with HTz-GlcNAc, cells are live-labeled with a cell-permeable dye aTCO-SiR (red), washed, and labeled with TCO-AF488 (green) Left: Wild-type SaOS2 cells cultured with 50 µM HTz-GlcNAc, then labeled with aTCO-SiR. Right: BOCTAG SaOS2 cells cultured with 50 µM HTz-GlcNAc, then labeled with aTCO-SiR. F) BOCTAG SaOS2 cells cultured with 50 µM HTz-GlcNAc, then labeled with aTCO-SiR followed by TCO-AF488. Scale bar is 20 µm. G) Time-point monitoring of HTz-GlcNAc incorporation by confocal microscopy over 3-days. Days 0-1) Overnight incubation with 50 µM HTz-GlcNAc; Metabolic Incorporation #1. Cells labeled with TCO-AF488 (green) at 24 hr. Days 1-2) (2x) Second consecutive overnight treatment with 50 µM HTz-GlcNAc for Metabolic Incorporation # 2; (1x) Overnight incubation in media. Both samples labeled with Sulfo-Cy3-Peg2-TCO (orange) at 48 hr. Days 2-3) (3x) Third consecutive overnight treatment with 50 µM HTz-GlcNAc for Metabolic Incorporation #3; (1x) Overnight incubation in media. Both samples labeled with Sulfo-Cy5-TCO (pink) at 72 hr. Scale bar is 10 µm.

Conversely, we found that if we reversed the order in which we labeled cells, first labeling with the cell-permeable aTCO-SiR followed by the cell-impermeable TCO-AF488, we did not observe significant cell-surface fluorescence (**Figure 4D-F**). This result indicated that aTCO-SiR was labeling both intracellular and extracellular glycoconjugates, with much brighter fluorescence from intracellular SiR. The brighter intracellular signal is likely due to the environmental dependence of SiR fluorescence, which derives from the equilibrium between a fluorescent zwitterion and a nonfluorescent spirolactone. ^57, 58^ This equilibrium is pH sensitive, with lower pH stabilizing the fluorescent zwitterion.^59^ This pH-dependent fluorogenicity has been shown to be advantageous for imaging acidic environments in live cells.^51^ As the organelles associated with glycoconjugates in the secretory and endolysosomal pathways are acidic, brighter fluorescence is expected from SiR dyes in those cellular environments.^60, 61^

### Visualization of Cell Surface Glycoprotein Turnover and Extracellular Vesicles

To evaluate HTz-GlcNAc as tool for spatiotemporal visualization of cell-surface glycan turnover, we performed a time course experiment in which HTz-GlcNAc containing glycans were labeled with different cell-impermeable TCO-fluorescent reporters every 24 hr and monitored by live-cell fluorescence confocal microscopy over the span of 3 days **(Figure 4G).** In this experiment, one set of BOCTAG cells was incubated once with HTz-GlcNAc for 24 hr, while another set of BOCTAG cells underwent three successive 24 hr treatments with HTz-GlcNAc. Both sets were fluorescently labeled at 24, 48, and 72 hr with three different colored cell-impermeable TCO-fluorophores, AF488 (green), Sulfo-Cy3 (orange), and Sulfo-Cy5 (pink) respectively.

Following TCO-AF488 labeling at 24 hr, we observed green-fluorescent labeling of extracellular vesicle membranes and the cell surface (**Figure 4G-Day 1 &** SI Fig 15). At 48 and 72 hr in BOCTAG cells that had undergone one treatment with HTz-GlcNAc, we observed internalization and progressive depletion of green-fluorescent signal, with no additional cell surface fluorescent labeling with Sulfo-Cy3 and Sulfo-Cy5 (**Figure 4G-Day 2 & 3** and SI Fig 16 & 17). In the BOCTAG sample that underwent daily treatment with HTz-GlcNAc, we observed internalization and reduction of the green-fluorescent signal at 48 hr, as well as Sulfo-Cy3 orange-fluorescent labeling of extracellular vesicle membranes and the cell surface (**Figure 4G-Day 2 &** SI Fig 18). At 72 hr, following the third and final treatment with HTz-GlcNAc we observed internalization and reduction of the orange-fluorescent signal, continued depletion of internalized green-fluorescent signal, along with Sulfo-Cy5 pink-fluorescent cell surface and extracellular vesicle membrane labeling (**Figure 4G-Day 3 &** SI Fig 19). Thus, the color of the label marks the age of the glycoconjugate.

In contrast, the sample that received only the initial dose of HTz-GlcNAc lacked fluorescent label on the cell surface after the first day, indicating that the cell surface fluorescence observed on day 2 and day 3 for cells that underwent daily HTz-GlcNAc treatment was from newly synthesized glycans. Consequently, daily metabolic incorporation of HTz-GlcNAc allowed us to visualize and monitor HTz-GlcNAc incorporation into newly expressed cell-surface glycans, as well as the secretion of new extracellular vesicles. This multiday live-cell monitoring also showed internalization of the previous day’s fluorescent signal, showing that glycoconjugates internalized from the cell surface were fully replaced with a newly biosynthesized HTz-labeled glycome.

### Real-Time Visualization of Intracellular Glycoconjugate Trafficking

While acquiring live-cell images of intracellular aTCO-SiR labeling (**Figure 4E**), we noticed rapid movement of fluorescently-labeled vesicular structures. We confirmed that HTz-GlcNAc could be used to image glycoconjugate movement in live cells in real-time through time-course video acquisition. Fluorescent vesicles were about 100-200 nm in diameter and tended to congregate to the cellular poles and projections, as well as around the nucleus. We observed real-time assembly of these small, labeled vesicles around the perimeter of larger structures (∼2 µm) within the cell (**Figure 5A** and SI Video 2). Additionally, HTz-GlcNAc metabolic incorporation and labeling with aTCO-SiR enabled visualization of glycoconjugate trafficking and secretory pathway dynamics during complex cellular processes like extracellular vesicle formation, membrane ruffling, and cell division (SI Video 3).

**Figure 5.**
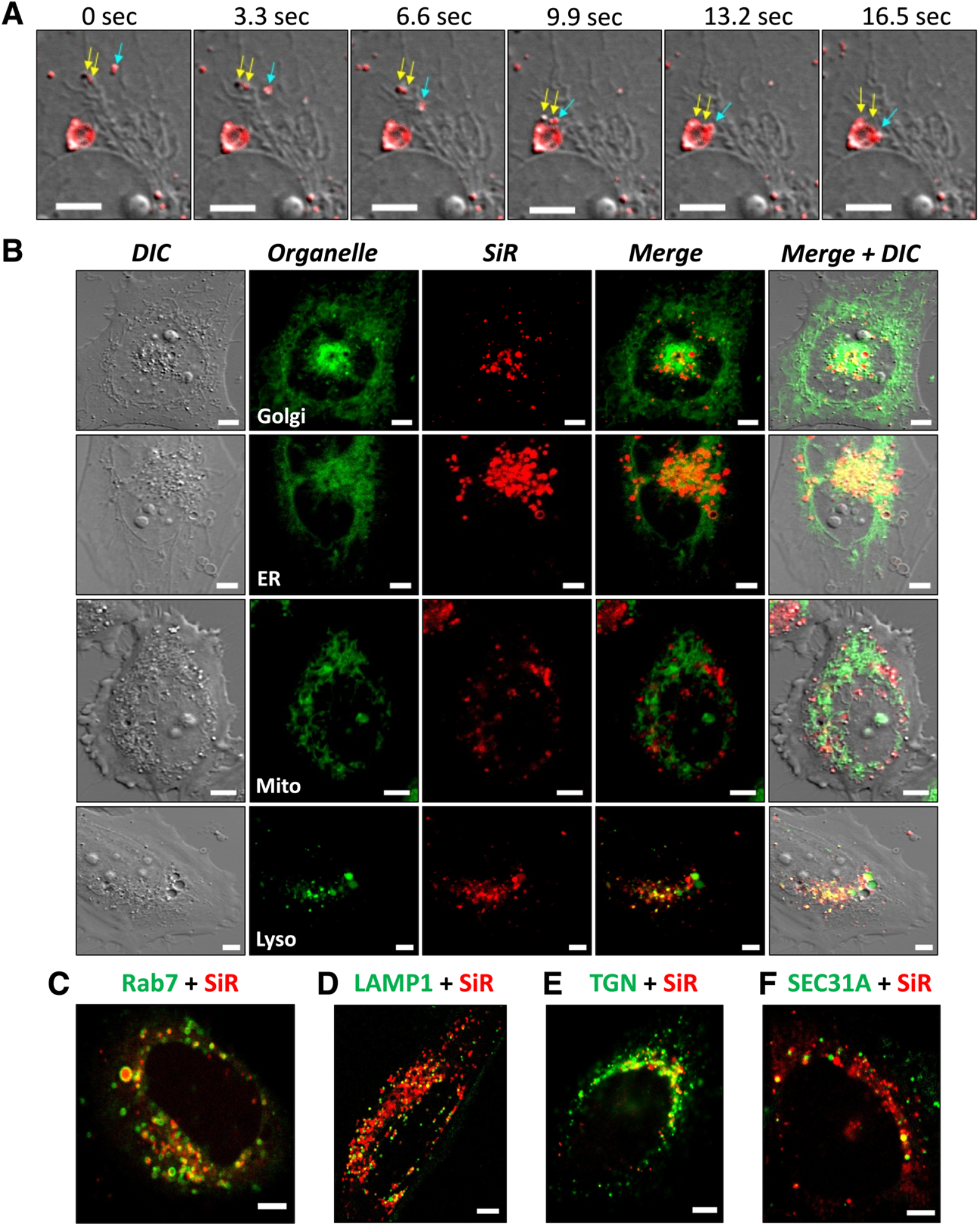
Real-Time Tracking of Secretory Pathway Glycan Trafficking. A) Live-cell real-time visualization of HTz-GlcNAc-tagged-SiR-labeled-vesicle (∼100 nm) organization around the perimeter of larger secretory vesicles (∼2 µm). B) Live cell colocalization of SiR-labeled glycans with BDP^®^ TMR Ceramide (Golgi), LumiTracker ER Green (endoplasmic Reticulum), LumiTracker Mito Orange CMTMRos (mitochondria), and LumiTracker Lyso Green (lysosomes). C) Live-cell colocalization of SiR-labeled glycans with vesicle markers: Rab7 (Endosomes), LAMP1 (Lysosomes), TGN (trans-Golgi network), and SEC31A (COPII). Scale bar, 5 µm.

Real-time monitoring following selective cell-surface and extracellular glycoconjugate labeling with cell-impermeable TCO-AF488 was consistent with what we saw in the 3-day time course experiment (**Figure 4G**), where we were able to observe internalization and depletion of cell-surface fluorescent signal in real-time (SI Video 4 & 5). Timelapse imaging over 7.5 hr showed that HTz-GlcNAc-tagged glycoconjugates were intimately associated with dynamic interactions between neighboring cells as well as cellular projections (SI Video 5). Additionally, we observed real-time encapsulation of a HTz-GlcNAc-tagged extracellular vesicle by a neighboring cell (SI Video 6), highlighting the potential utility of HTz-GlcNAc as a tool for isolating, tracking, and characterizing extracellular vesicles and analyzing their function in cell-cell communication.

### Secretory and Endolysosomal Pathway Specific Glycoconjugate Tagging

After observing HTz-GlcNAc-labeled intracellular structures, we used fluorescence confocal microscopy to identify which organelles were labeled. We measured HTz-GlcNAc colocalization with small molecule organelle markers followed by Pearson correlation coefficient (r) calculation and observed very weak colocalization with the mitochondria (r = 0.19), moderate colocalization with the endoplasmic reticulum (r = 0.44) and Golgi apparatus (r = 0.45), and strong colocalization with the lysosome (r = 0.75) (**Figure 5B**), consistent with labeling proteins in the secretory and endolysosomal pathways. To further dissect the types of vesicles that HTz-GlcNAc were visualized by fluorescent labeling of HTz-GlcNAc-containing glycoconjugates, we performed colocalization analysis with fluorescently-tagged marker proteins: Rab7 (endosome), LAMP1 (lysosome), SEC31A (COPII ER vesicles), and TGN46 (trans-Golgi network) (**Figure 5C)**. We observed the most HTz-GlcNAc labeling in Rab7-positive vesicles and, to a lesser extent, also observed labeling in LAMP1-, SEC31A-, and TGN46-positive vesicles as well. These results indicate that HTz-GlcNAc can be used to monitor the secretory pathway as well as cell-surface glycoprotein internalization in the endolysosomal pathway.

### Visualization of Stimuli-Induced Spatiotemporal Changes in Glycoconjugate Tracking

After imaging glycan labeling during homeostasis, we were curious if we could observe dynamic changes in HTz-GlcNAc-glycan trafficking in response to an external stimulus. To evaluate the potential utility of using HTz-GlcNAc as a tool for understanding the impact of stimuli on glycoconjugate trafficking, we treated cells with brefeldin A, a compound that inhibits anterograde transport through the secretory pathway and causes collapse of the Golgi into the endoplasmic reticulum.^62, 63^ To determine the impact of brefeldin A on HTz-GlcNAc-positive vesicular trafficking, we compared BOCTAG cells that had been treated with brefeldin A for the duration of the 24 hr HTz-GlcNAc incubation to cells that were not treated with brefeldin A. Here we observed that cells treated with brefeldin A for 24 hr exhibited bright and perinuclear intracellular fluorescence that was diffuse across the cell rather than the defined small vesicles that localize to the poles of cells not treated with brefeldin A. Additionally, we observed that HTz-GlcNAc-labeled intracellular vesicles treated with brefeldin A for 30 min just prior to imaging showed reduced migration to the poles of the cell and these vesicles localized more around the nucleus and endoplasmic reticulum (**Figure 6** and SI Video 7). These observations are consistent with secretory pathway dependence of HTz-GlcNAc labeling and show that we can use HTz-GlcNAc to learn how small molecules and other external stimuli perturb secretory pathway dynamics in real-time.

**Figure 6.**
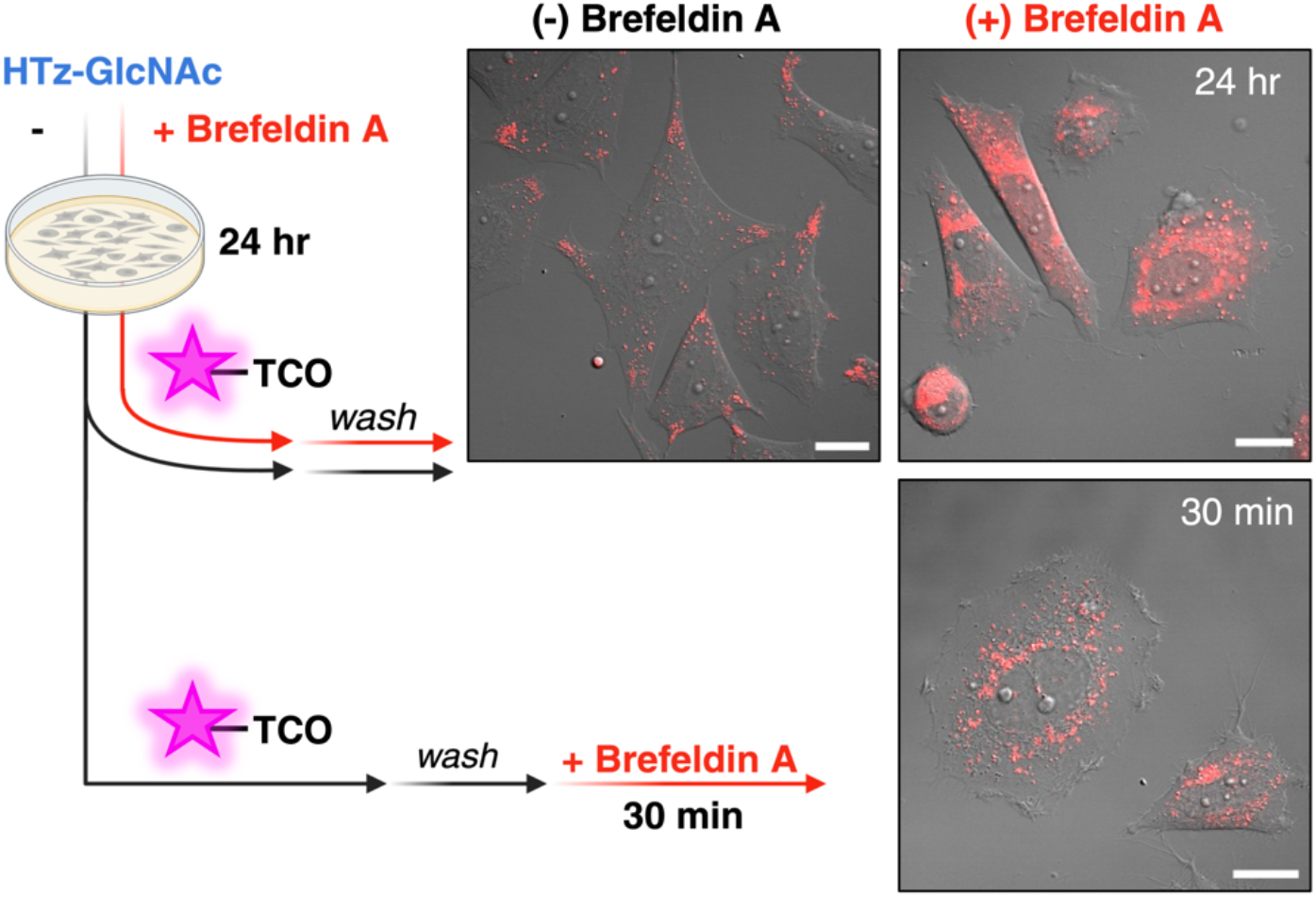
Chemical Stimuli Induced Temporal Alterations in Secretory Glycan Trafficking. BOCTAG cells were incubated with HTz-GlcNAc +/-brefeldin A for 24 hr. Cells were washed and labeled with aTCO-SiR and evaluated using live-cell fluorescence confocal microscopy. A third set of cells were incubated with HTz-GlcNAc for 24 hr, washed, labeled with aTCO-SiR, and incubated with brefeldin A for 30 min directly preceding confocal microscopy. Scale bar, 20 µm.

## Discussion

Here we report a mini-tetrazine *N*-acetylglucosamine, HTz-GlcNAc, as a next-generation “clickable” GlcNAc analog that is converted to UDP-HTz-GlcNAc in mammalian cells via an engineered GlcNAc salvage pathway, allowing HTz-GlcNAc to be incorporated into glycoconjugates. We demonstrate by enzymatic digestion and small molecule inhibition that HTz-GlcNAc is preferentially incorporated into N-glycans, making it an important addition to the limited N-linked glycan toolbox. HTz-GlcNAc can be used for glycoprotein labeling, enrichment, and live-cell imaging of intra- and extracellular glycoconjugates. We imaged intracellular glycoconjugate movement in real-time, monitored cell-surface glycan internalization and extracellular vesicle production across 3 days, tracked intracellular HTz-GlcNAc labeling in the secretory and endolysosomal pathways, and monitored the effect of an external stimulus on HTz-GlcNAc vesicle trafficking.

The BOCTAG dependence of HTz-GlcNAc labeling as well as the detection of UDP-HTz-HexNAc support the idea that labeling occurs through an enzymatic process. Our data do not directly address whether epimerization interconverting UDP-HTz-GlcNAc and UDP-HTz-GalNAc occurs. However, glycosidase sensitivity and small molecule inhibition studies demonstrate that the majority of proteins labeled with HTz-GlcNAc are N-linked glycoproteins. These results are consistent with studies of metabolism of other N-acyl-modified GlcNAc analogs, in which MGAT1, MGAT2, and MGAT5 are most tolerant of bulky modifications.^64, 65^ For example, the Hanover group recently reported a GlcNAc analog for CuAAC labeling, 1,3-Pr_2_-6-OTs GlcNAlk (MM-JH-01), that is selectively incorporated into mature N-glycans by MGAT1. Similar to HTz-GlcNAc, MM-JH-01 labeling displays sensitivity to PNGase F and kifunensine treatments.^64^

Proteomics analyses of HTz-GlcNAc-labeled proteins from whole-cell lysates revealed that HTz-GlcNAc preferentially labels N-linked glycoproteins, further confirming its specificity for N-linked glycans. Importantly, we found that we could selectively enrich lower abundance glycoproteins by selectively labeling the cell surface. We focused on glycoproteins in this study, but our work does not exclude the possibility that HTz-GlcNAc is also incorporated into other glycoconjugates, such as glycolipids, which will be investigated in future work.^64-67^

HTz-GlcNAc has broad potential applications due to its fast reaction kinetics and utility in real-time live cell imaging. HTz-GlcNAc enabled spatiotemporally-resolved visualization of changes in intracellular N-glycoconjugate trafficking through the secretory and endolysosomal pathways in response to an external stimulus (brefeldin A), serving as proof-of-concept for using HTz-GlcNAc as a tool to understand factors affecting glycan biosynthesis and turnover. In combination with fluorogenic aTCO-SiR, HTz-GlcNAc enables imaging of intracellular glycoconjugates, without the need to fix cells or wash away unreacted intracellular fluorophore, facilitating observation of real-time glycoconjugate dynamics in live cells. In the future, HTz-GlcNAc may be used to monitor N-glycoconjugate trafficking in response to other stressors or small-molecule probes. With appropriate detection reagents, one could envision using HTz-GlcNAc for whole-organism imaging to visualize N-glycoconjugate trafficking during different physiological states. HTz-GlcNAc is a versatile tool for microscopy and enrichment tool for studying the secretory pathway, vesicular formation and dynamics, cell-cell interactions, and for visualizing mammalian glycoconjugates in real-time.

## Acknowledgements

This work was supported by the NIH (5F30DK136209, R01 GM132460, with instrument support provided by P20GM104316, P20 GM103446, P20GM139760, S10OD025185, S10OD030321, S10OD016361), the Welch Foundation (I-1686), and the state of Delaware. This content is solely the responsibility of the authors and does not necessarily represent the official views of the NIH. We would like to thank Jeff Caplan, Sylvain Le Marchand, and the Delaware Biotechnology Institute Bioimaging Center. We would like to thank the UTSW Quantitative Light Microscopy Core where microscopy access was supported by an NIH grant (1S10OD021684-01). We would like to thank the UTSW Children’s Medical Center Research Institute Metabolomics Core which is supported by funding from the Cancer Prevention Research Institute of Texas (CPRIT Core Facilities Support Award RP240494). We would like to thank the UTSW Proteomics Core facility. We would also like to thank William Trout; Papa Nii Asare-Okai from the UD Mass Spectrometry Core Facility; and Shi Bai and the UD NMR Facility. A.S.H and A.M.S would like to thank the NIH for support through the Chemical-Biology Interface (CBI) training grant (T32GM133395). M.W.N.B would like to thank the UTSW Sara and Frank McKnight Fellowship Program for support. S.L.J would like to thank the University of Delaware Summer Scholars Program for support. J.M.F and A.S.H would like to acknowledge that this research was partially funded by the NSF through the University of Delaware Materials Research Science and Engineering Center (DMR-2011824). We thank Benjamin Schumann (Dresden University of Technology) for sharing the pSBbi-AGX1-NahK plasmid

