## Supporting Information for "Time-Resolved Tracking and Secretome Mapping of Intracellular, Cell Surface, and Extracellular Glycoconjugates with a Mini-Tetrazine *N*-Acetylglucosamine"

Table of Contents

1. Supplemental Figures……………………………………………………..........................S3
2. General Information: Materials and Methods..…......……………….............................S22
3. Synthetic Procedures...........................……………………….............................S23
4. Biochemistry Methods..……………………………………………..............................S30
   4.1 Flow Cytometry Measurement of Cell Surface HTz-GlcNAc and HTz-GalNAc..........................................................................................S30
   4.2 In Gel Fluorescence Analysis of HTz-GlcNAc and HTz-GalNAc-Labeled Glycoproteins..........................................................................................S30
   4.3 PNGaseF Release of N-Glycans from HTz-GlcNAc and HTz-GalNAc Labeled Glycoproteins..........................................................................................S31
   4.4 Kifunensine Treatment of BOCTAG Cells.....................................................S31
   4.5 OSMI-1 Treatment of BOCTAG SaOS-2 Cells...............................................S31
   4.6 Analysis of HTz-GlcNAc Impact on Fucosylation and Sialylation of Glycans in BOCTAG Cells..........................................................................................S32
   4.7 Analysis of HTz-GlcNAc Toxicity on WT and BOCTAG Cells with CellTiter-Glo..........................................................................................S32
   4.8 Metabolomics Sample Preparation.................................................................S32
   4.9 Metabolomics Analysis for Detection and Quantification of UDP-HTz-GlcNAc..........................................................................................S32
   4.10 Enrichment and Proteomics Analysis of HTz-GlcNAc-Labeled Proteins.........S33
   4.11 On-Bead Digestion, Proteomics Data Acquisition, and Peptide Identification and Quantification..........................................................................................S34
   4.12 Enrichment and Immunoblot Analysis of HTz-GlcNAc-Labeled Proteins........S35
   4.13 Live Cell Confocal Fluorescence Microscopy of HTz-GlcNAc.......................S35
   4.14 Dual Labeling Confocal Fluorescence Microscopy of HTz-GlcNAc...............S35
   4.15 Multiday Time-Point Labeling Confocal Fluorescence Microscopy of HTz-GlcNAc..........................................................................................S36
   4.16 Live Cell Confocal Fluorescence of HTz-GlcNAc with Vesicle Markers.........S37
   4.17 Live Cell Confocal Fluorescence Microscopy of HTz-GlcNAc with Brefeldin A Treatment..........................................................................................S37
5. References………………………………..………………...............................S37
6. $^{1}$H, $^{13}$C, $^{19}$F, 2D NMR Spectra…………..………………………..........................S39

### Supplemental Figures

*Route 1:*

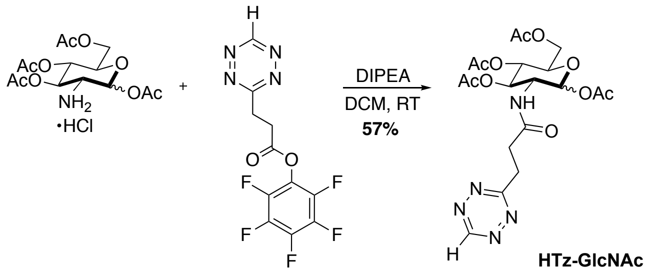


*Route 2:*

*Route 3:*

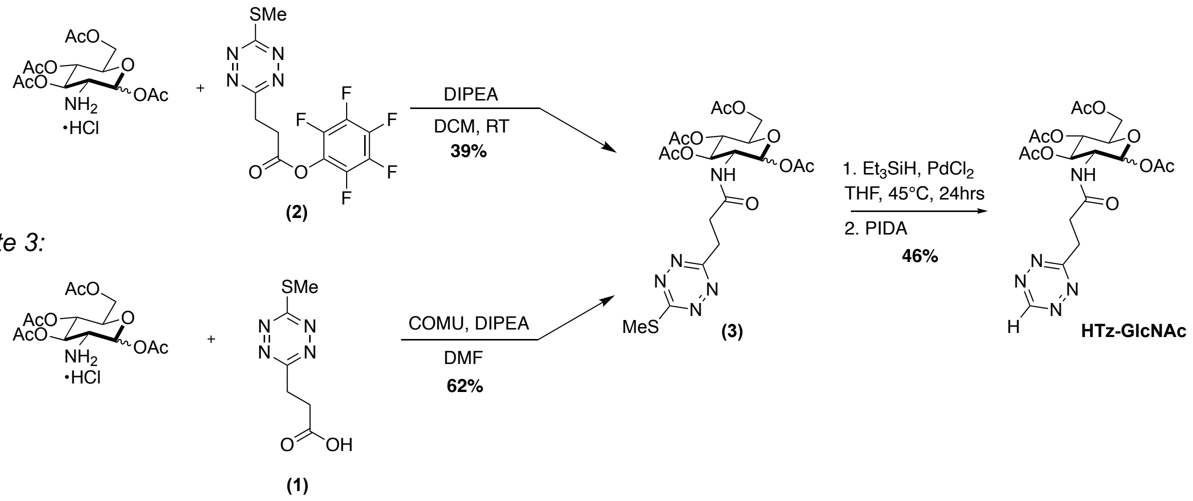


**Supplemental Figure 1. HTz-GlcNAc Synthetic Routes.**

*Route 1:*

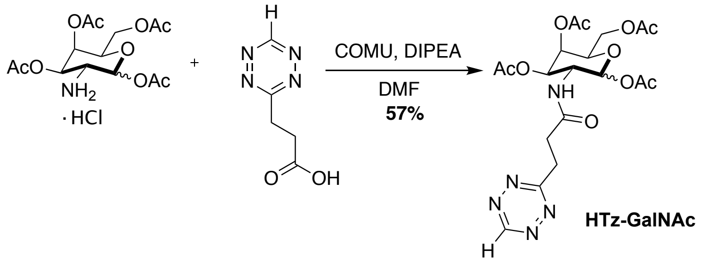


*Route 2:*

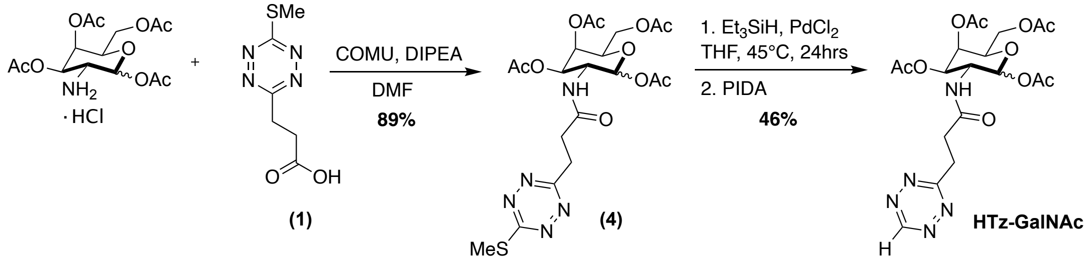


**Supplemental Figure 2. HTz-GalNAc Synthetic Routes.**

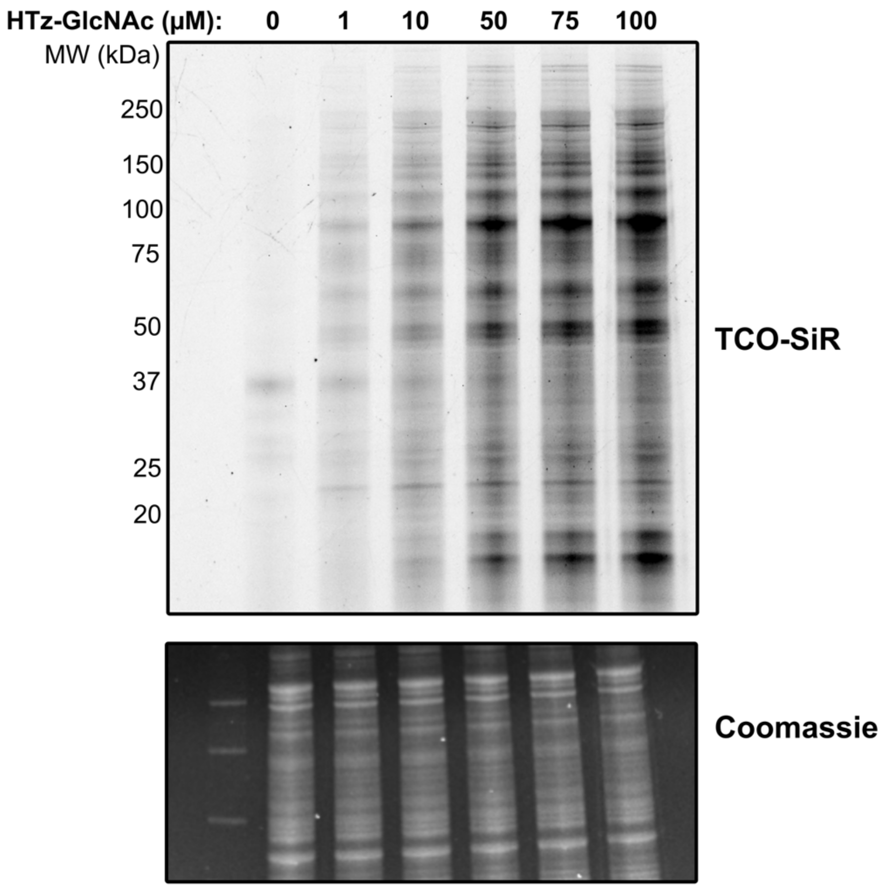


**Supplemental Figure 3. HTz-GlcNAc Exhibits Dose-Dependent Labeling by In-Gel Fluorescence.** BOCTAG SaOS-2 cells were cultured with the indicated concentrations of HTz-GlcNAc followed by TCO-SiR labeling and visualization by in-gel fluorescence. Gels were Coomassie-stained to confirm even protein loading. Gel is representative of three biological replicates.

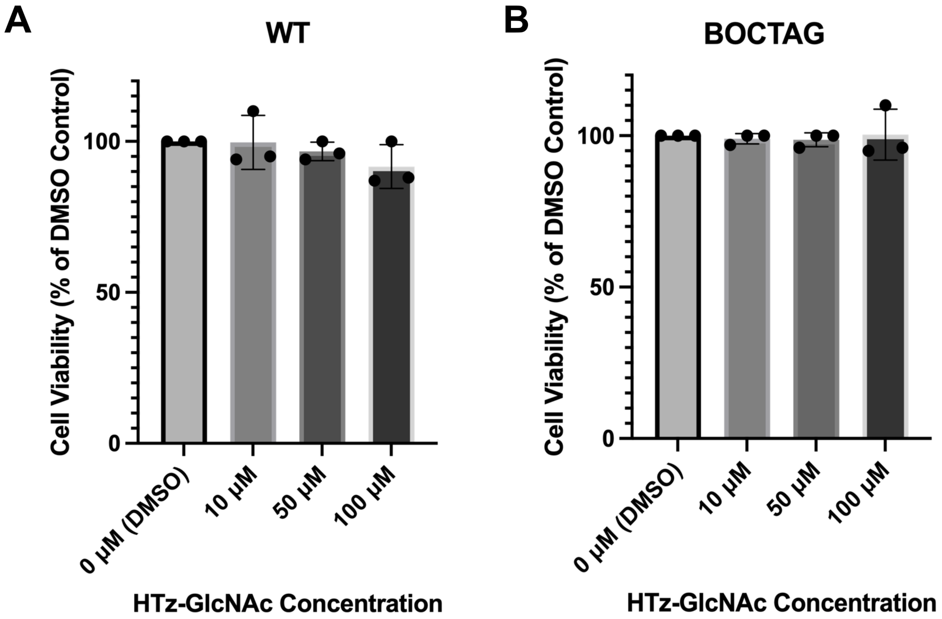


**Supplemental Figure 4. HTz-GlcNAc is Not Toxic to WT and BOCTAG Cells.** Toxicity of varying concentrations of HTz-GlcNAc to **A)** WT SaOS-2 and **B)** BOCTAG SaOS-2 cells were analyzed using CellTiter-Glo. Cell viability is shown as percent live cells as compared to the DMSO-treated control. Statistical significance was analyzed with ordinary one-way ANOVA: for WT, $p$=0.35; for BOCTAG, $p$=0.96.

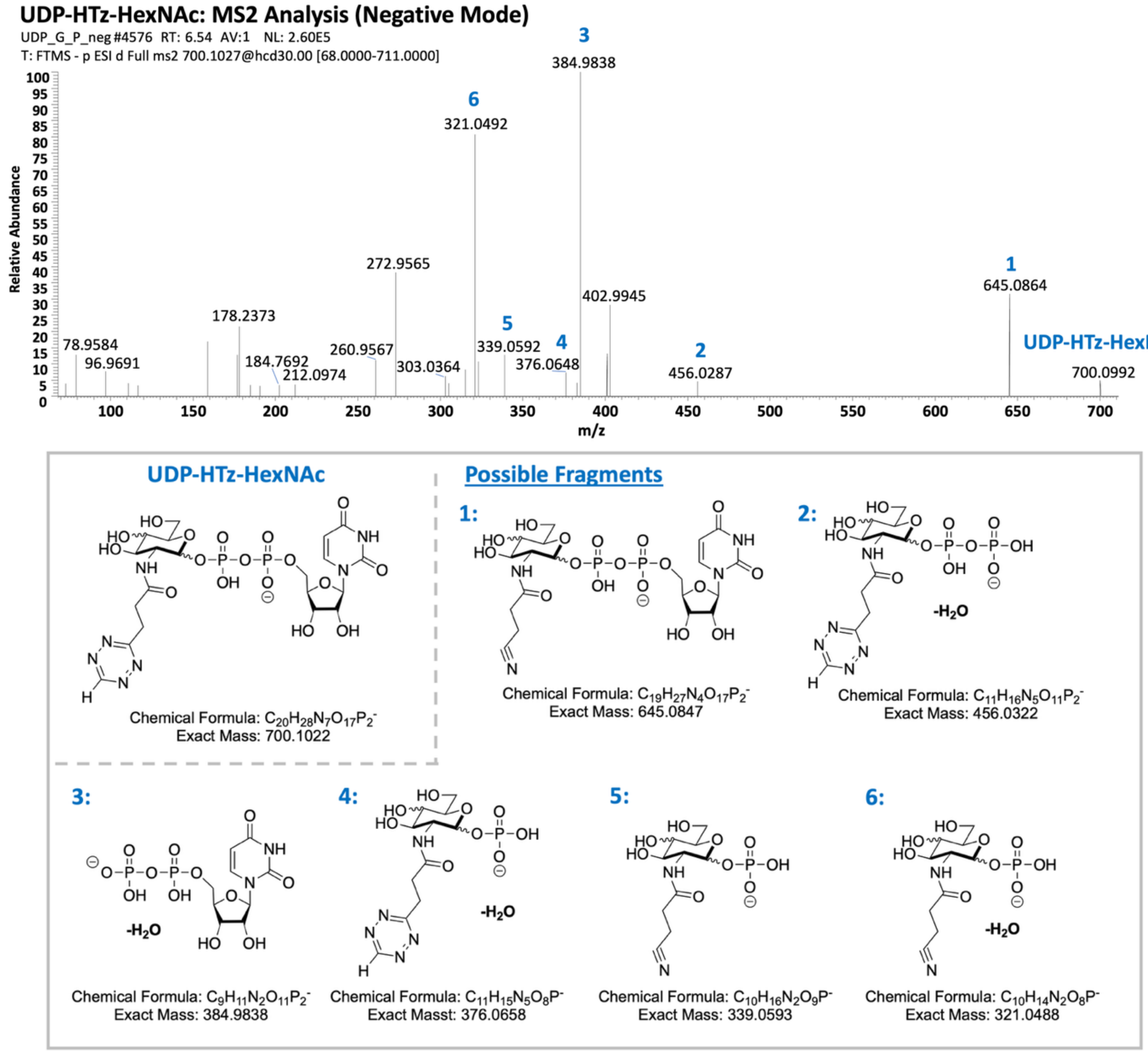


**Supplemental Figure 5. MS/MS detection of UDP-HTz-HexNAc in negative ion mode.** The fragment ion spectrum of the precursor ion at m/z 700.1027 was recorded and possible structures of six selected fragment ions were proposed for comparison with the corresponding UDP-HexNAc fragments.

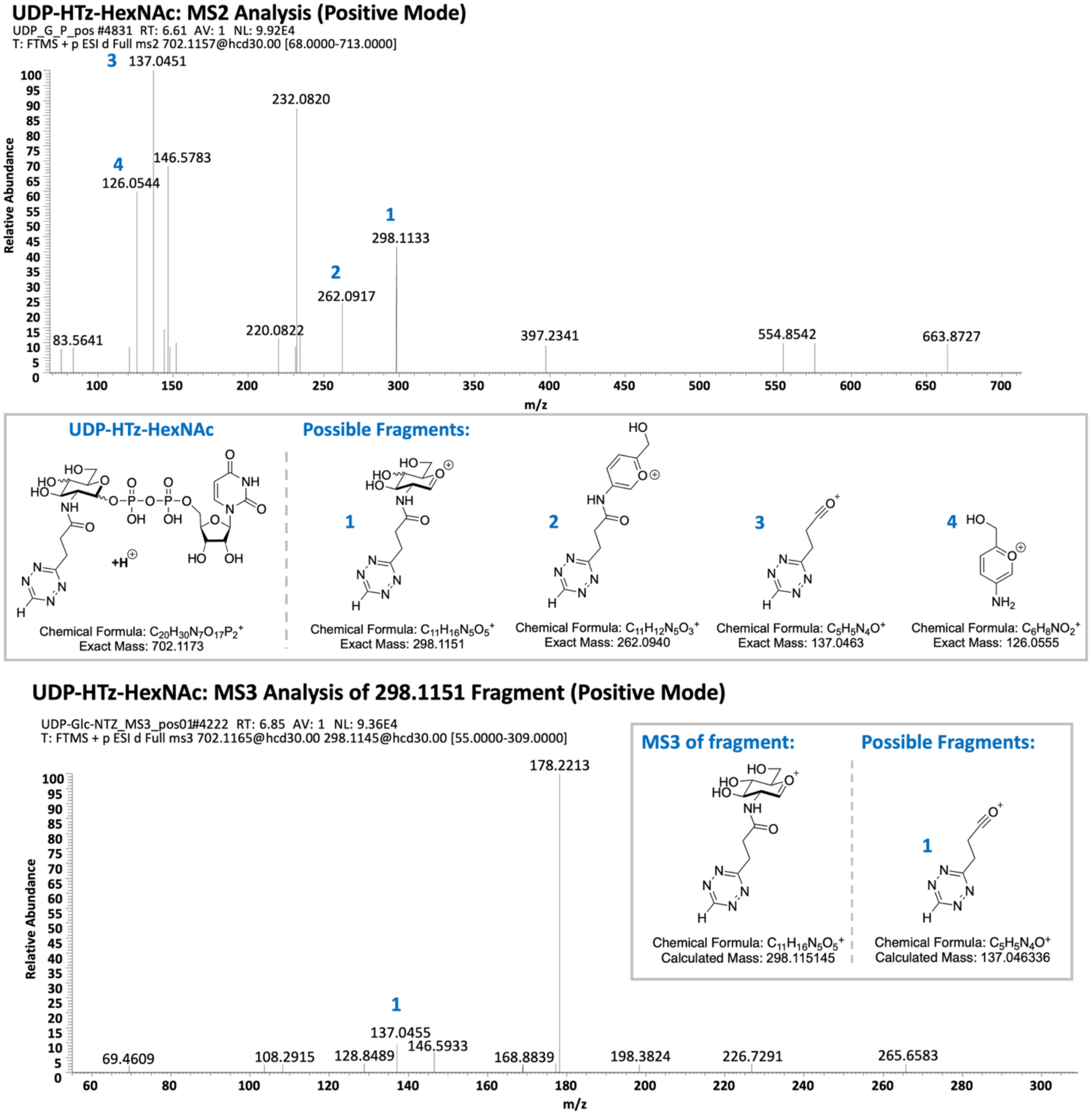


**Supplemental Figure 6. MS/MS and MS/MS/MS detection of UDP-HTz-HexNAc in positive ion mode.** The MS/MS spectrum of the precursor ion at m/z 702.1157 and the $MS^3$ spectrum of the targeted ion at m/z 298.1151 were recorded and possible structures of selected fragment ions were proposed for comparison with the corresponding UDP-HexNAc fragments.

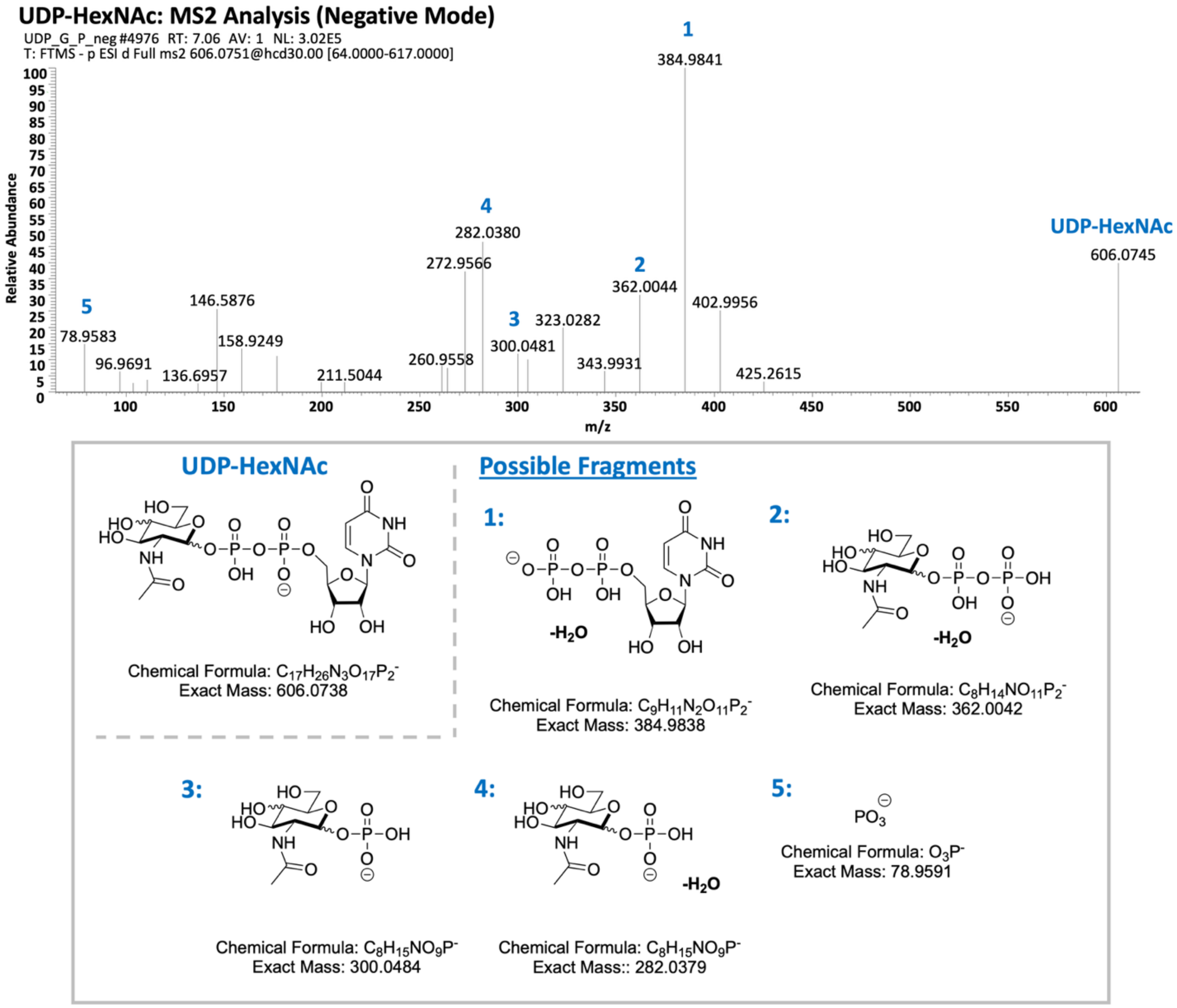


**Supplemental Figure 7. MS/MS detection of UDP-HexNAc in negative ion mode.** The fragment ion spectrum of the precursor ion at m/z 606.0751 was recorded and possible structures of five selected fragment ions were proposed for comparison with the corresponding UDP-HTz-HexNAc fragments.

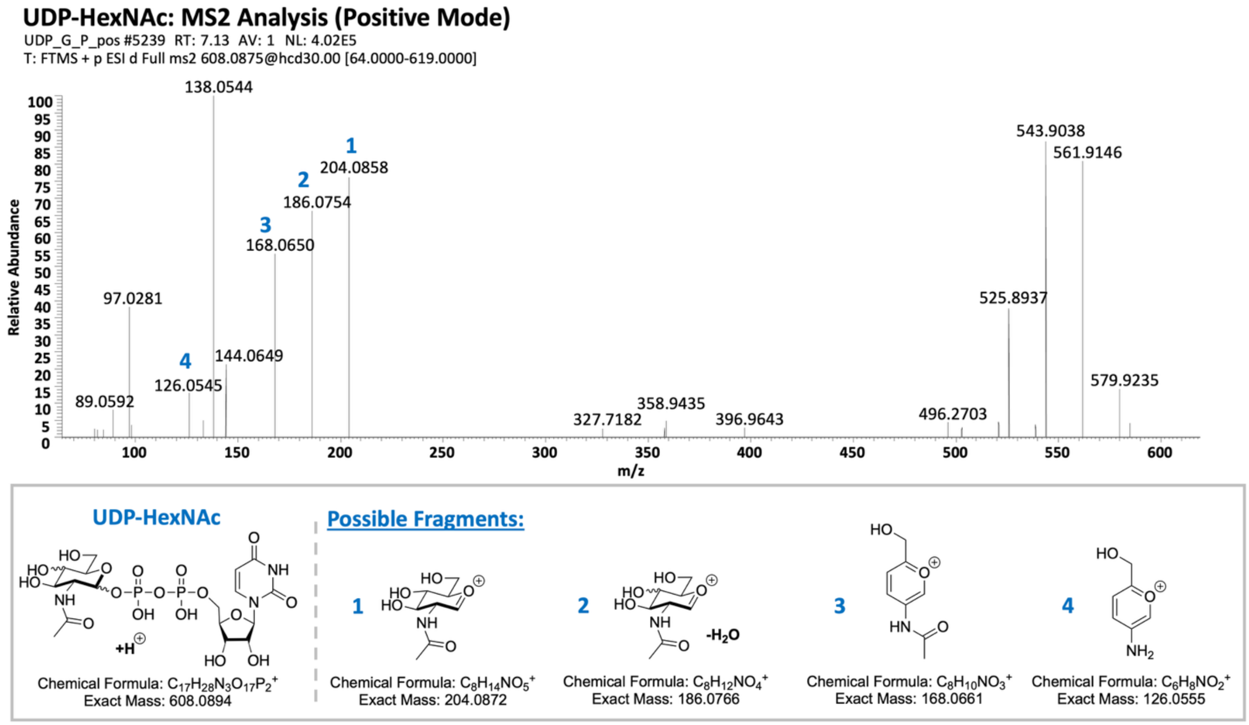


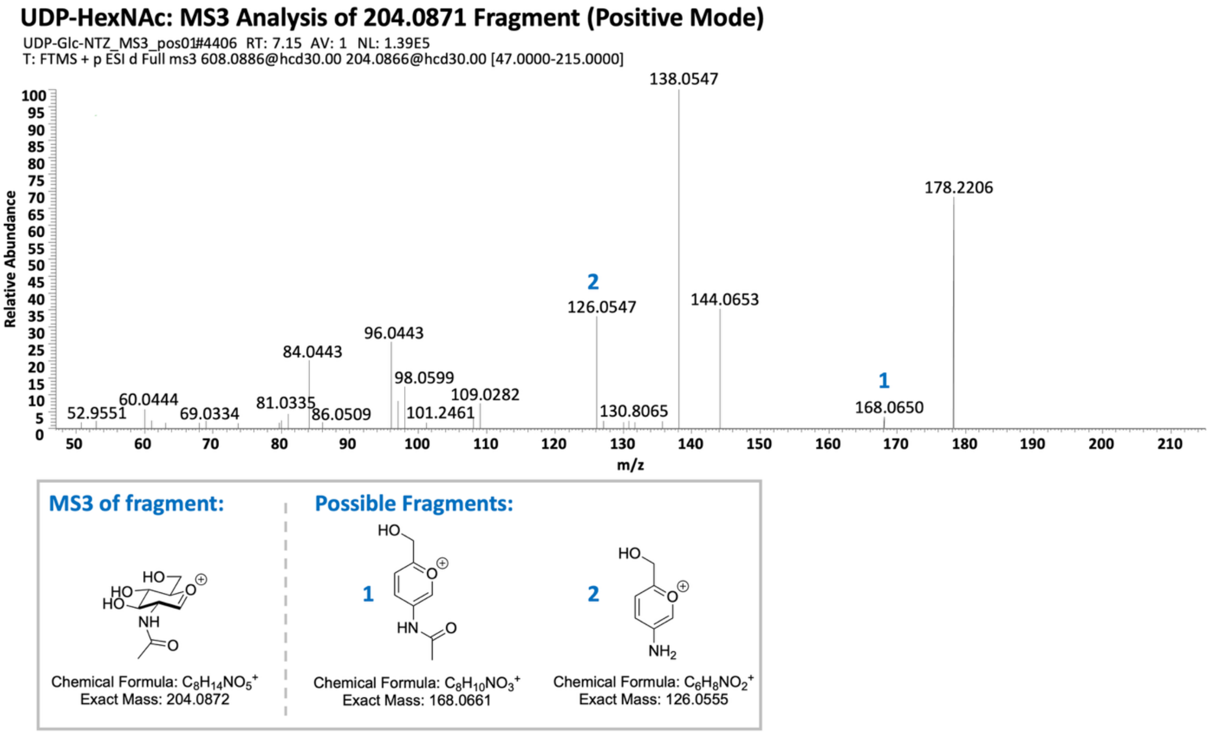


**Supplemental Figure 8. MS/MS and MS/MS/MS detection of UDP-HexNAc in positive ion mode.** The MS/MS spectrum of the precursor ion at m/z 608.0875 and the $MS^3$ spectrum of the targeted ion at m/z 204.0871 were recorded and possible structures of selected fragment ions were proposed for comparison with the corresponding UDP-HTz-HexNAc fragments.

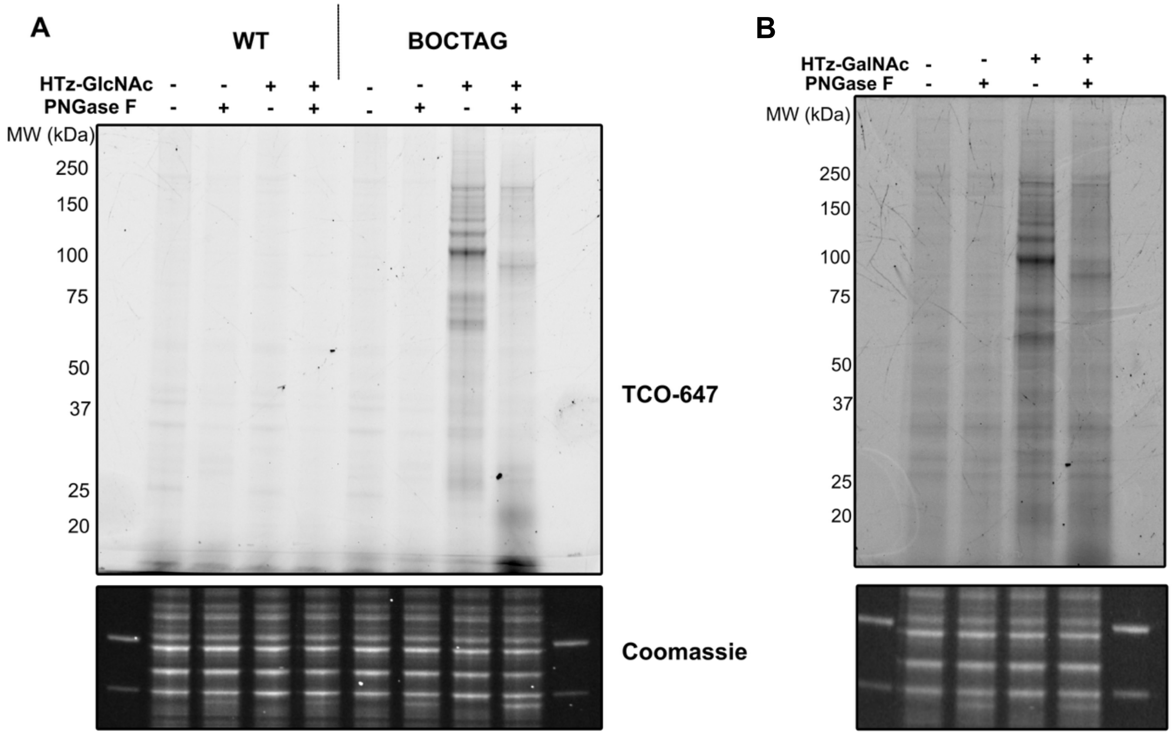


**Supplemental Figure 9. HTz-GlcNAc and HTz-GalNAc label similar species that are sensitive to PNGase F. A)** WT and BOCTAG SaOS-2 cells treated with HTz-GlcNAc and labeled, intact, with TCO-CF647 (1 μM) show BOCTAG-dependent labeling and PNGase F-sensitivity by in-gel fluorescence. **B)** BOCTAG SaOS-2 cells treated with HTz-GalNAc show PNGase F sensitivity. Gels were Coomassie-stained to confirm consistent protein loading. Gels are representative of three biological replicates.

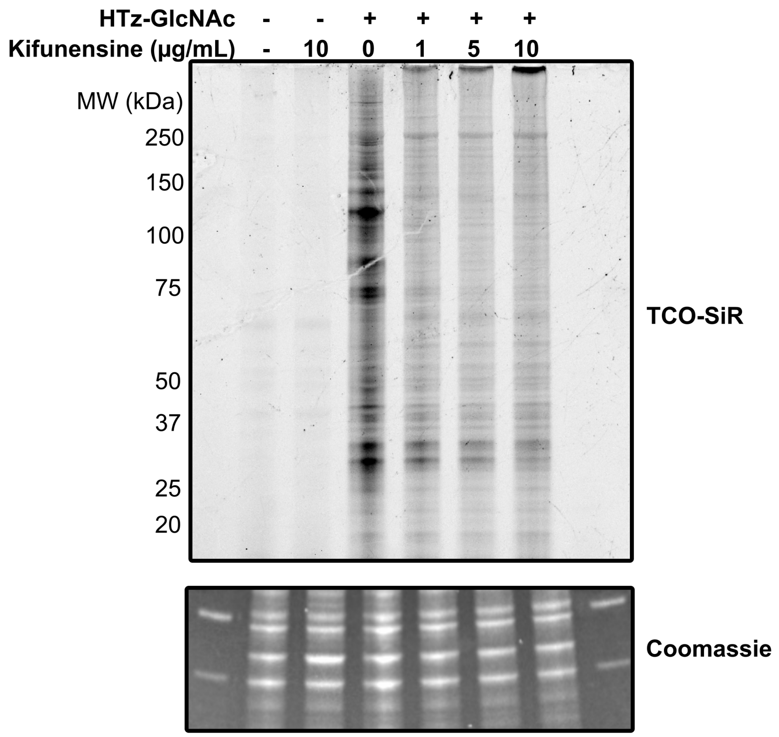


**Supplemental Figure 10. Kifunensine Inhibits HTz-GlcNAc Labeling in SaOS-2 BOCTAG Cells.** BOCTAG SaOS-2 cells were cultured with 50 μM HTz-GlcNAc in the presence of the indicated concentrations of kifunensine. Increasing kifunensine concentrations are associated with reduced HTz-GlcNAc labeling by in-gel fluorescence. Coomassie staining demonstrates consistent protein loading. Gel represents three biological replicates.

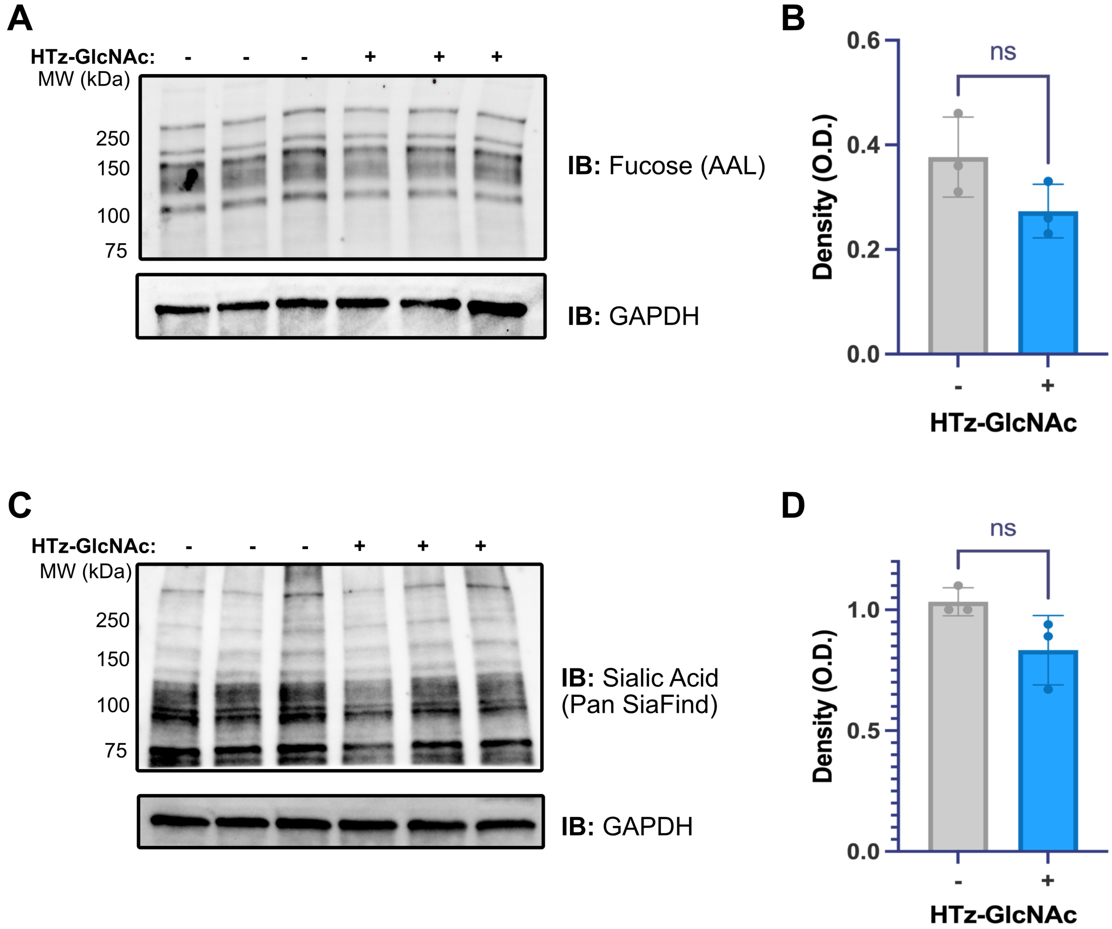


**Supplemental Figure 11. HTz-GlcNAc Impact on Glycan Terminal Modifications. A)** Immunoblot of BOCTAG cells treated with DMSO or HTz-GlcNAc and probed for α-fucose with AAL. GAPDH was used as a loading control. **B)** Quantification of the AAL blot, normalized to GAPDH; statistical significance was analyzed with an unpaired t test: $p$=0.12. **C)** Immunoblot of BOCTAG cells treated with DMSO or HTz-GlcNAc and probed for sialic acid with PanSpecific SiaFind Lectenz. GAPDH was used as a loading control. **D)** Quantification of the SiaFind blot, normalized to GAPDH; statistical significance was analyzed with an unpaired t test: $p$=0.089.

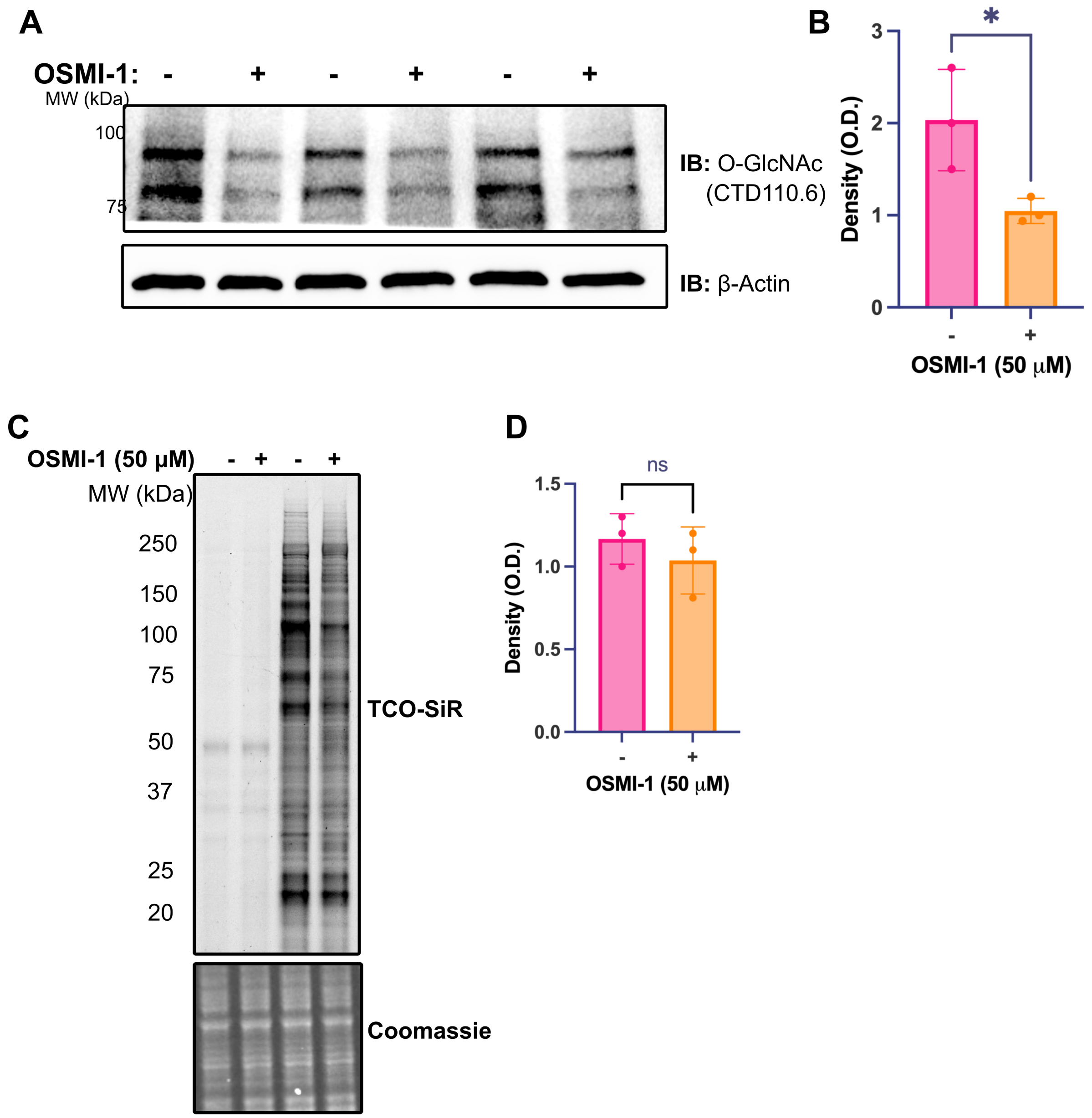


**Supplemental Figure 12. OSMI-1 Treatment of WT and BOCTAG Cells Treated with HTz-GlcNAc or HTz-GalNAc and Verification of OSMI-1 Efficacy. A)** BOCTAG SaOS-2 cells cultured with 50 μM HTz-GlcNAc in combination with DMSO or OSMI-1. Lysates were analyzed by immunoblot, probing with an O-GlcNAc antibody (CTD110.6). **B)** Quantification of CTD110.6 immunoblot, normalized to β-actin. Statistical significance was analyzed by unpaired t test, with * indicating *p* value between 0.05 and 0.01. **C)** BOCTAG SaOS-2 cells were cultured with HTz-GlcNAc in combination with DMSO or OSMI-1, and the impact on TCO-SiR labeling was analyzed by in-gel fluorescence. Coomassie staining confirmed even protein loading. Image is representative of three biological replicates. **D)** Quantification of TCO-SiR fluorescence intensity, normalized to Coomassie stain for total protein. Statistical significance was analyzed by unpaired t test, *p*=0.42.

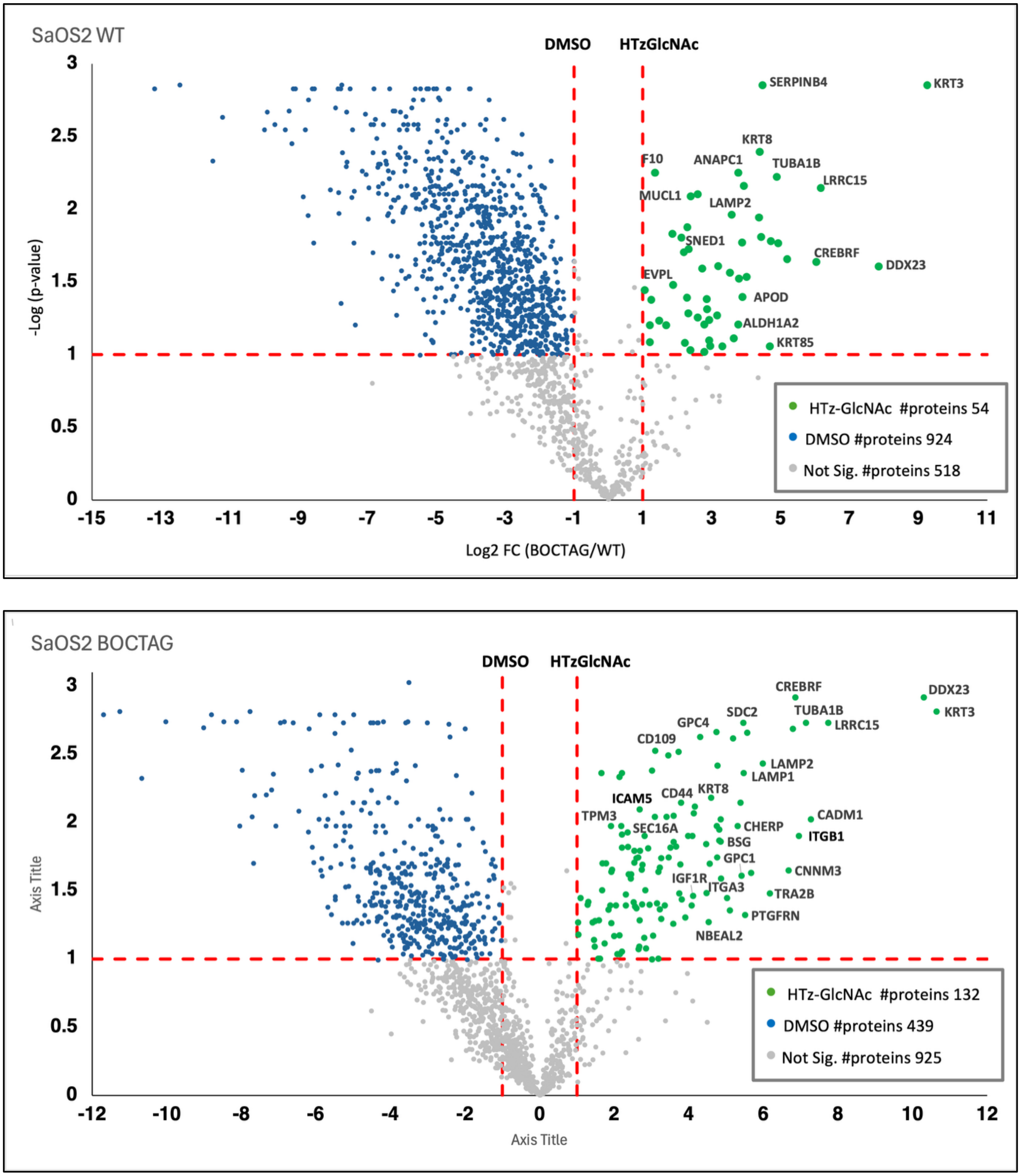


**Supplemental Figure 13. DMSO control proteomics volcano plots.** Wild-type SaOS2 cells (top panel) or BOCTAG SaOS2 cells were cultured with or without 50 □M HTz-GlcNAc, then reacted with TCO-biotin. Biotinylated proteins were enriched on neutravidin resin, then subjected to on-bead trypsin digestion, followed by LC/MS to identify and quantify unmodified peptides. Protein abundance was calculated by label-free quantitation based on precursor ion intensity.

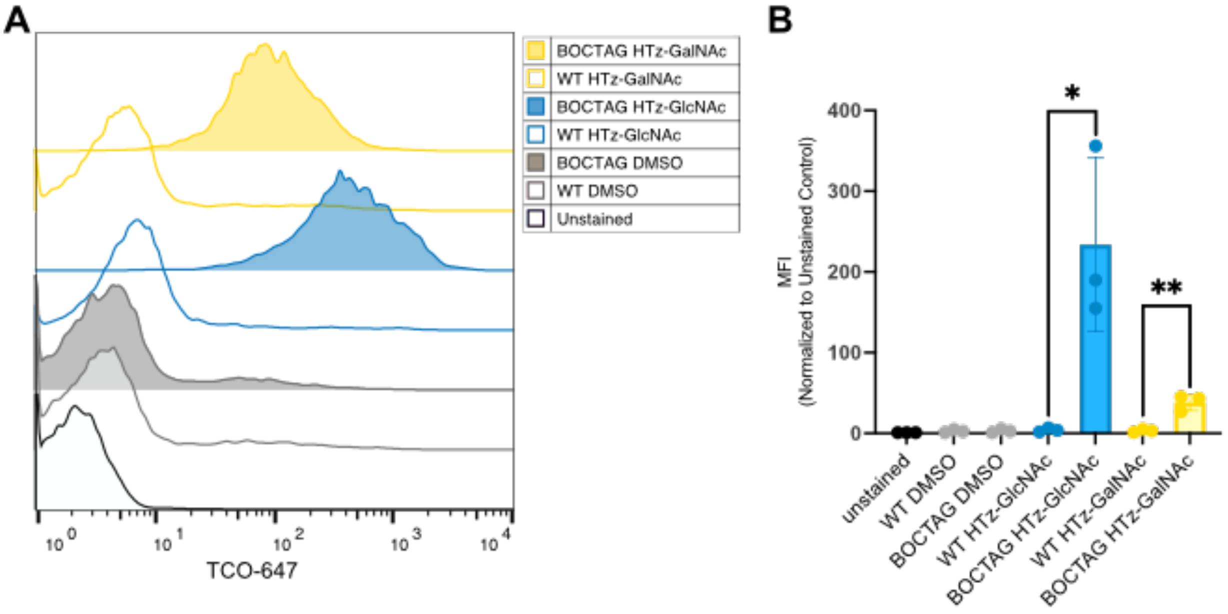


**Supplemental Figure 14. HTz-GlcNAc and HTz-GalNAc Cell Surface Labeling are Dependent on BOCTAG Machinery. A)** Representative flow cytometry data from WT and BOCTAG SaOS-2 cells cultured with DMSO, 50 µM HTz-GlcNAc, or 50 µM HTz-GalNAc and labeled with cell-impermeable TCO-CF647. **B)** Quantification of mean fluorescence intensity (MFI) measured by flow cytometry. Symbols represent biological replicates (n=3), and error bars represent standard deviation. All data were normalized to unstained control. Statistical significance was analyzed by unpaired t test, with * indicating *p* value between 0.05 and 0.01, and ** indicating *p* value between 0.01 and 0.001.

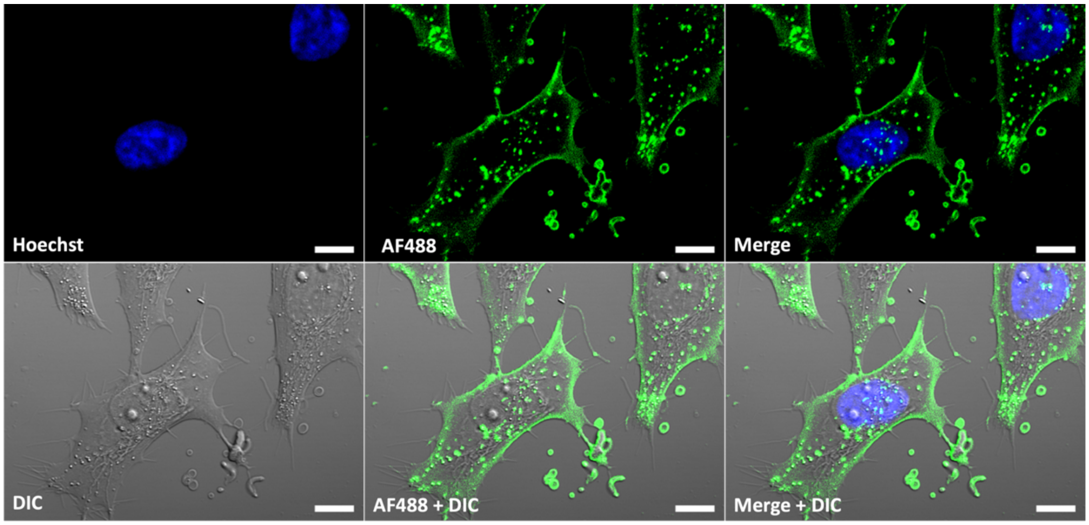


**Supplemental Figure 15. BOCTAG cells cultured with HTz-GlcNAc, then labeled with TCO-AF488.** Day 0-1: Cells treated with 50 µM HTz-GlcNAc for 24 hr; Day 1: Cells were washed, labeled with 1 µM TCO-AF488 (green) and 5 µM Hoechst (blue), washed again, and imaged live at RT. Scale bar is 10 µm.

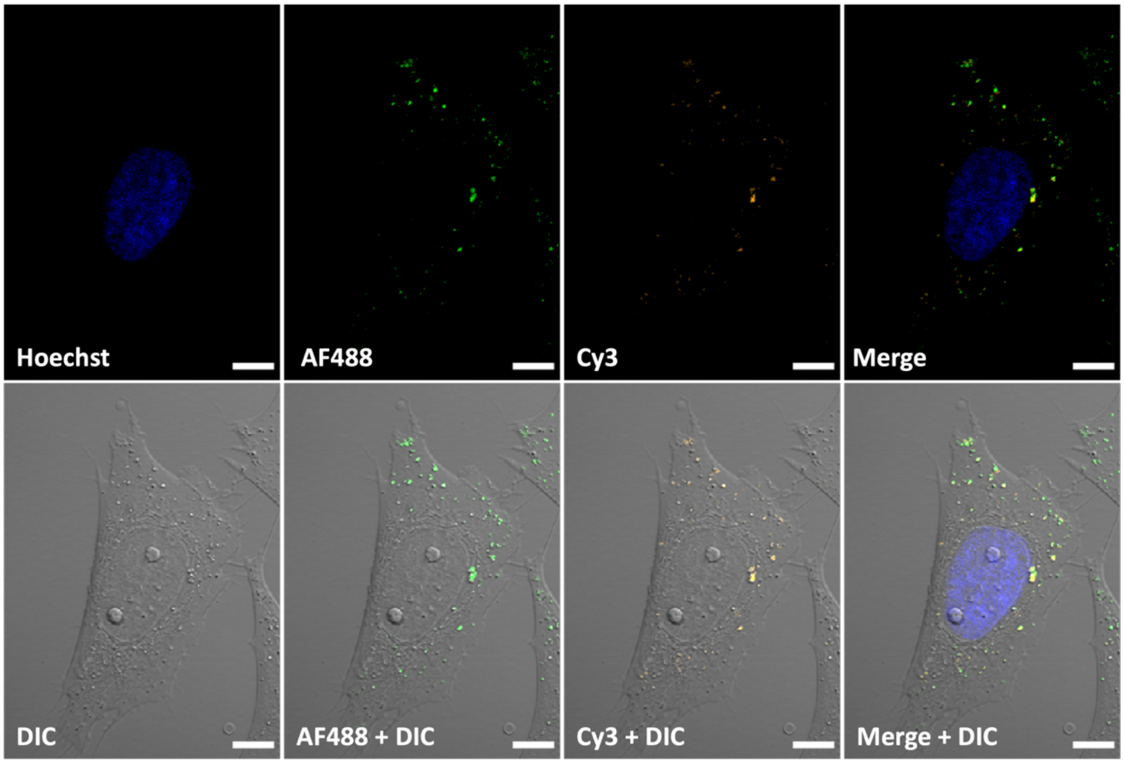


**Supplemental Figure 16. BOCTAG cells cultured once with HTz-GlcNAc (1x), then labeled with TCO-AF488 at 24 hr and Sulfo-Cy3-Peg2-TCO at 48 hr.** Day 0-1: Cells treated with 50 µM HTz-GlcNAc for 24 hr; Day 1: Cells were washed, labeled with 1 µM TCO-AF488 (green), washed again, and quenched with 50 µM methylphenyl-Tz-amine; Day 1-2: Incubated overnight in media; Day 2: Cells were labeled with Sulfo-Cy3-Peg2-TCO (orange) and 5 µM Hoechst (blue), washed, and imaged live at RT. Scale bar is 10 µm.

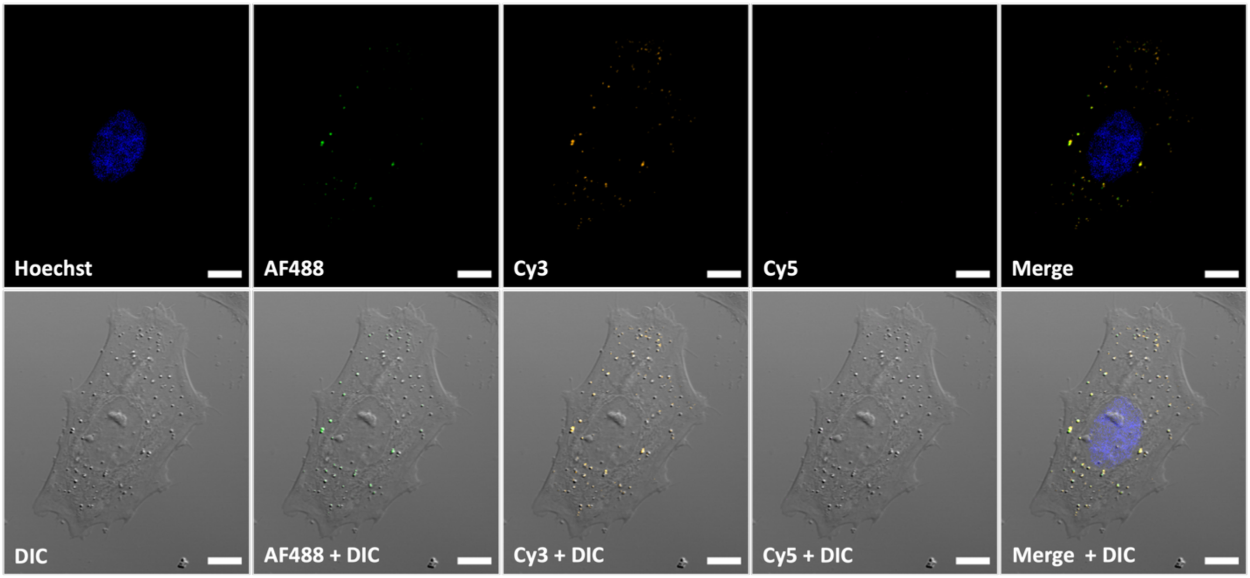


**Supplemental Figure 17. BOCTAG cells cultured once with HTz-GlcNAc (1x), then labeled with TCO-AF488 at 24 hr, Sulfo-Cy3-Peg2-TCO at 48 hr, and Sulfo-Cy5-TCO at 72 hr.** Day 0-1: Cells treated with 50 µM HTz-GlcNAc for 24 hr; Day 1: Cells were washed, labeled with 1 µM TCO-AF488 (green), washed again, and quenched with 50 µM methylphenyl-Tz-amine; Day 1-2: Incubated overnight in media; Day 2: Cells were labeled with Sulfo-Cy3-Peg2-TCO (orange), washed, and quenched with 50 µM methylphenyl-Tz-amine. Day 2-3: Incubated overnight in media; Day 3: Cells were labeled with Sulfo-Cy5-TCO (pink) and 5 µM Hoechst (blue), washed, and imaged live at RT. Scale bar is 10 µm.

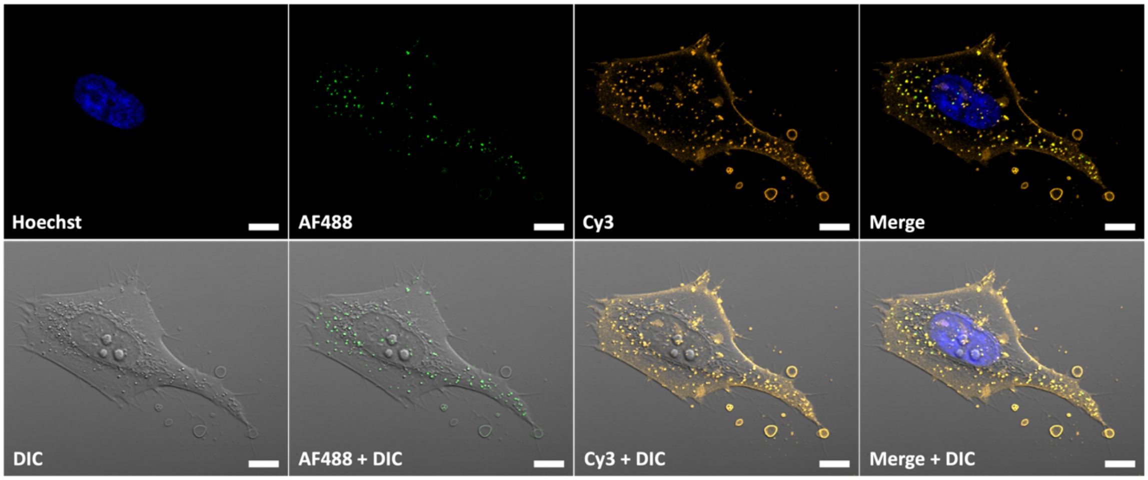


**Supplemental Figure 18. BOCTAG cells cultured twice with HTz-GlcNAc (2x) and labeled with TCO-AF488 at 24 hr and Sulfo-Cy3-Peg2-TCO at 48 hr.** Day 0-1: Cells treated with 50 µM HTz-GlcNAc for 24 hr; Day 1: Cells were washed, labeled with 1 µM TCO-AF488 (green), washed again, and quenched with 50 µM methylphenyl-Tz-amine; Day 1-2: Incubated overnight with 50 µM HTz-GlcNAc; Day 2: Cells were washed, labeled with Sulfo-Cy3-Peg2-TCO (orange) and 5 µM Hoechst (blue), washed, and imaged live at RT. Scale bar is 10 µm.

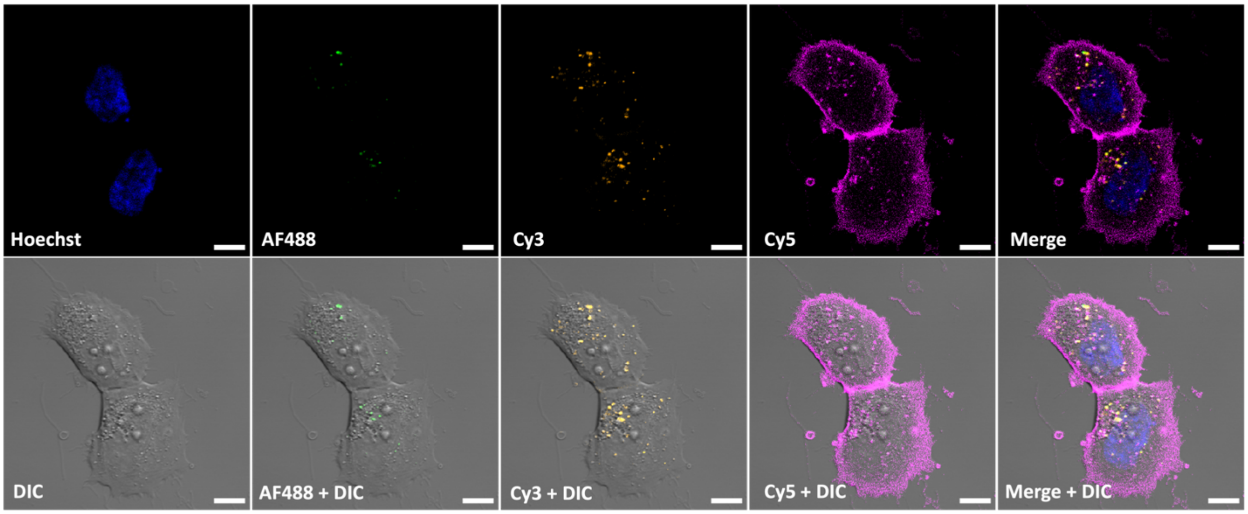


**Supplemental Figure 19. BOCTAG cells cultured three times with HTz-GlcNAc (3x) and then labeled with TCO-AF488 at 24 hr, Sulfo-Cy3-Peg2-TCO at 48 hr, and Sulfo-Cy5-TCO at 72 hr.** Day 0-1: Cells treated with 50 µM HTz-GlcNAc for 24 hr; Day 1: Cells were washed, labeled with 1 µM TCO-AF488 (green), washed again, and quenched with 50 µM methylphenyl-Tz-amine; Day 1-2: Incubated overnight with 50 µM HTz-GlcNAc; Day 2: Cells were washed, labeled with Sulfo-Cy3-Peg2-TCO (orange), washed, and quenched with 50 µM methylphenyl-Tz-amine. Day 2-3: Incubated overnight with 50 µM HTz-GlcNAc; Day 3: Cells were washed, labeled with Sulfo-Cy5-TCO (pink) and 5 µM Hoechst (blue), washed, and imaged live at RT. Scale bar is 10 µm.

### General Information: Materials and Methods

All synthesis reagents were purchased from Sigma Aldrich, Fisher Scientific, Alfa Aesar, TCI Chemicals, J&K Scientific, Chem-Impex International, and Combi-Blocks. NMR solvents were purchased from Cambridge Isotope Laboratories, Inc. Anhydrous dichloromethane and tetrahydrofuran were freshly prepared by an alumina column solvent purification system. Unless otherwise noted, all reactions were performed in flame-dried flasks equipped with rubber septa sealed gas-inlet adaptor, positive pressure of nitrogen, and magnetic stirring. All solvents were anhydrous and transferred via stainless steel needle or cannula with a Luer-lock disposable polypropylene syringe. Reaction progress was monitored by thin layer chromatography (TLC) in which 250 μm glass plates coated with silica gel (Sorbtech Silica HD TLC plates w/ UV 254 or Supelco® TLC Silica Gel 60G $F_{254}$ 25 Glass Plates) were used and visualized with shortwave 254 nm UV light or developed upon heating with $KMnO_4$. Flash chromatography was carried out on silica gel Silicycle Siliaflash P60 silica gel (40-63 μm, 60Å).

SaOS-2 cells were purchased from ATCC (#HTB-35), and BOCTAG cells were generated by transfecting SaOS-2 cells with pSBbi-AGX1-NahK[1], a gift from the Schumann lab, using Lipofectamine 2000 (Thermo Fisher), according to the manufacturers' instructions, with a 10:1 w/w mixture of pSBbi and pCMV(CAT)T7-SB100 plasmid DNA. Cells were selected with Hygromycin B (Thermo Fisher, 80 ug/mL) for 14 days to obtain stable cells. Expression of the BOCTAG system was confirmed by immunoblotting, as previously described in the literature.[1] Cells were cultured in Corning DMEM media with 4.5g/L glucose, sodium pyruvate; without L-glutamine, phenol red (Thermo Fisher; Catalog No. MT17205CV) supplemented with 10 % fetal bovine serum (FBS; Bio-techne), 2 mM L-glutamine, and 2 mM penicillin + streptomycin under sterile conditions at 37 °C with 5 % $CO_2$. Live cell imaging was performed in Opti-MEM™ reduced serum medium without phenol red (Thermo Fisher; Catalog No. 11058021). TCO-AF488, Sulfo-PEG2-Cy3-TCO, and Sulfo-Cy5-TCO were purchased from BroadPharm while aTCO-SiR[1] was synthesized in-house following literature protocol. Lysosome stain (LumiTracker-Lyso Green; Catalog No. B1201), mitochondria stain (LumiTracker Mito Orange CMTMRos; Catalog No. 2252-500ug), endoplasmic reticulum stain (LumiTracker ER Green; Catalog No. 3711-50ug), and Golgi stain (BDP-TMR Ceramide; Catalog No. 3058-250ug) were purchased from Lumiprobe.

All NMR spectra were recorded on Bruker AV 400 MHz, Neo 400 MHz, AV III 600 MHz, and Neo 600 MHz spectrometers. Chemical shifts are reported in parts per million (ppm) and referenced to their residual non-deuterated solvents peaks: $CDCl_3$ ($^1$H: 7.26 ppm, $^{13}$C: 77.16 ppm), methanol-d4 ($^1$H: 3.31 ppm, $^{13}$C: 49.00 ppm), DMSO-d6 ($^1$H: 2.50 ppm, $^{13}$C: 39.52 ppm), and deuterium oxide ($^1$H: 4.79 ppm). Coupling constants (J) are reported to the nearest 0.1 MHz. Multiplicities are reported as follows singlet (s), doublet (d), triplet (t), quartet (q), multiplet (m), 'broad' (br), coupling constant in Hz, integration, and assignment for carbohydrates was based on two-dimensional COSY, HSQC, and HMBC experiments. $^{13}$C NMR resonances are proton decoupled and an APT pulse sequence was used to determine the type of carbon as noted:

methylene and quaternary ($CH_2$ and C) carbons appear 'up' and methyl and methine ($CH_3$ and CH) carbons appear 'down'. $^{19}F$ NMR data were collected with proton decoupling. High-resolution mass spectra (HR-MS), ESI mode, were obtained on a Thermo Q-Exactive Orbitrap or a Waters GCT Premier in the Mass Spectroscopy Facility at the Department of Chemistry and Biochemistry, University of Delaware. Images were taken on Zeiss LSM880 Confocal Microscope (Plan-Apopchromat 40 water objective) or Andor Dragonfly 600 Spinning Disk Confocal Microscope (Plan-Apochromat 63x/1.4 oil objective).

**Synthetic Procedures:**

#### Methyl 3-(6-(methylthio)-1,2,4,5-tetrazin-3-yl) propanoic acid (1)

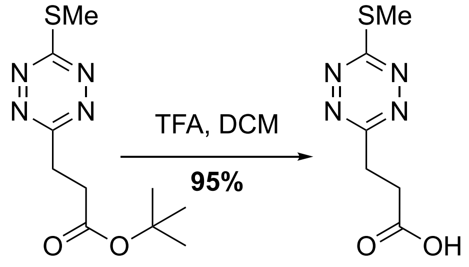


**(1)**

A round bottom flask was charged with a stir bar and *tert*-butyl 3-(6-(methylthio)-1,2,4,5-tetrazin-3-yl)propanoate[2] (4.34 g, 16.93 mmol, 1 eq), which was then dissolved in 17 mL of DCM (1 M). While stirring trifluoracetic acid (26 mL, 338.6 mmol, 20 eq) was added, the flask was capped and allowed to stir at room temperature until all the starting material was consumed. Reaction was monitored by TLC (100 % DCM). After the starting material was consumed the reaction was concentrated via rotary evaporation. A bright pink solid (3.21 g, 16.03 mmol, 95 %) was isolated using a silica plug (0 to 2 % methanol in DCM).

$^1H$ NMR (400 MHz, $CDCl_3$) $\delta$ 3.58 (t, 2H, $J$ = 7.0 Hz), 3.08 (t, 2H, $J$ = 7.1 Hz), 2.73 (s, 3H)

$^{13}C$ NMR (101 MHz, $CDCl_3$) $\delta$ 177.2 (C), 175.7 (C), 165.8 (C), 30.3 ($CH_2$), 28.7 ($CH_2$), 13.2 ($CH_3$)

HRMS (ESI+) [ M + H]$^+$ m/z calculated for $[C_6H_9N_4O_2S]^+$ 201.044623, found 201.04505.

**Pentafluorophenyl 3-(6-(methylthio)-1,2,4,5-tetrazin-3-yl) propanoate (2)**

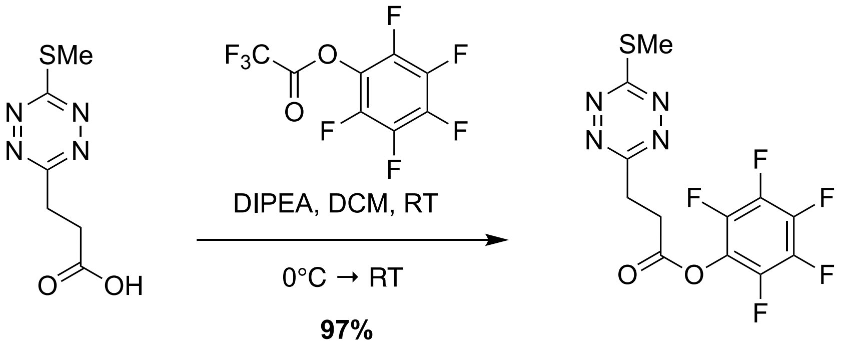


(1) (2)

A flame-dried round bottom flask was charged with **1** (315.5 mg, 1.58 mmol, 1.0 eq), which was suspended in anhydrous DCM (15.8 mL, 0.1 M) under $N_2$. Diisopropylethylamine (823 µL, 4.73 mmol, 3.0 eq) was added and the suspension was stirred at RT for 30 min. The mixture was cooled to 0 °C and pentafluorophenyltrifluoroacetate (298 µL, 1.73 mmol, 1.1 eq) was added dropwise over 5 min, the reaction was allowed to stir at 0 °C for 1 h, followed by 2 h at RT. The reaction solution was directly purified via a silica plug (100 % DCM) to yield a dark red solid (557.6 mg, 97 %)

$^{1}$H NMR (600 MHz, $CDCl_3$) $\delta$ 3.7 (t, 2H, $J$ = 7.0 Hz), 3.4 (t, 2H, $J$ = 7.2 Hz), 2.7 (s, 3H)

$^{13}$C NMR (151 MHz, $CDCl_3$) $\delta$ 176.5 (C), 168.1 (C), 165.2 (C), 141.0 (dm, CF, $J$ = 235.9 Hz), 139.5 (dm, CF, $J$ = 227.6 Hz), 137.8 (dm, CF, $J$ = 258.9 Hz), 124.8 (tm, C, $J$ = 15.3 Hz), 29.9 ($CH_2$), 28.9 ($CH_2$), 13.3 ($CH_3$)

$^{19}$F NMR (376 MHz, $CDCl_3$) $\delta$ -152.46 to -152.50 (m, 2F), -157.70 (t, 1F, $J$ = 22.2 Hz), -162.1 to -162.2 (m, 2F)

HRMS (ESI+) $[M + H]^+$ m/z calculated for $[C_{12}H_8F_5N_4O_2S]^+$ 367.028813, found 367.02826.

**SMeTz-GlcNAc (3)**

*Route 1:*

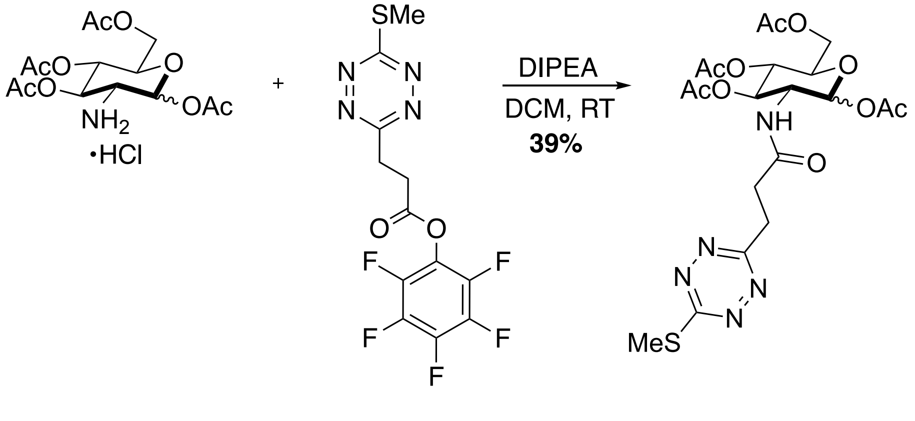


(2) (3)

A flame-dried round bottom flask was charged with 1,3,4,6-tetraacetyl-β-d-glucosamine hydrochloride (354.4 mg, 923.4 µmol, 1.0 eq), which was suspended in DCM (12.25 mL, 0.08 M). Diisopropylethylamine (483 µL, 2.770 mmol, 3.0 eq) was added to the suspension and stirred at RT for 10 min. Compound **2** (500.1 mg, 1.365 mmol, 1.5 eq) was dissolved in DCM (5.25 mL, 0.3 M) and added dropwise to the reaction over 10 minutes. The reaction mixture was then allowed to stir at RT for 4hr. A pink oil was isolated via flash chromatography (0 → 40 % Ethyl Acetate in DCM) the oil was precipitated with a mixture of DCM : pentane and dried via rotatory evaporation/ high vac to yield a bright pink solid (190.7 mg, 39 %).

*Route 2:*

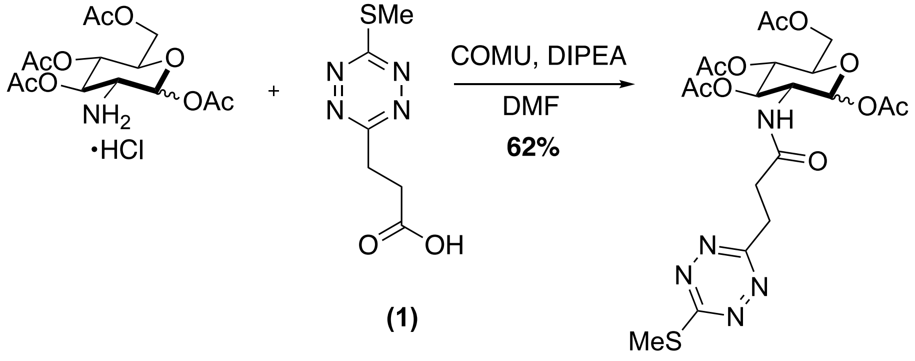


(1)

(3)

A flame-dried round bottom flask was charged with 1,3,4,6-tetraacetyl-β-d-glucosamine hydrochloride (434.8 mg, 2.016 mmol, 2.0 eq), **1** (201.0 mg, 1.004 mmol, 1.0 eq), COMU (867.2 mg, 2.025 mmol, 2.0 eq), DIPEA (702 µL, 4.033 mmol, 4.0 eq), and anhydrous DMF (5.0 mL, 0.2 M).[3] The reaction was allowed to stir overnight at RT under $N_2$. The crude mixture was diluted with ethyl acetate and washed with 1.0 M aqueous HCl (2x) , sat. aqueous sodium bicarbonate solution (2x), and brine (1x). The organic layer was dried over sodium sulfate, filtered, and concentrated via rotary evaporation. A pink oil was isolated via flash chromatography (0 → 40 % Ethyl Acetate in DCM), the oil was precipitated with a mixture of DCM : pentane and dried via rotatory evaporation/ high vac to yield a bright pink solid (327.9 mg, 62 %)

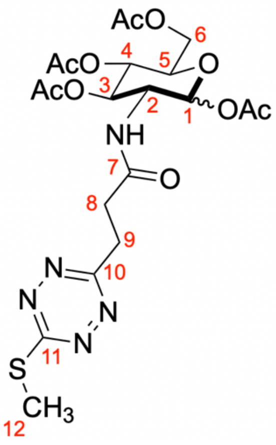


$^{1}$H NMR (600 MHz, Acetone-D$_6$) $\delta$ (Anomers >9$\alpha$ : 1β) 7.16 (d, $J$ = 9.6 Hz, 1H, amide NH), 6.16 (d, $J$ = 3.8 Hz, 1H, $\alpha$H$_1$), 5.28 (dd, $J$ = 10.9, 9.7 Hz, 1H, $\alpha$H$_3$), 5.13 (t, $J$ = 9.9 Hz, 1H, $\alpha$H$_5$), 4.44 (ddd, $J$ = 11.1, 9.2, 3.8 Hz, 1H, $\alpha$H$_2$), 4.23 (dd, $J$ = 12.1, 4.0 Hz, 1H, $\alpha$H$_{6a}$), 4.22-4.19 (m, 1H, $\alpha$H$_5$), 4.06 (dd, $J$ = 12.1, 2.0 Hz, 1H, $\alpha$H$_{6b}$), 3.68- 3.58 (m, 2H, $\alpha$H$_{9a,b}$), 3.2 (t, $J$ = 6.7 Hz, 2H, $\alpha$H$_{8a,b}$), 2.75 (s, 3H, $\alpha$H$_{12}$), 2.01 (s, 3H, acetyl), 1.99 (s, 3H, acetyl), 1.96 (s, 3H, acetyl), 1.86 (s, 3H, acetyl)

$^{13}$C NMR (151 MHz, Acetone-D$_6$) $\delta$ 176.3 (C), 171.4 (C), 170.9 (C), 170.8 (C), 170.6 (C), 169.9 (C), 167.4 (C), 91.8 (CH, $\alpha$C$_1$), 71.3 (CH, $\alpha$C$_3$), 70.8 (CH, $\alpha$C$_5$), 69.2 (CH, $\alpha$C$_4$), 62.4 (CH$_2$, $\alpha$C$_6$), 51.6 (CH, $\alpha$C$_2$), 31.5 (CH$_2$, $\alpha$C$_8$), 22.7 (CH$_3$, acetyl), 20.6 (3x CH$_3$, 3x acetyl), 13.4 (CH$_3$, $\alpha$C$_{12}$) * $\alpha$C$_9$ obscured by solvent peak

HRMS (ESI+) [ M + H]$^+$ m/z calculated for $[C_{20}H_{28}N_5O_{10}S]^+$ 530.155692, found 530.15582.

**HTz-GlcNAc**

*Route 1:*

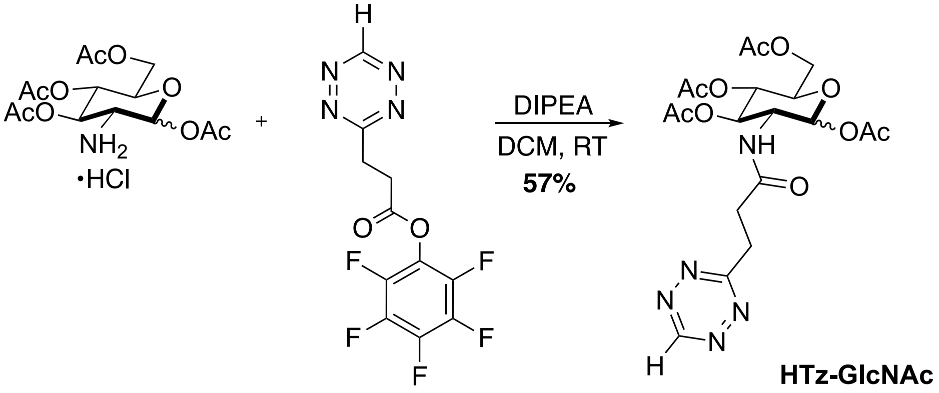


A flame-dried round bottom flask was charged with 1,3,4,6-tetraacetyl-β-d-glucosamine hydrochloride (21.2 mg, 55.24 μmol, 1.0 eq), which was suspended in DCM (750 μL, 0.07 M) and cooled to 0 °C. Diisopropylethylamine (28.8 μL, 165.7 μmol, 3.0 eq) was added to the suspension and stirred at 0 °C for 10 min. HTz-Pentafluorophenyl ester (26.5 mg, 82.77 μmol, 1.5 eq), prepared as previously described[4], was dissolved in DCM (300 μL, 0.3 M) and added dropwise to the reaction over 5 minutes. The reaction mixture was then allowed to stir at RT for 3 hr. A pink oil was isolated via flash chromatography (0 → 50 % Ethyl Acetate in DCM), the oil was precipitated with a mixture of DCM : pentane and dried via rotatory evaporation/ high vac to yield a bright pink solid (15.3 mg, 57 %).

*Route 2:*

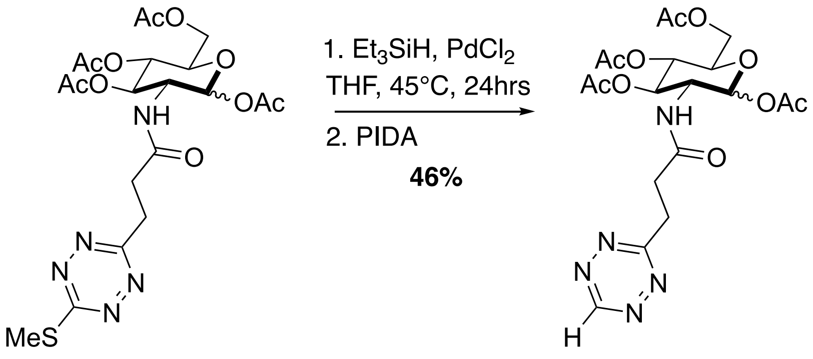


**(3)** **HTz-GlcNAc**

A flame-dried Schlenk tube was charged with **3** (122.0 mg, 230.4 µmol, 1.0 eq) and palladium II chloride (8.05 mg, 45.4 µmol, 20 mol %, 0.2 eq) and evacuated and purged three times with $N_2$. Then triethylsilane (110.3 µL, 690.6 µmol, 3.0 eq) was added to anhydrous THF (3.4 mL, 0.07 M), the mixture was added to the Schlenk tube under $N_2$, and the suspension was allowed to stir at 45 °C for 20-24 hr. The reaction was cooled to RT and PIDA (223 mg, 691.2 µmol, 3.0 eq) was added and allowed to stir at RT for 1 hr. The suspension was then filtered through celite with ethyl acetate. The filtrate was washed with saturate sodium bicarbonate, water, and brine. The organic layer was dried over sodium sulfate, filtered, and concentrated via rotary evaporation. A pink oil was isolated via flash chromatography (0 → 30 % Acetone in Chloroform), the oil was precipitated with a mixture of DCM : pentane and dried via rotatory evaporation/ high vac to yield a bright pink solid (51.2 mg, 46 %)

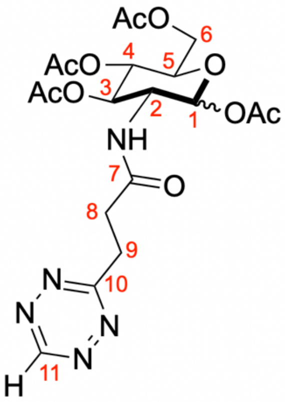


$^1$H NMR (600 MHz, Acetone-D$_6$) $\delta$ (Anomers >9$\alpha$ : 1β) 10.35 (s, 1H, $\alpha$H$_{11}$), 7.40 (d, $J$ = 9.3 Hz, 1H, amide NH), 6.09 (d, $J$ = 3.60 Hz, 1H, $\alpha$H$_1$), 5.29 (dd, $J$ = 9.6 Hz, 1H, $\alpha$H$_3$), 5.10 (t, $J$ = 10.1 Hz, 1H, $\alpha$H$_4$), 4.40 (ddd, $J$ = 11.1, 9.1, 3.6 Hz, 1H, $\alpha$H$_2$), 4.21 (dd, $J$ = 12.4, 4.2 Hz, 1H, $\alpha$H$_{6a}$), 4.16 (ddd, $J$ = 10.1, 4.2, 2.2 Hz, 1H, $\alpha$H$_5$), 4.03 (dd, $J$ = 12.4, 2.2 Hz, 1H, $\alpha$H$_{6b}$), 3.61- 3.51 (m, 2H, $\alpha$H$_{9a,b}$), 2.96- 2.88 (m, 2H, $\alpha$H$_{8a,b}$), 2.17 (s, 3H, acetyl), 2.00 (s, 3H, acetyl), 1.99 (s, 3H, acetyl), 1.98 (s, 3H, acetyl)

$^{13}$C NMR (151 MHz, Acetone-D$_6$) $\delta$ 172.9 (C), 172.1 (C), 171.0 (C), 170.7 (C), 169.9 (C), 169.7 (C), 159.0 (C), 91.1 (CH, $\alpha$C$_1$), 71.2 (CH, $\alpha$C$_3$), 70.6 (CH, $\alpha$C$_5$), 69.1 (CH, $\alpha$C$_4$), 62.0 (CH$_2$, $\alpha$C$_6$),

51.4 (CH, $\alpha C_2$), 32.5 ($CH_2$, $\alpha C_8$), 31.1 ($CH_2$, $\alpha C_9$), 20.9 ($CH_3$, acetyl), 20.7 ($CH_3$, acetyl), 20.6 (2x $CH_3$, 2x acetyl)

HRMS (ESI+) [ M + H]$^+$ m/z calculated for $[C_{19}H_{26}N_5O_{10}]^+$ 484.167969, 484.16785.

**SMeTz-GalNAc (4)**

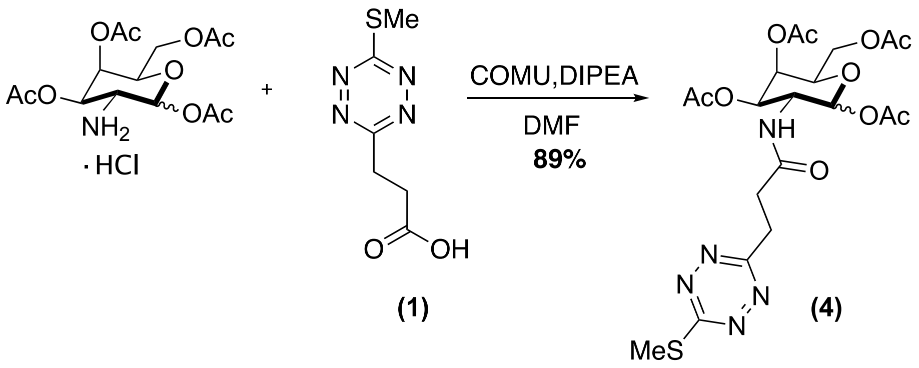


**(1)** **(4)**

A flame-dried round bottom flask was charged with 1,3,4,6-tetraacetyl-d-galactosamine hydrochloride[5] (1.077 g, 2.81 mmol, 1.1 eq), **1** (500 mg, 2.50 mmol, 1.0 eq), COMU (2.14 g, 4.99 mmol, 2.0 eq), DIPEA ( 1.74 mL, 9.99 mmol, 4.0 eq), and the flask was carefully evacuated and purged with $N_2$. Then anhydrous DMF (12.5 mL, 0.2 M) was added, and reaction was allowed to stir overnight at RT under $N_2$. The crude mixture was diluted with ethyl acetate and washed with 1.0 M aqueous HCl (2x), sat. aqueous sodium bicarbonate solution (2x), and brine (2x). The organic layer was dried over sodium sulfate, filtered, and concentrated via rotary evaporation. A bright pink solid (1.185g, 90 %) was isolated via flash chromatography (0 → 30 % Ethyl Acetate in DCM).

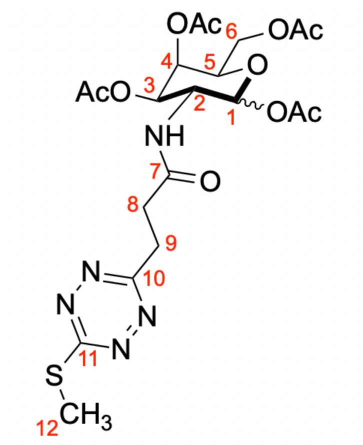


$^1$H NMR (600 MHz, $CDCl_3$) $\delta$ (Anomers >9$\alpha$ : 1β) 5.79 (d, $J$ = 9.5 Hz, 1H, amide NH), 5.71 (d, $J$ = 8.8 Hz, 1H, $\alpha H_1$), 5.36 (d, $J$ = 3.4 Hz, 1H, $\alpha H_4$), 5.11 (dd, $J$ = 11.3, 3.4 Hz, 1H, $\alpha H_3$), 4.40 (dt, $J$ = 11.4, 9.1 Hz, 1H, $\alpha H_2$), 4.15 (dd, $J$ = 11.4, 6.5 Hz, 1H, $\alpha H_{6a}$), 4.10 (dd, $J$ = 11.3, 6.5 Hz, 1H, $\alpha H_{6b}$), 4.03 (t, $J$ = 6.5 Hz, 1H, $\alpha H_5$), 3.56 (td, $J$ = 6.6, 2.9 Hz, 2H, $\alpha H_9$), 2.82 (t, $J$ = 6.8 Hz, 2H, $\alpha H_8$), 2.71 (s, 3H, $\alpha H_{12}$), 2.15 (s, 3H, acetyl), 2.11 (s, 3H, acetyl), 2.03 (s, 3H, acetyl), 2.01(s, 3H, acetyl).

$^{13}$C NMR (151 MHz, $CDCl_3$) $\delta$ 170.6 (C), 171.6 (C), 171.1 (C), 170.8 (C), 170.5 (C), 170.0 (C), 166.6 (C), 93.2 (CH, $\alpha C_1$), 72.6 (CH, $\alpha C_5$), 70.7 (CH, $\alpha C_3$), 66.7 (CH, $\alpha C_4$), 61.7 ($CH_2$, $\alpha C_6$), 49.8 (CH, $\alpha C_2$), 32.6 ($CH_2$, $\alpha C_8$), 29.6 ($CH_2$, $\alpha C_9$), 21.2 ($CH_3$, 2x Acetyl), 21.0 ($CH_3$, 2x Acetyl), 13.6 ($CH_3$, $\alpha C_{12}$).

HRMS (ESI+) [ M + H]$^+$ m/z calculated for $[C_{20}H_{28}N_5O_{10}S]^+$ 530.155692, found 530.15613.

**HTz-GalNAc**

*Route 1:*

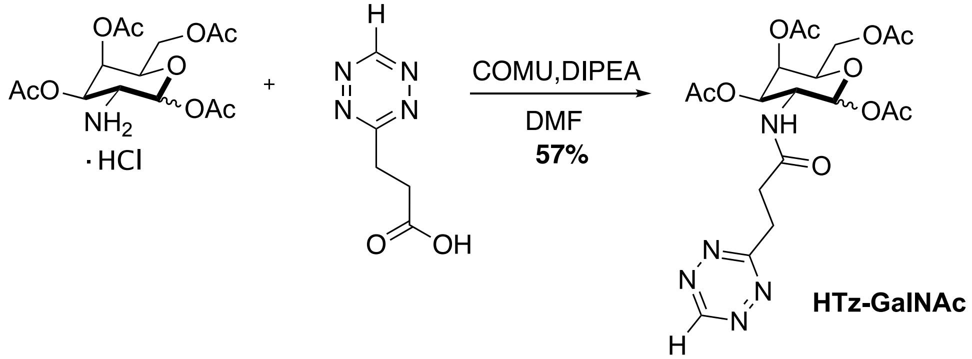


A dry 7 mL vial was charged with 1,3,4,6-tetraacetyl-d-galactosamine hydrochloride[5] (95.1 mg, 247.8 µmol, 1.15 eq), COMU (194 mg, 453 µmol, 2.1 eq), and HTz-acid (33.2 mg, 215.4 µmol, 1.0 eq), which was prepared as previously described.[4] Anhydrous DMF (1.08 mL, 0.2 M) and DIPEA ( 151 µL, 862 µmol, 4.0 eq) were added to the vial, the vial was sparged with $N_2$, and the reaction was allowed to stir at RT overnight. The crude mixture was diluted with ethyl acetate and washed with 1.0 M aqueous HCl (2x), sat. aqueous sodium bicarbonate solution (2x), and brine (2x). The organic layer was dried over sodium sulfate, filtered, and concentrated via rotary evaporation. A pink oil was isolated via flash chromatography (0 → 20 % Acetone in DCM), the oil was precipitated with a mixture of DCM : pentane and dried via rotatory evaporation/ high vac to yield a bright pink solid (58.9 mg, 57 %).

*Route 2:*

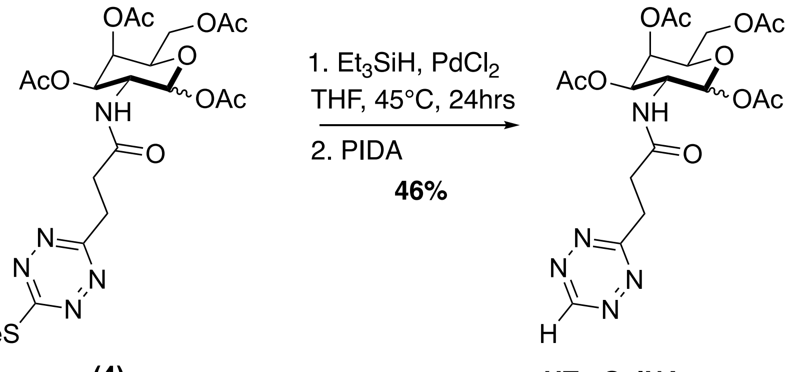


**(4)** **HTz-GalNAc**

A flame-dried Schlenk tube was charged with **4** (32.6 mg, 61.6 µmol, 1.0 eq) and palladium II chloride (2.9 mg, 16.4 µmol, 27 mol %, 0.27 eq) and evacuated and purged three times with $N_2$. Then triethylsilane (54.2 µL, 339.3 µmol, 5.5 eq) was added to anhydrous THF (1.13 mL, 0.05 M), the mixture was added to the Schlenk tube under $N_2$, and the suspension was allowed to stir at 45 °C for 20-24 hr. The reaction was cooled to RT and PIDA (54.7 mg, 170 µmol, 2.8 eq) was added and allowed to stir at RT for 1 hr. The suspension was then filtered through celite with ethyl acetate. The filtrate was washed with water and brine, then dried over sodium sulfate, filtered, and concentrated. A pink oil was isolated via flash chromatography (0 → 20 % Acetone in DCM), the oil was precipitated with a mixture of DCM : pentane and dried via rotatory evaporation/ high vac to yield a bright pink solid (13.6 mg, 46 %).

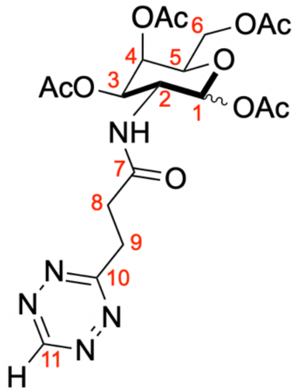


$^1H$ NMR (600 MHz, $CDCl_3$) $\delta$ (Anomers >9$\alpha$ : 1β) 10.19 (s, 1H, $\alpha H_{11}$), 5.71 (d, 1H, $J$ = 8.7 Hz, $\alpha H_1$), 5.68 (d, 1H, $J$ = 9.6 Hz, amide NH), 5.36 (d, 1H, $J$ = 3.2 Hz, $\alpha H_4$), 5.11 (dd, 1H, $J$ = 11.3, 3.5 Hz, $\alpha H_3$), 4.40 (dt, 1H, $J$ = 11.2, 9.3 Hz, $\alpha H_2$), 4.15 (dd, 1H, $J$ = 11.5, 6.7 Hz, $\alpha H_{6a}$), 4.10 (dd, 1H, $J$ = 11.5, 6.5 Hz, $\alpha H_{6b}$), 4.02 (t, 1H, $J$ = 6.5 Hz, $\alpha H_5$), 3.68 (td, 2H, $J$ = 6.8, 3.5 Hz, $\alpha H_{9a,b}$), 2.89 (t, 2H, $J$ = 6.5 Hz, $\alpha H_{8a,b}$), 2.14 (s, 3H, Acetyl-$CH_3$), 2.13 (s, 3H, Acetyl-$CH_3$), 2.04 (s, 3H, Acetyl-$CH_3$), 2.04 (s, 3H, Acetyl-$CH_3$)

$^{13}C$ NMR (151 MHz, $CDCl_3$) $\delta$ 171.6 (C), 171.3 (C), 171.0 (C), 170.6 (C), 170.3 (C), 169.8 (C), 158.2 (CH of Tetrazine), 93.4 (CH, $\alpha C_1$), 72.0 (CH, $\alpha C_5$), 70.4 (CH, $\alpha C_3$), 65.9 (CH, $\alpha C_4$), 61.5 ($CH_2$, $\alpha C_6$), 50.1 (CH, $\alpha C_2$), 32.1 ($CH_2$ next to Tz), 30.2 ($CH_2$ next to amide), 21.0 (2x acetyl-$CH_3$), 20.8 (2x acetyl-$CH_3$)

HRMS (ESI+) $[M + H]^+$ m/z calculated for $[C_{19}H_{26}N_5O_{10}]^+$ 484.167969, found 484.16897.

### Biochemistry Methods

*Flow Cytometry Measurement of Cell Surface HTz-GlcNAc and HTz-GalNAc*

WT or BOCTAG SaOS-2 cells were plated in a 6-well dish (1.5 x $10^5$ cell/well) and allowed to adhere overnight. The next day, each cell line was treated with HTz-GlcNAc (40 µM), HTz-GalNAc (40 µM), or an equal volume of DMSO as a vehicle control. After treatment, the cells were incubated overnight. On the third day, cells were lifted with 10 mM EDTA in Dulbecco's Phosphate Buffered Saline (DPBS, Sigma Aldrich and counted with the Countess cell counter. Cells were pelleted by centrifugation, then resuspended in DPBS containing 2% BSA (flow buffer) to a concentration of 1 x $10^6$ cells/mL. 2.5 x $10^5$ cells were used for each sample and placed in a v-bottom 96-well plate. Cells were washed three times in flow buffer. Cells were then resuspended in 1 µM TCO-CF647 (in flow buffer, Biotium) and incubated at RT for 5 min, shielded from light. Supernatant was removed, and cells were washed three more times in flow buffer. Cells were then resuspended in 400 µL of flow buffer and analyzed by flow cytometry (FACSCalibur). Samples were gated for single cells by forward and side scatter, and 10,000 cells were analyzed per sample using the FL4 channel (647 nm). Fold-change of gMFI between WT and BOCTAG SaOS-2 cells was calculated and analyzed using GraphPad Prism.

*In Gel Fluorescence Analysis of HTz-GlcNAc and HTz-GalNAc-Labeled Glycoproteins*

WT and BOCTAG SaOS-2 cells were plated in 6 cm dishes (3 x $10^5$ cells/dish) and incubated overnight. Cells were treated with HTz-GlcNAc (40 µM), HTz-GalNAc (40 µM), or an equal volume of DMSO as a vehicle control for 15 – 20 h. Cells were lifted with 10 mM EDTA in DPBS and washed 3 times with 2 % FBS in DPBS (cell buffer). Cells were resuspended in 1 µM TCO-CF647 in cell buffer for 5 min at RT, shielded from the light. Cells were pelleted by centrifugation, and supernatant was discarded. Cells were washed 3 times in ice-cold DPBS and lysed in RIPA (50 mM Tris pH 7.5, 150 mM NaCl, 0.1% SDS, 0.5% sodium deoxycholate, and 1% NP-40) with added HALT protease inhibitors (Thermo Fisher) and benzonase (1µL/mL, Sigma). Protein concentration of lysates was quantified by BCA, and lithium dodecyl sulfate (LDS) loading dye (NuPAGE, Invitrogen) with DTT was added. Samples were boiled for 5 minutes at 95 °C, 25 µg of protein/sample was loaded and resolved by gel electrophoresis (Invitrogen NuPAGE 4-12% Bis-Tris gel, MOPS running buffer). Gel fluorescence was imaged with a Bio Rad ChemiDoc imager. Gels were stained with Coomassie to confirm consistent protein loading.

*PNGaseF Release of N-Glycans from HTz-GlcNAc and HTz-GalNAc Labeled Glycoproteins*

Samples were prepared as above through BCA quantification of lysates. 20 µg protein/sample was treated with either PNGaseF (diluted 1:10 in DPBS, 10% of total reaction volume, Promega) or an equal amount of DPBS. Reactions were incubated overnight at 37 °C and quenched by boiling at 95 °C for 10s before immediately being placed on ice. Samples were prepared with LDS and DTT, resolved by gel electrophoresis, and imaged as described above.

*Kifunensine Treatment of BOCTAG Cells*

BOCTAG cells were plated in a 6-well plate, with 3 x $10^5$ cells/well and cultured for 16 h. Cells were treated with either DMSO as a vehicle control, HTz-GlcNAc (40 µM), or HTz-GlcNAc (40 µM) with varying concentrations of kifunensine (1, 5, and 10 µg/mL). Cells were cultured for 15-20 h and then harvested, labeled with TCO-CF647, and processed as described above, followed by in-gel fluorescence analysis.

*OSMI-1 Treatment of BOCTAG SaOS-2 Cells*

BOCTAG cells were plated in a 6-well plate, with 3 x $10^5$ cells/well and allowed to incubate overnight. The next day, cells were treated with either DMSO as a vehicle control, OSMI-1 (50 µM, MedChemExpress), and HTz-GlcNAc (50 µM). Cells were cultured with the compounds for 16 h and then harvested, labeled with TCO-CF647, and processed as described above, followed by in-gel fluorescence analysis. To confirm OSMI-1 efficacy, the aforementioned samples were resolved by SDS-PAGE, transferred to a PVDF membrane, blocked with CarboFree (Vector Labs) and probed with an O-GlcNAc antibody in CarboFree buffer overnight at 4 °C (CTD110.6, 1:500, Thermo Fisher). The membrane was washed 3 times in TBS-T (0.1% Tween) incubated with goat anti-mouse-HRP (1:5000) for 1 h at RT, washed 3X in TBS-T (0.1% Tween), developed with Pico ECL, and imaged (ChemiDoc, Bio Rad). The membrane was probed with anti-beta-actin-HRP as a loading control (1:5000, Cell Signaling Technology). Bands were quantified in FIJI and normalized to the loading control. Statistical analysis was performed in GraphPad Prism.

*Analysis of HTz-GlcNAc Impact on Fucosylation and Sialylation of Glycans in BOCTAG Cells*

Lysates from BOCTAG cells treated with either DMSO or HTz-GlcNAc were resolved by SDS-PAGE and transferred to PVDF membranes. Membranes were blocked with CarboFree Blocking Buffer (Vector Labs) and probed with either AAL-FITC (1:1000, Vector Labs) or Biotinylated PanSpecific SiaFind (1.25 µg/mL, Lectenz) overnight at 4 °C in CarboFree buffer. Membranes were washed 3 times with TBS-T (0.1% Tween). The AAL blot was imaged for fluorescence (ChemiDoc, BioRad), and the SiaFind blot was incubated with Streptavidin-POD (1:1000) for 1 h at RT, before washing 3 times with TBS-T (0.1% Tween), developing with Pico ECL, and imaging chemiluminescence (ChemiDoc, BioRad). For loading controls, blots were incubated for 1 h at RT with GAPDH-HRP (1:5000, Cell Signaling Technologies). Band quantification was performed FIJI, and bands were normalized to GAPDH. Statistical analysis was performed using GraphPad Prism.

*Analysis of HTz-GlcNAc Toxicity on WT and BOCTAG Cells with CellTiter-Glo*

To assess the toxicity of HTz-GlcNAc to WT and BOCTAG cells, cells were plated in a white-walled 96-well plate (Corning, #3610) at a density of 3000 cells/well in DMEM. Cells were allowed to adhere to the plate overnight and treated the next day with varying concentrations of HTz-GlcNAc or DMSO as a vehicle control. Cells were cultured with the compound for 20 h, and the CellTiter-Glo 2.0 (Promega) assay was performed according to the manufacturer's specifications. Luminescence data was normalized to the DMSO-treated control, and statistical analysis was performed in GraphPad Prism.

*Metabolomics Sample Preparation*

Samples were prepared according to a standard protocol.[6] Briefly, 1 x$10^6$ WT or BOCTAG SaOS2 cells were plated in 6-cm tissue culture plates and allowed to adhere for 16 h. The next day, cells were treated with HTz-GlcNAc (50 µM) or DMSO, diluted in DMEM, and allowed to incubate for 20 h. Cells were washed with normal saline then scraped into 80% (vol/vol) acetonitrile in water. The cell lysate/acetonitrile mixture was subjected to three freeze-thaw cycles before centrifugation at 21,000 *g*. The metabolite-containing supernatant was collected, and BCA was performed to quantify and normalize sample concentrations. Volumes equivalent to 20 µg protein/sample were diluted to 100 µL, and samples were vortexed and centrifuged again. The supernatants were collected and stored at –80 °C until analysis.

*Metabolomics Analysis for Detection and Quantification of UDP-HTz-GlcNAc*

Metabolomic data were acquired with a Thermo Fisher Scientific Orbitrap Fusion Lumos mass spectrometer (LC–MS/MS). A MS2 negative or positive method was set up for UDP-HTz-

HexNAc and UDP-HexNAc. Precursor MS data were acquired were acquired with a resolving power of 120,000 full width at half maximum (FWHM) with a scan range set to 80–1,200 daltons; the automatic gain control (AGC) target was set to 400,000 with an automatic maximum inject time. MS2 spectra were acquired with a data-dependent method in individual polarities and were acquired with a resolving power of 15,000 FWHM; the AGC target was set to 50,000, with an automatic maximum inject time. A top-10 data-dependent MS schema was used with an isolation window of 0.7 Da. Analytes were fragmented with a collision energy of 30 normalized collision energy (NCE). A MS3 method was set up for UDP-HTz-HexNAc and UDP-HexNAc identification with a targeted method in negative polarity. Precursor MS data were acquired with a resolving power of 60,000 full width at half maximum (FWHM) with a scan range set to 80–1,200 daltons; the automatic gain control (AGC) target was set to 400,000 with an automatic maximum inject time. Precursor ions 702.1168 (UDP-HTz-GlcNAc ) and 608.0888 (UDP-GlcNAc ) were selected for fragmentation in MS2 level, and product ion MS data were acquired with a resolving power of 30,000 FWHM; the AGC target was set to 50,000, with an automatic maximum inject time with an isolation window of 0.7 Da. Analytes were fragmented with a collision energy of 30 normalized collision energy (NCE). Then, 298.1151 (a fragment of UDP-HTz-HexNAc ) and 204.0866 (a fragment of UDP-HexNAc) were selected for fragmentation in MS3 level, and product ion MS data were acquired with a resolving power of 30,000 FWHM; the AGC target was set to 100,000, with an automatic maximum inject time with isolation windows of 2.5 and 2.5 Da for MS1 and MS2 respectively. Analytes were fragmented with a collision energy of 30 normalized collision energy (NCE). Metabolites were chromatographically resolved using published methods.[7] The percent of HTz-modified UDP-HexNAc compared to total UDP-HexNAc was calculated, and the data was plotted in GraphPad Prism.

*Enrichment and Proteomics Analysis of HTz-GlcNAc-Labeled Proteins*

WT and BOCTAG SaOS-2 cells were plated in 15 cm tissue culture plates (5 x $10^6$ cells/dish) and cultured for 15-20 h. The next day, all cells were treated with HTz-GlcNAc (40 μM) for 15-20 h. Cells were harvested with EDTA (10 mM in DPBS) and washed 3 times in cold PBS. For whole lysate labeling, cells were lysed in RIPA (with HALT protease inhibitors and benzonase), and protein concentration was quantified by BCA. For cell-surface-specific enrichment and analysis, cells were lifted and washed as above, followed by labeling with TCO-Biotin (10 μM; Biotium) for 15 min at room temperature, with gentle shaking. Cells were then washed 3 times with cold DPBS and lysed in RIPA and quantified by BCA. Protein (300 μg/sample in 250 μL) was incubated with washed neutravidin beads (Cytiva) for 1.5 h at room temperature, rotating end-over-end, to remove endogenously biotinylated proteins. The supernatant was collected, and volume was adjusted to 270 μL with DPBS. Lysates were labeled with TCO-Biotin (10 μM; Biotium) for 45 min at room temperature, shaking at 400 rpm. Samples were precipitated overnight with methanol (10-fold excess) at –80 °C. The next day, precipitates were pelleted for 20 min at 3700*g* at 4 °C. Samples were washed twice with ice-cold methanol, pelleting between each wash. Pellets were air-dried on the bench until no visible methanol remained. Pellets were resuspended in RapiGest (0.1% in PBS; Waters) and sonicated (VWR Symphony) for 25 min at RT, followed by centrifugation for 5 min at 3700*g* at room temperature. The RapiGest supernatant was collected and incubated at 4 °C. Pellets were resuspended in urea (6 M in DPBS) and sonicated (VWR Symphony) for 25 min at RT, followed by centrifugation. The urea supernatant was collected and incubated at 4 °C. Finally, pellets were resuspended in DPBS and sonicated for 25 min, followed by centrifugation, and all supernatants were combined and added to washed lys-dimethylated neutravidin beads (Cytiva). Samples were incubated with the beads for 2 h at room temperature, rotating end-over-end. After 2 h, the supernatant was removed, and beads were washed three times with SDS (0.05 % w/v in PBS), six times with urea (6 M in PBS), six times with AmBic (50 mM in LC/MS grade water), four times with LC/MS-grade acetonitrile in LC/MS-grade water (40 % v/v). All liquid was aspirated from the samples, beads were pelleted by centrifugation and stored at -20 °C until on-bead digestion and proteomics analysis.

*On-Bead Digestion, Proteomics Data Acquisition, and Peptide Identification and Quantification*

100 µL of 2 M urea in 100 mM Tris-HCl was added to cover the beads. To this, 4 µL of 500 mM of TCEP was added and incubated at RT for 30 mins. Following this, 4 µL of 500 mM IAA was added and samples were incubated at RT in the dark with shaking. The liquid was discarded and 1 µg trypsin in 80 µL 100 mM Tris-HCl pH? was added and allowed to shake overnight at 37 °C. The solution was transferred to a new tube. To the beads, 50 µL of 0.1 % TFA was added and incubated for 5 min with shaking. This solution was added to the previous one. Following solid-phase extraction cleanup with an Oasis HLB µelution plate (Waters), the resulting peptides were reconstituted in 10 µL of 2 % (v/v) acetonitrile (ACN) and 0.1 % trifluoroacetic acid in water. 2 µL of each sample were injected onto a Q Exactive HF mass spectrometer coupled to an Ultimate 3000 RSLC-Nano liquid chromatography system. Samples were injected onto a 75 µm i.d., 15-cm long EasySpray column (Thermo) and eluted with a gradient from 0 – 28 % buffer B over 90 min. Buffer A contained 2 % (v/v) acetonitrile and 0.1 % formic acid in water, and buffer B contained 80 % (v/v) acetonitrile, 10 % (v/v) trifluoroethanol, and 0.1 % formic acid in water. The mass spectrometer operated in positive ion mode with a source voltage of 2.5 kV and an ion transfer tube temperature of 300 °C. MS scans were acquired at 120,000 resolution in the Orbitrap and up to 20 MS/MS spectra were obtained in the ion trap for each full spectrum acquired using higher-energy collisional dissociation (HCD) for ions with charges 2-8. Dynamic exclusion was set for 20 s after an ion was selected for fragmentation.

Raw MS data files were analyzed using Proteome Discoverer v3.0 (Thermo), with peptide identification performed using Sequest HT searching against the human reviewed protein database from UniProt (downloaded January 4, 2024; 20,354 entries). Fragment and precursor tolerances of 10 ppm and 0.6 Da were specified, and three missed cleavages were allowed. Carbamidomethylation of Cys was set as a fixed modification and oxidation of Met was set as a variable modification. The false-discovery rate (FDR) cutoff was 1 % for all peptides. Peptide peak intensities were summed for all peptides matched to a protein for protein quantitation.

*Enrichment and Immunoblot Analysis of HTz-GlcNAc-Labeled Proteins*

WT and BOCTAG SaOS-2 cells were plated in 6 cm dishes (5 x $10^5$ cells/dish) and incubated overnight. All plates were treated with HTz-GlcNAc (40 µM) for 15-20 h. The next day, cells were harvested with EDTA (10 mM in DPBS) and washed three times with ice-cold DPBS. Cell pellets were lysed in RIPA (with HALT protease inhibitors and benzonase). Protein concentration was quantified by BCA (Pierce, 23225). Protein from each sample (150 µg) was labeled with TCO-Biotin (10 µM; Biotium) for 45 min at room temperature with shaking (400 rpm). Samples were diluted to 375 µL in DPBS and incubated on washed neutravidin beads (Cytiva) for 2 h at room temperature, with end-over-end rotation. The supernatant was removed, and beads were washed three times with PBS containing 0.05 % Tween. Samples were eluted by boiling in SDS loading dye for 10 min with shaking (400 rpm). For immunoblot analysis, 10 µg protein/sample was loaded for each input, and 25 % of the eluted volume was loaded per enriched sample. Samples were resolved by SDS-PAGE and transferred to PVDF membranes. Membranes were blocked with non-fat milk (3 % in TBS containing 0.1% Tween) for 1 h at room temperature and incubated with the following primary antibodies overnight at 4 °C: anti-LAMP1 (1:1000) and anti-IntegrinB1 (1:1000). Blots were washed three times in TBS containing 0.1% Tween. All blots were incubated with goat-anti rabbit-HRP secondary antibody (1:5000) for 1 h at room temperature and washed three times in TBS containing 0.1% Tween. Blots were developed with Pico ECL and imaged (BioRad ChemiDoc).

*Live Cell Confocal Fluorescence Microscopy of HTz-GlcNAc*

WT and BOCTAG SaOS-2 cells were plated in 35 mm Mattek dishes (No. 1.5 coverslip, 14 mm glass diameter, 8 x $10^4$ cells/dish) and incubated overnight at 37 °C with 5 % $CO_2$. All plates were treated with HTz-GlcNAc (50 µM) or an equivalent volume of DMSO as a vehicle control for 20-24 h at 37 °C with 5 % $CO_2$. The next day, cells were washed 2 times for 10 min with media, treated with 1 µM TCO-fluorophore in media (1 mL), and incubated at 37 °C for 30 min. Cells were washed 3 times for 30 min in media, followed by 2 rinses with 1 mL Gibco Opti-MEM reduced-serum media, and the cells were imaged live in fresh Opti-MEM. All imaging was done with live cells at ambient conditions (RT, open to air).

*Dual Labeling Confocal Fluorescence Microscopy of HTz-GlcNAc*

WT and BOCTAG SaOS-2 cells were plated in 35 mm Mattek dishes (No. 1.5 coverslip, 14 mm glass diameter, 8e4 cells/dish) and incubated overnight at 37 °C with 5 % $CO_2$. All plates were treated with HTz-GlcNAc (50 µM) or an equivalent volume of DMSO as a vehicle control for 20-24 h at 37 °C with 5% $CO_2$. The next day, cells were washed 2 times for 10 min with media, treated with 1 µM TCO-fluorophore #1 in media (1 mL), and incubated at 37 °C for 30 min. Cells were washed 2 times for10 min in media, treated with 1 µM TCO-fluorophore #2 in media (1 mL), and incubated at 37 °C for 30 min. Cells were washed 2 times for 20 min in media, followed by 2 rinses with 1 mL Gibco Opti-MEM reduced serum media, and imaged live in fresh Opti-MEM. All imaging was done with live cells at ambient conditions (RT, open to air).

*Multiday Time-Point Labeling Confocal Fluorescence Microscopy of HTz-GlcNAc*

BOCTAG SaOS-2 cells were plated in 35 mm Mattek dishes (No. 1.5 coverslip, 14 mm glass diameter, 8 x$10^4$ cells/dish for 24-h samples, 3 x $10^4$ cells/dish for 48-h and 72-h samples) and incubated overnight at 37 °C with 5 % $CO_2$. All plates were treated with HTz-GlcNAc (50 µM, Metabolic Incorporation # 1) or an equivalent volume of DMSO as a vehicle control for 20-24 hr at 37 °C with 5% $CO_2$. At 24 h, all samples were washed 2 times for 10 min with media, treated with 1 µM TCO-AF488 and Hoechst (15 µM, for imaging samples only), and incubated at 37 °C for 30 min. The samples to be imaged at 24 h were washed 3 times for 30 min in media at 37 °C, rinsed twice with Opti-MEM, and imaged live in fresh Opti-MEM. The 48- and 72-h samples were washed 3 times for 10 min in media, incubated with commercial methylphenyl tetrazine-amine (50 µM) for 30 min at 37 °C, and washed 3 times for 10 min in media. Samples were then treated with or without HTz-GlcNAc (50 µM, Metabolic Incorporation #2) in media and incubated overnight at 37 °C with 5 % $CO_2$. At 48 h, all samples were washed 2 times for 10 min with media, treated with 1 µM Sulfo-PEG2-Cy3-TCO and Hoechst (15 µM, for imaging samples only), and incubated at 37 °C for 30 min. The samples to be imaged at 48 h were washed 3 times for 30 min in media at 37 °C, rinsed twice with Opti-MEM, and imaged live in fresh Opti-MEM. The 72-h samples were washed 3 times for 10 min in media, incubated with commercial methylphenyl tetrazine-amine (50 µM) for 30 min at 37 °C, and washed 3 times for 10 min in media. Samples were then treated with or without HTz-GlcNAc (50 µM, Metabolic Incorporation #3) in media and incubated overnight at 37 °C with 5 % $CO_2$. At 72 h, all samples were washed 2 times for 10 min with media, treated with 1 µM Sulfo-Cy5-TCO and Hoechst (15 µM), and incubated at 37 °C for 30 min. The samples were washed 3 times for 30 min in media at 37 °C, rinsed twice with Opti-MEM, and imaged live in fresh Opti-MEM. All imaging was done with live cells at ambient conditions (RT, open to air).

Sample Breakdown:

1x = Metabolic Incorporation of HTz-GlcNAc #1 (0-24 h)

2x = Metabolic Incorporation of HTz-GlcNAc #1 (0-24 h) + Metabolic Incorporation of HTz-GlcNAc #2 (28-48 h)

3x = Metabolic Incorporation of HTz-GlcNAc #1 (0-24 h) + Metabolic Incorporation of HTz-GlcNAc #2 (28-48 h) + Metabolic Incorporation of HTz-GlcNAc #3 (52-72 h)

*24hr:* Labeling with TCO-AF488 (green) at 24 h

*48hr:* Labeling with TCO-AF488 (green) at 24 h + Labeling with Sulfo-PEG2-Cy3-TCO (orange) at 48 h

*72hr:* Labeling with TCO-AF488 (green) at 24 h + Labeling with Sulfo-PEG2-Cy3-TCO (orange) at 48 h + Labeling with Sulfo-Cy5-TCO (pink) at 72 h

*Live Cell Confocal Fluorescence of HTz-GlcNAc with Vesicle Markers*

Plasmids for GFP-tagged vesicle markers were obtained from Addgene (LAMP1 for lysosome #34831, Rab7 for endosome #12605, SEC31A for COPII vesicles (66613), and TGOLN2 for Trans Golgi Network vesicles #54279), sequence-verified, and mini-prepped (Wizard Plus SV Minipreps, Invitrogen). BOCTAG SaOS-2 cells were transfected with the respective plasmids using Lipofectamine 3000 (Thermo Fisher), cultured with 40 μM HTz-GlcNAc for 16 h, and labeled with TCO-SiR, as described above. Cells were imaged in biological triplicate as described above using the LSM880 confocal microscope with a 63x objective, with 3 fields of view per biological sample.

*Live Cell Confocal Fluorescence Microscopy of HTz-GlcNAc with Brefeldin A Treatment*

BOCTAG SaOS-2 cells were plated in 35 mm Mattek dishes (No. 1.5 coverslip, 14 mm glass diameter, 8 x $10^4$ cells/dish) and incubated overnight at 37 °C with 5 % $CO_2$. All plates were treated with HTz-GlcNAc (50 μM) (or an equivalent volume of DMSO as a vehicle control) for 20-24 h at 37 °C with 5 % $CO_2$. The overnight Brefeldin A sample was treated with 1.5 μg/mL Brefeldin A during the HTz-GlcNAc incubation. The next day, all cells were washed 2 times for 10 min with media, treated with 1 μM aTCO-SiR in media (1 mL), and incubated at 37 °C for 30 min. Cells were washed 3 times for 30 min in media, followed by 2 rinses with 1 mL Gibco Opti-MEM reduced-serum media, and the cells were imaged live in fresh Opti-MEM. The 30 min Brefeldin A sample was treated with 1.5 μg/mL during the last 30 min wash in media just prior to imaging. All imaging was done with live cells at ambient conditions (RT, open to air).

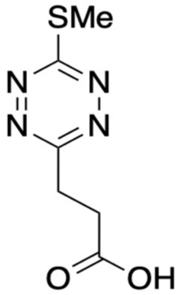

**(1)**

**$^{1}$H NMR (400 MHz, $CDCl_3$)**

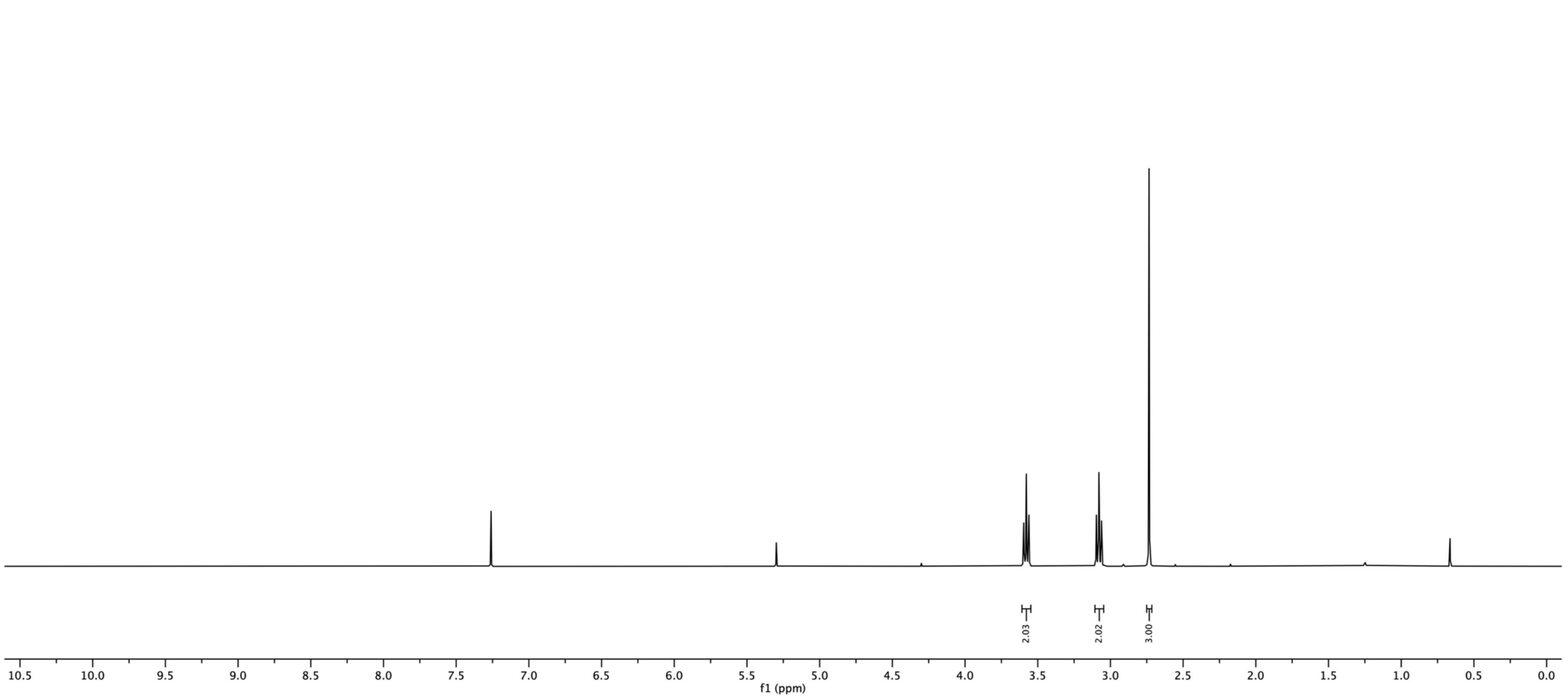

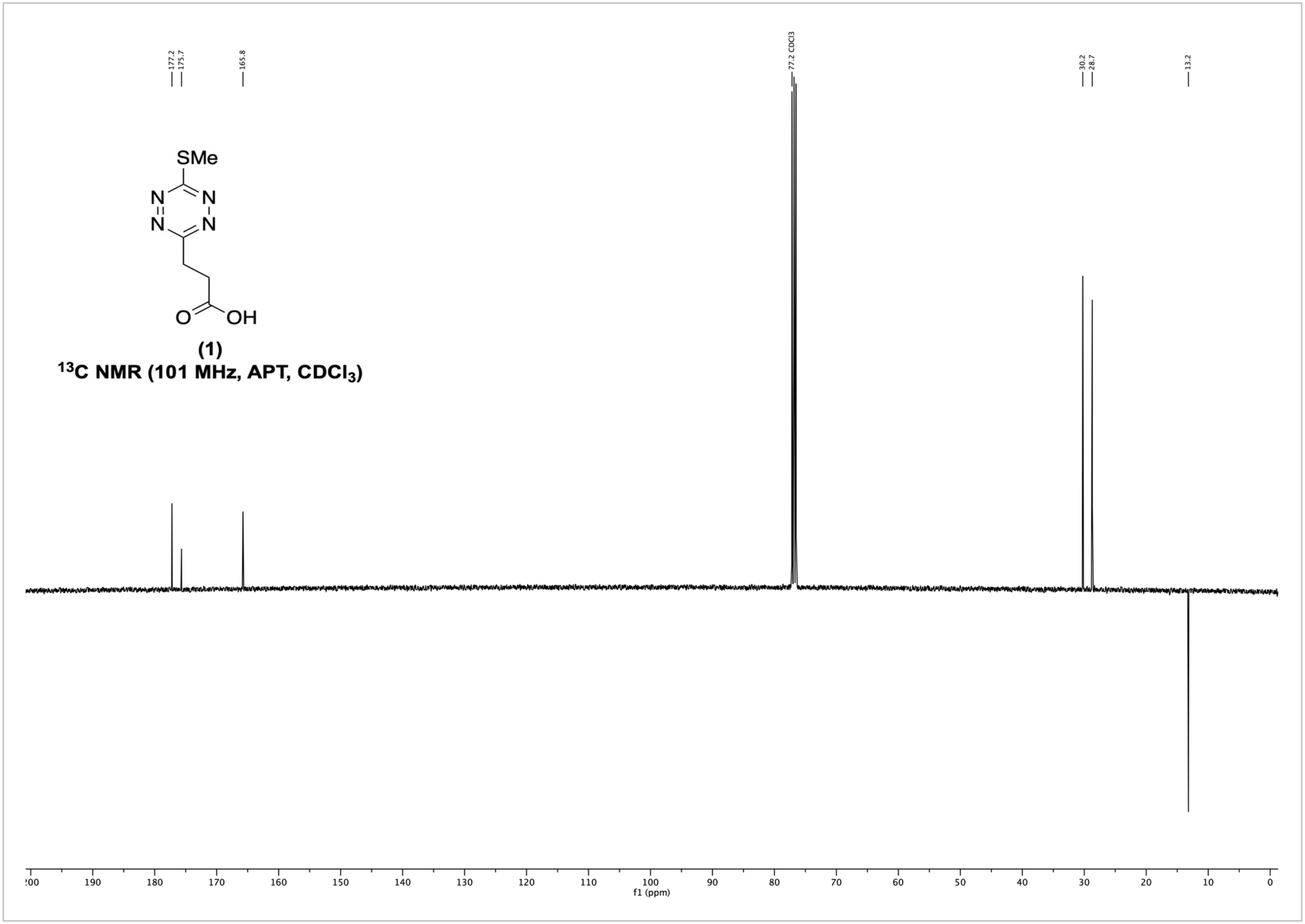
177.2
175.7
165.8
77.2 CDCl3
30.2
28.7
13.2
SMe
N
N
N
N
O
OH
(1)
$^{13}$C NMR (101 MHz, APT, $CDCl_3$)
f1 (ppm)

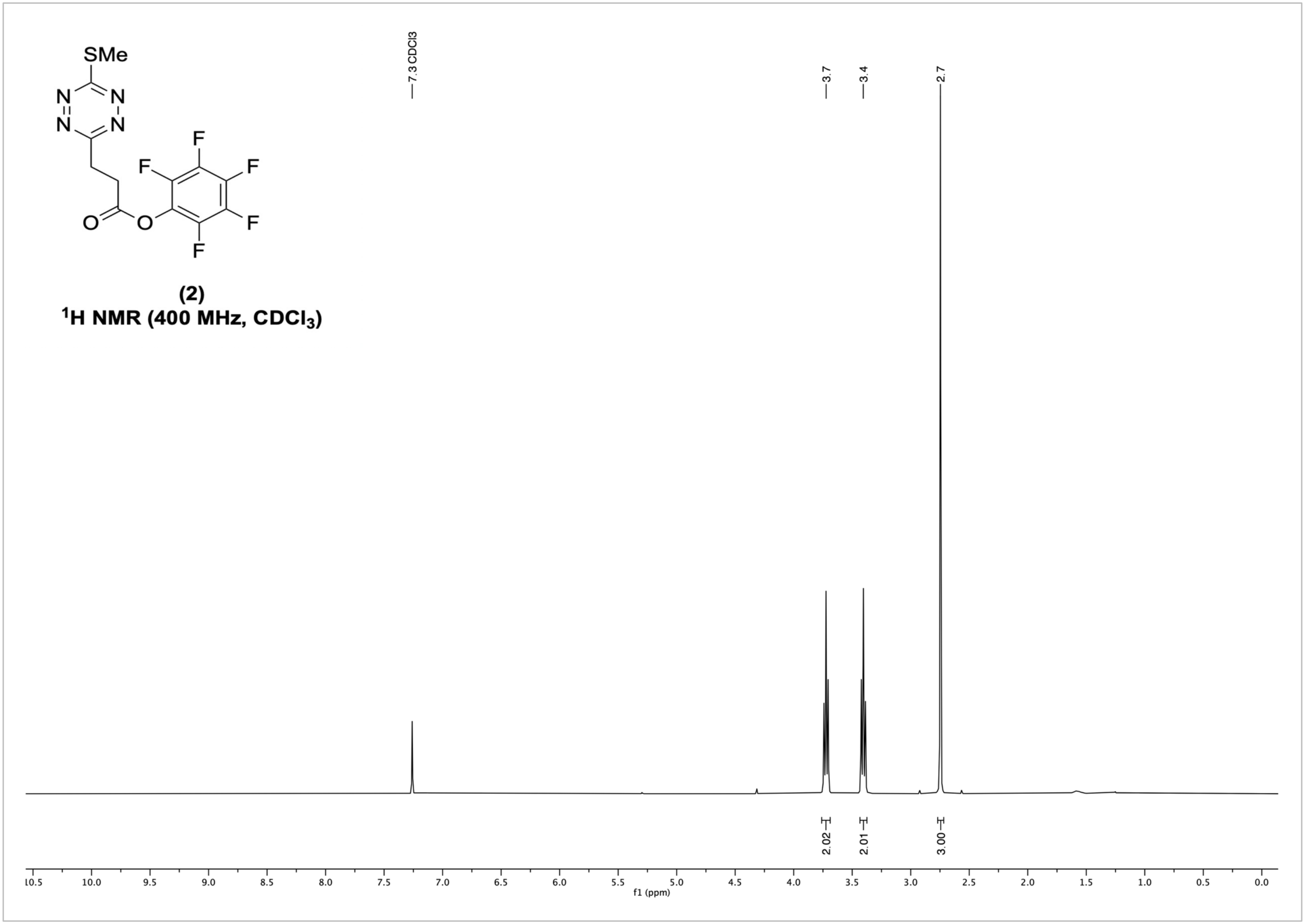
SMe
N
N
N
N
F
F
F
F
F
O
O
(2)
1H NMR (400 MHz, CDCl3)
7.3 CDCl3
3.7
3.4
2.7
2.02
2.01
3.00
f1 (ppm)

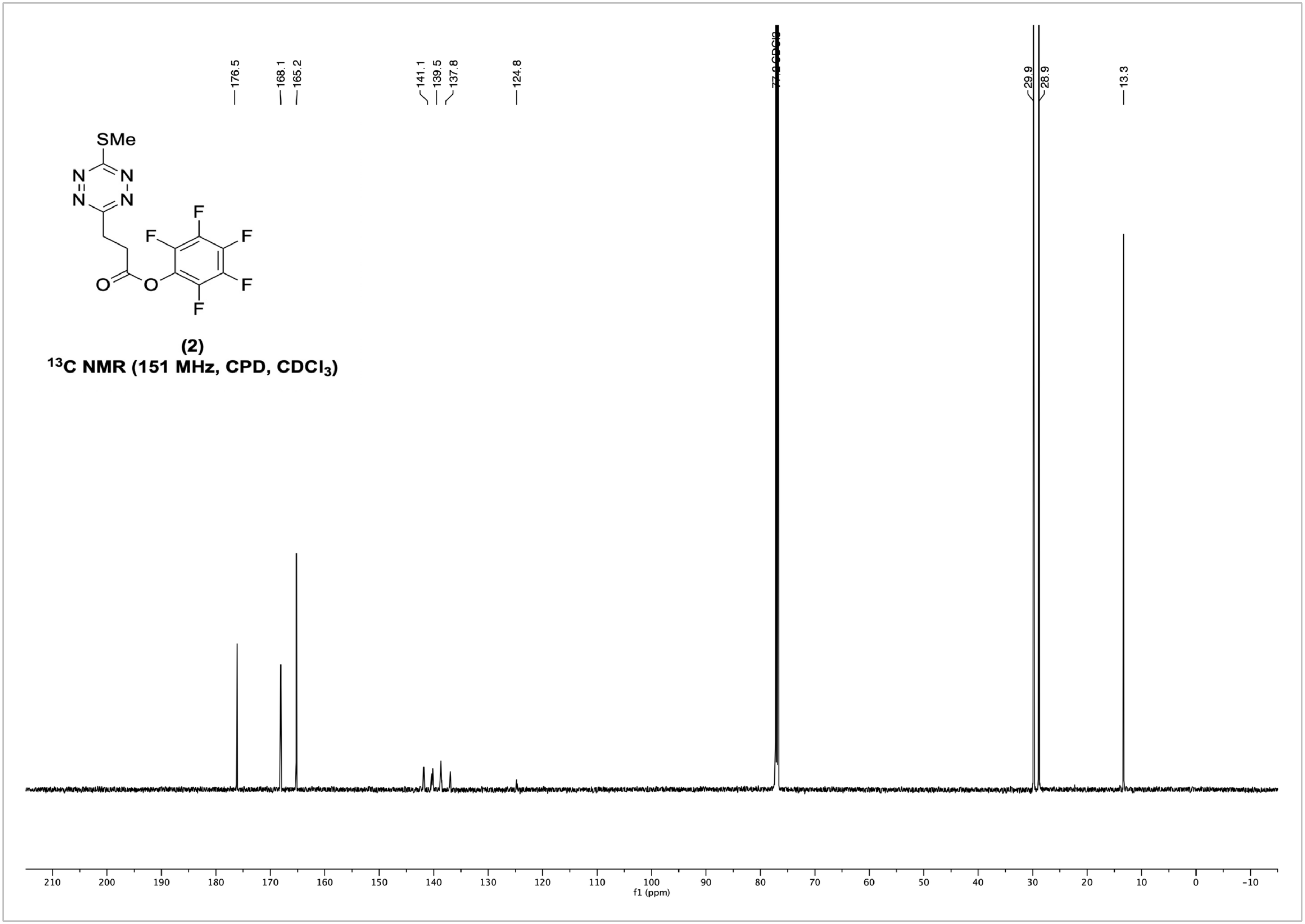


**(2)**

**$^{13}$C NMR (151 MHz, CPD, $CDCl_3$)**

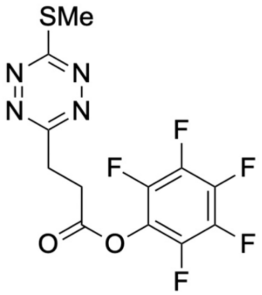

**(2)**

**$^{13}C$ NMR (151 MHz, C13FCPD, $CDCl_3$)**

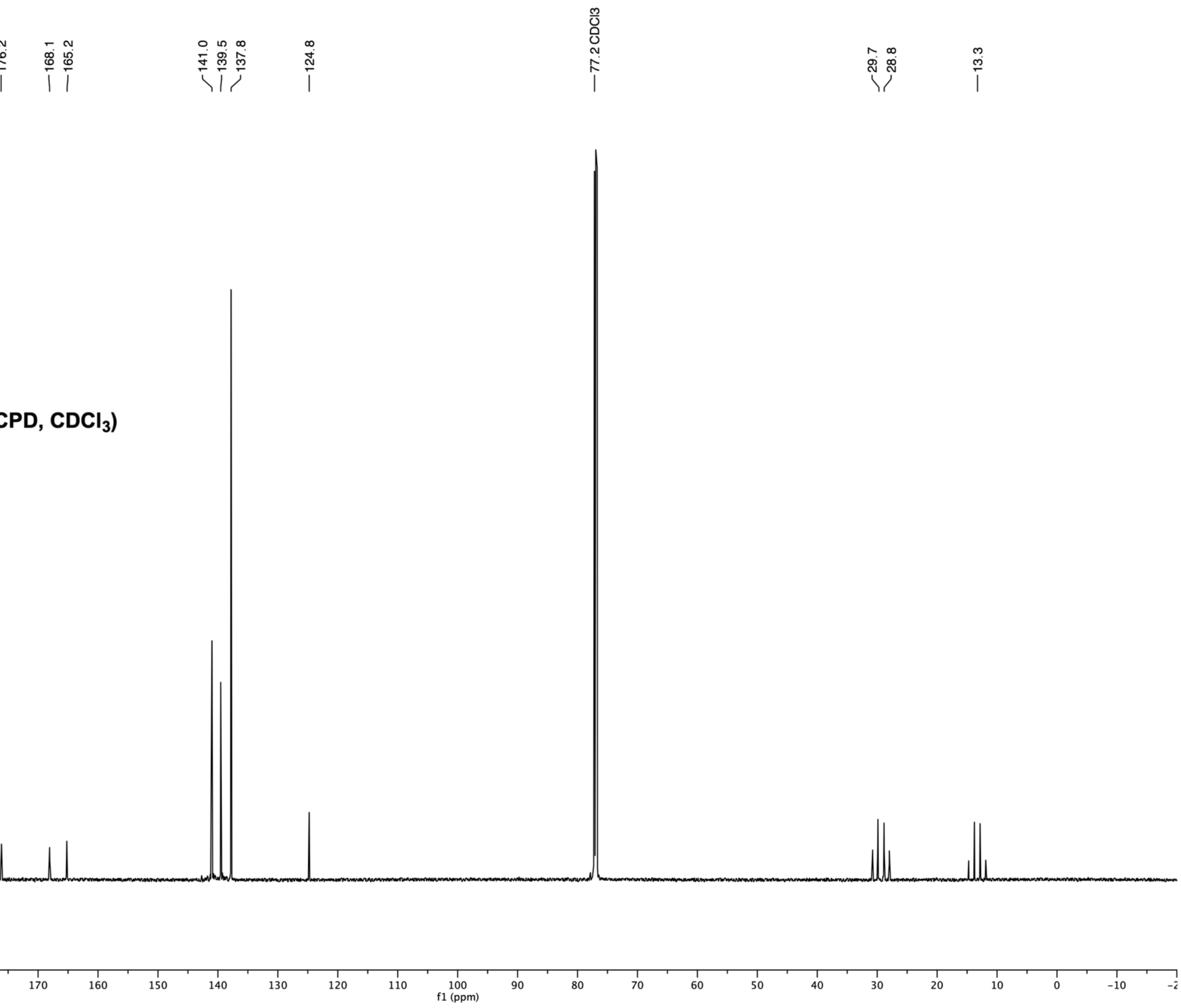

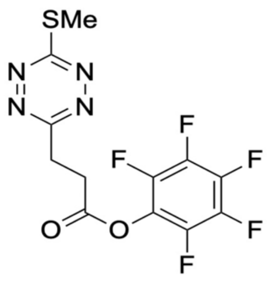

**(2)**

**$^{13}$C NMR (151 MHz, CPD, $CDCl_3$)**

**$^{13}$C NMR (151 MHz, C13FCPD, $CDCl_3$)**

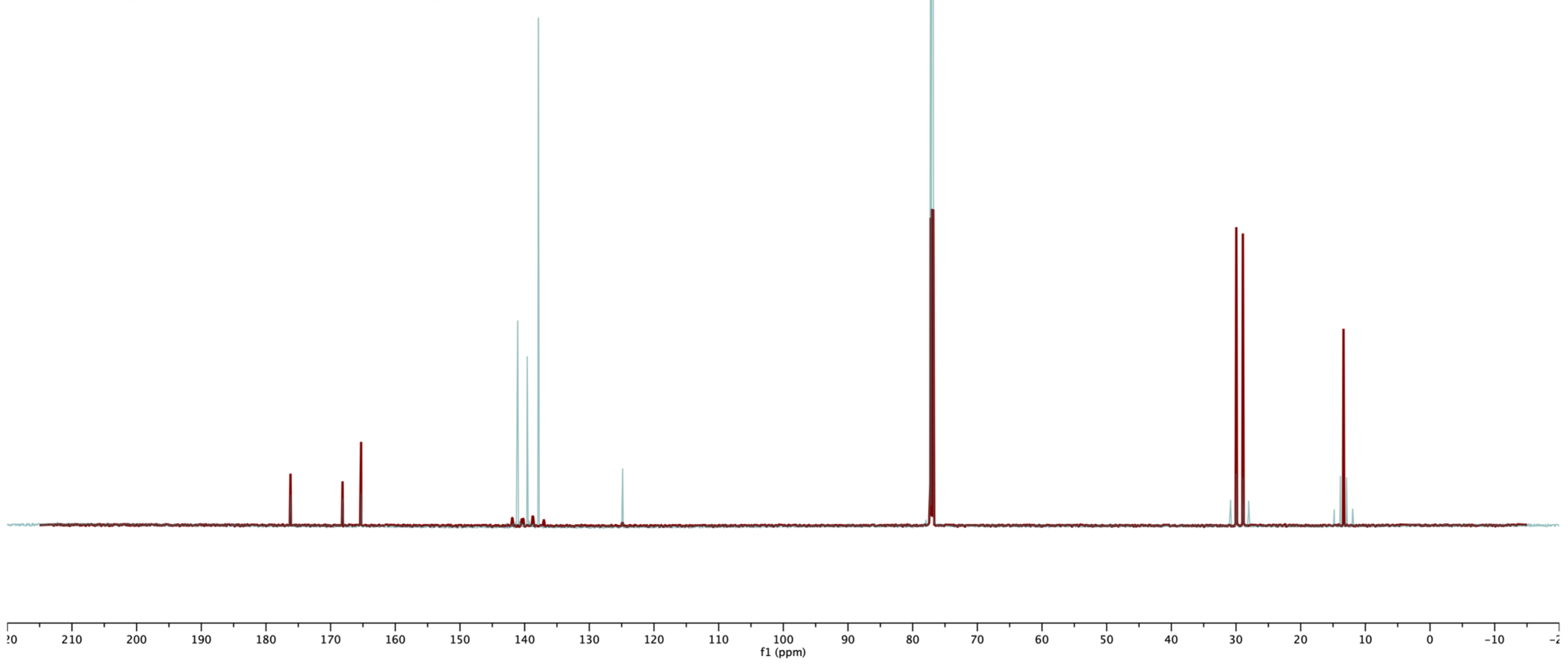

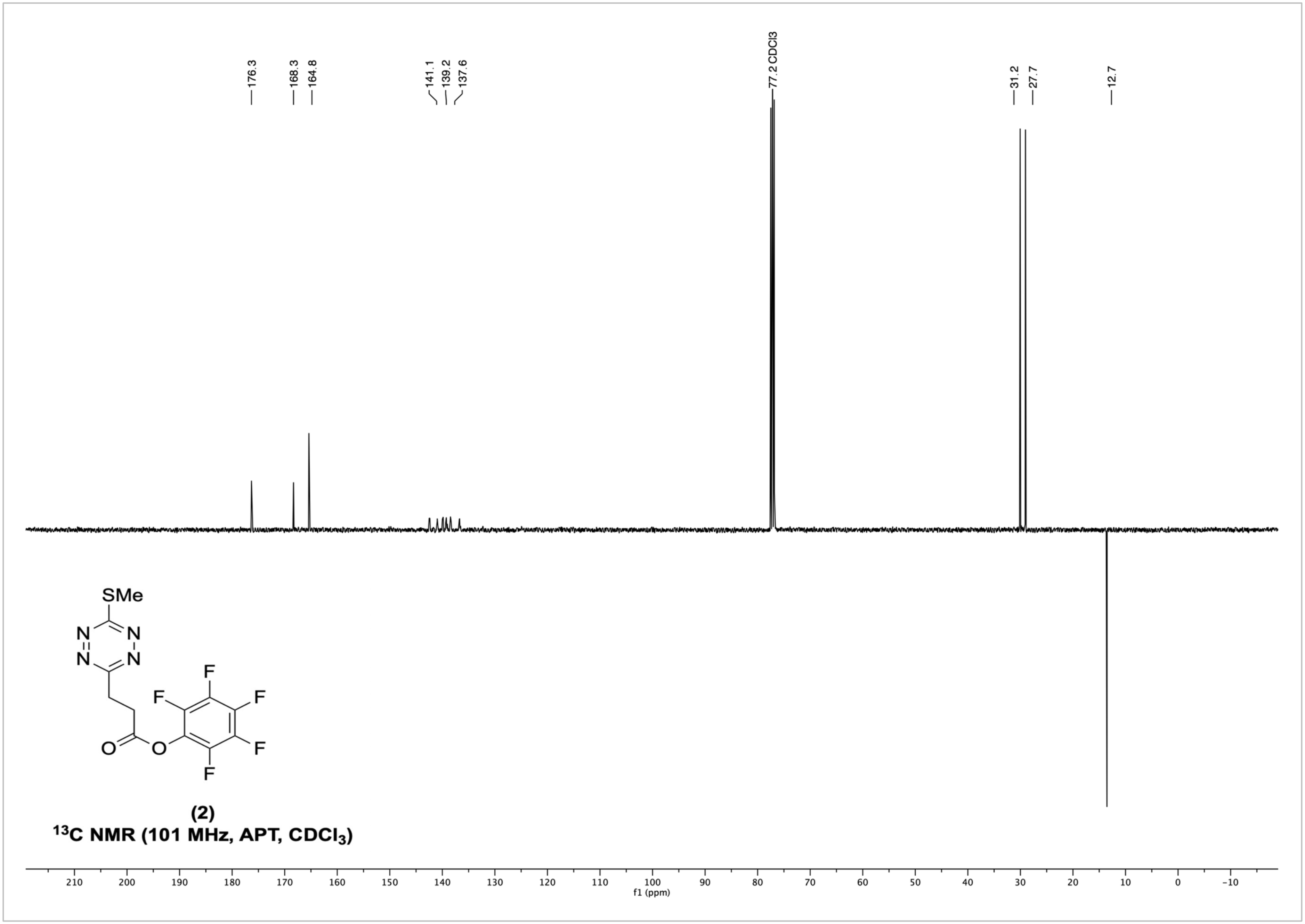


(2)

$^{13}C$ NMR (101 MHz, APT, $CDCl_3$)

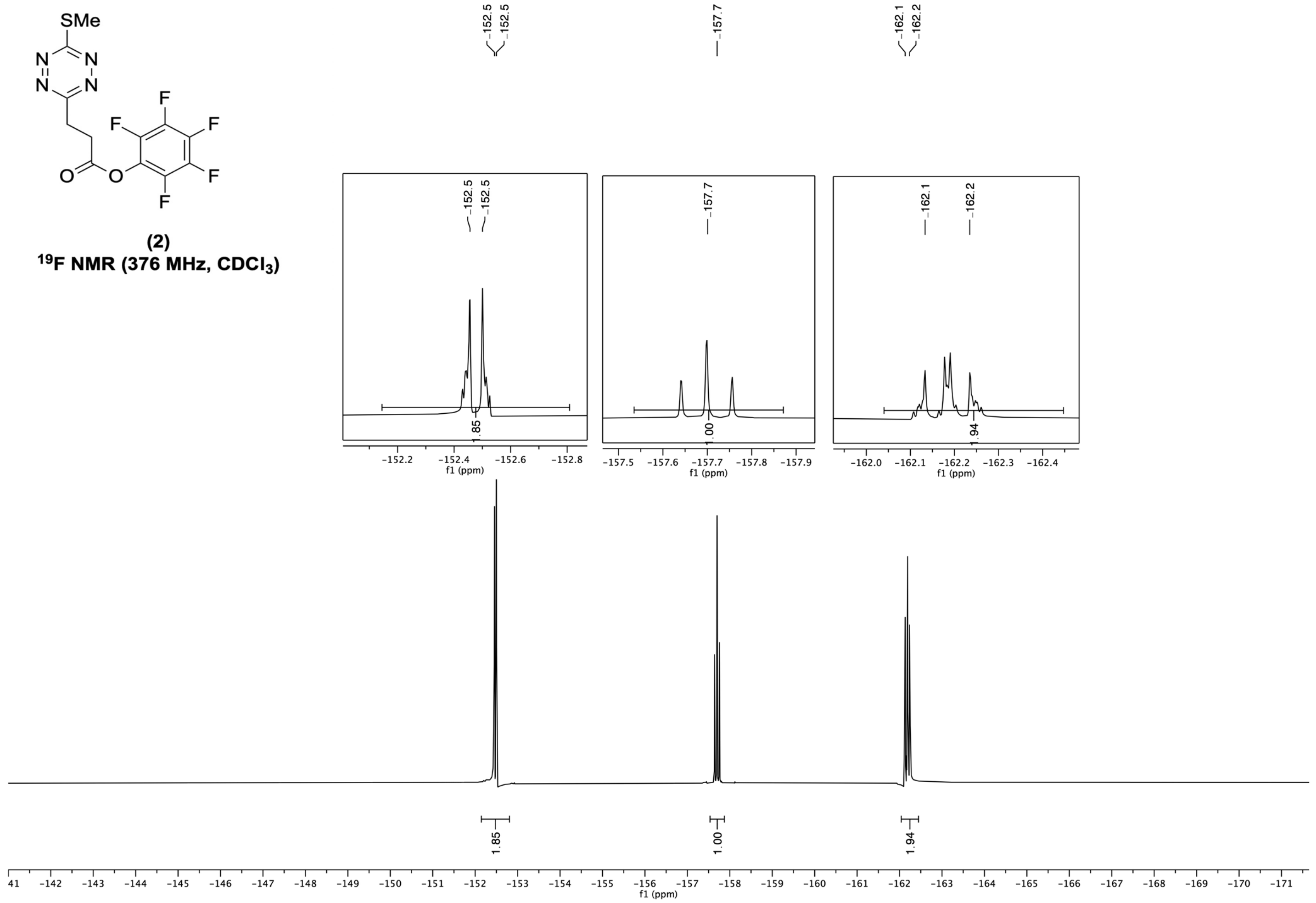


**(2)**
**$^{19}$F NMR (376 MHz, $CDCl_3$)**

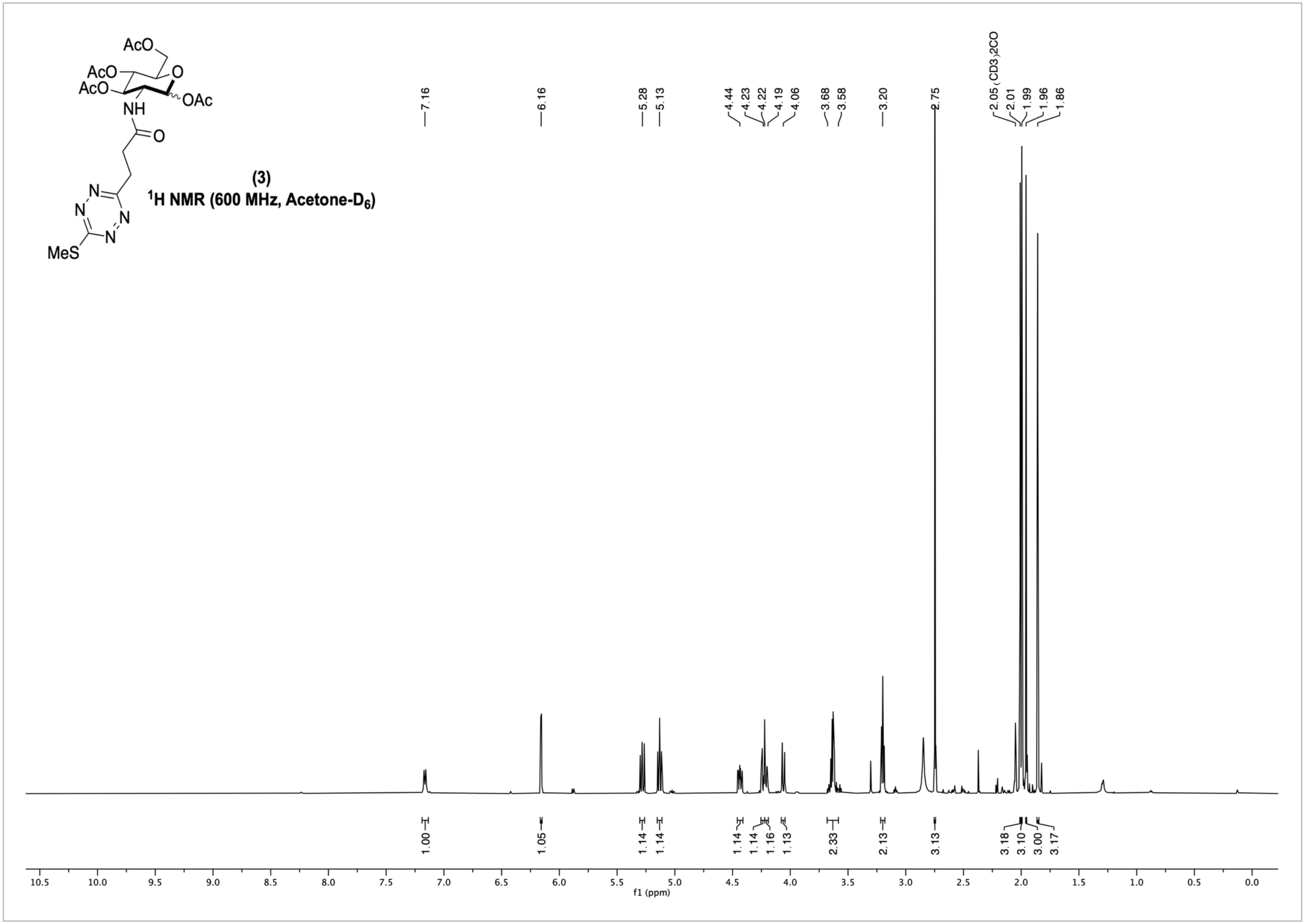
(3)
1H NMR (600 MHz, Acetone-D6)
7.16
6.16
5.28
5.13
4.44
4.23
4.22
4.19
4.06
3.68
3.58
3.20
2.75
2.05 (CD3)2CO
2.01
1.99
1.96
1.86
1.00
1.05
1.14
1.14
1.14
1.14
1.16
1.13
2.33
2.13
3.13
3.18
3.10
3.00
3.17
f1 (ppm)

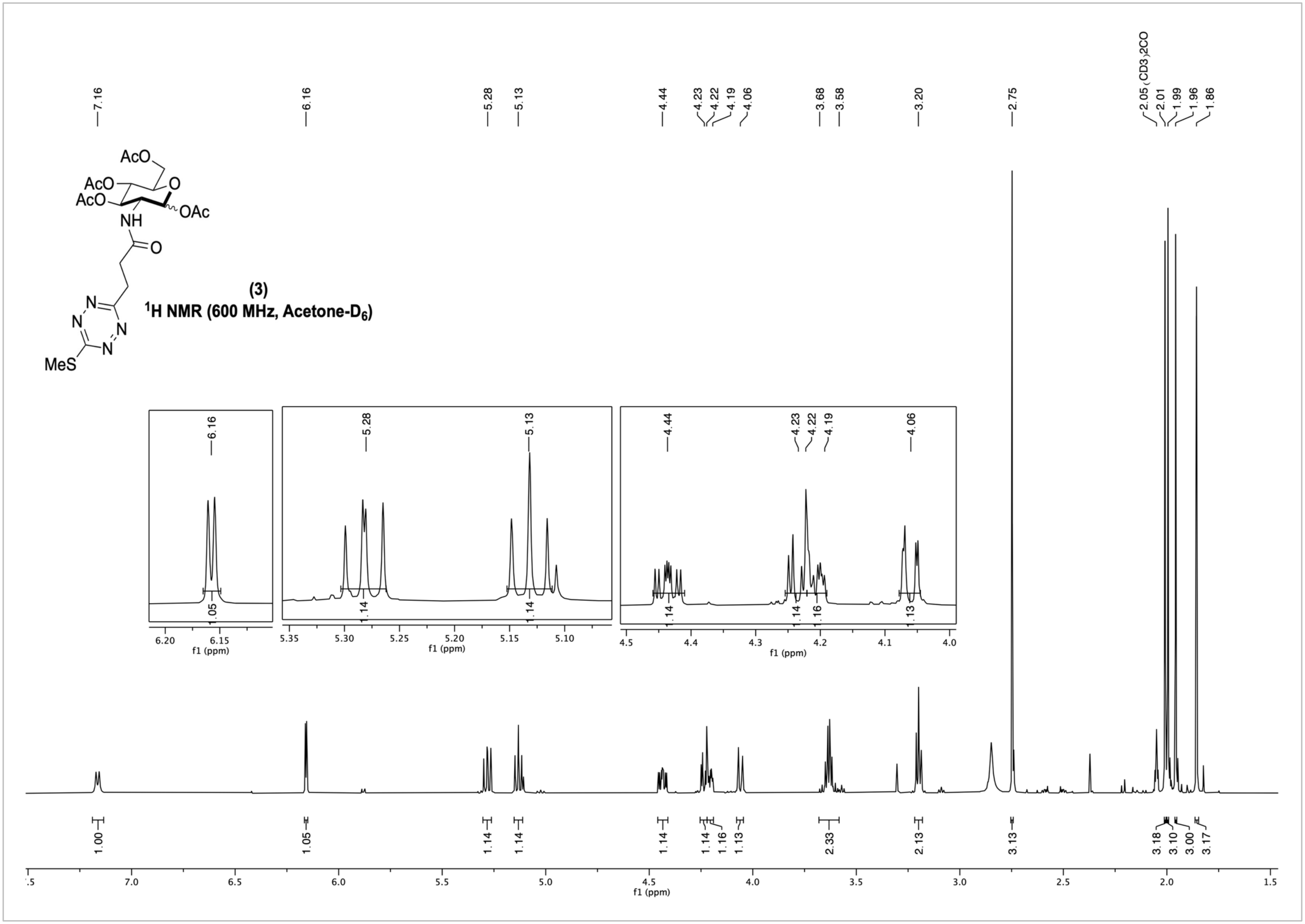
AcO
AcO
AcO
NH
O
OAc
O
N
N
N
N
MeS
(3)
$^1$H NMR (600 MHz, Acetone-D$_6$)
7.16
6.16
5.28
5.13
4.44
4.23
4.22
4.19
4.06
3.68
3.58
3.20
2.75
2.05 (CD3)2CO
2.01
1.99
1.96
1.86
1.00
1.05
1.14
1.14
1.14
1.14
1.16
1.13
2.33
2.13
3.13
3.18
3.10
3.00
3.17
f1 (ppm)

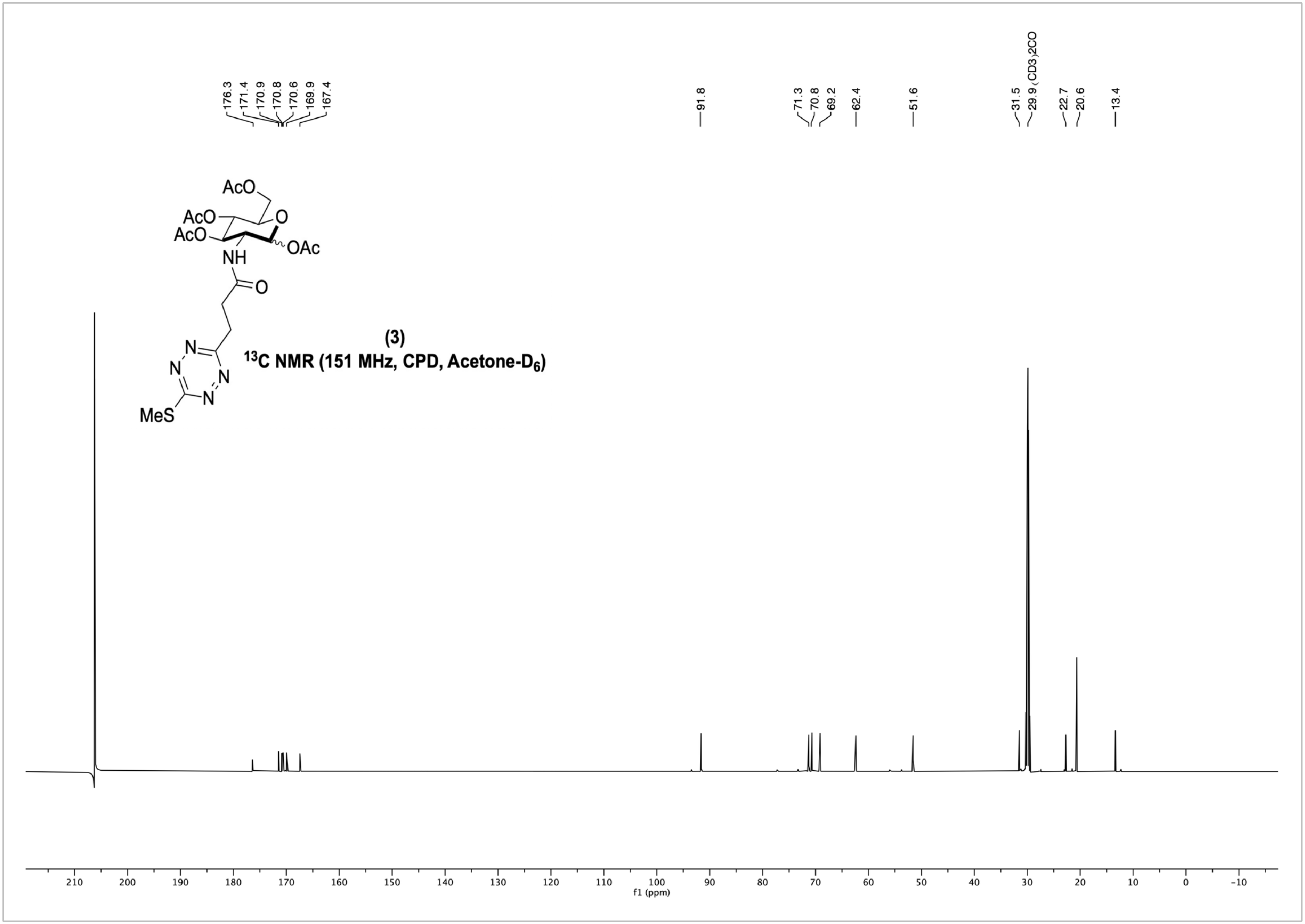


(3)

$^{13}$C NMR (151 MHz, CPD, Acetone-$D_6$)

**SMe-Tz-GlcNAc (3) 2D NMR- COSY (Acetone-D6)**

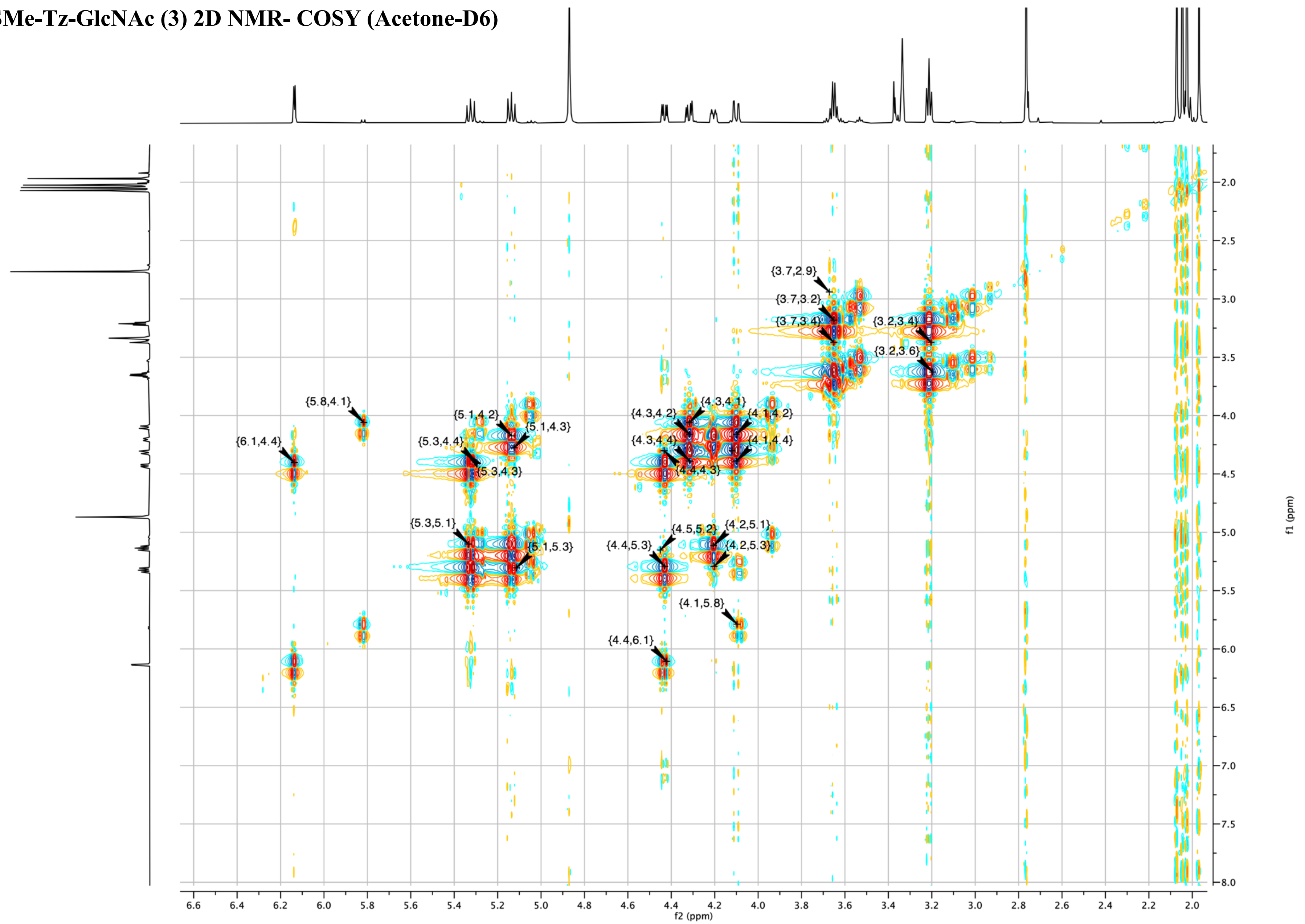

SMe-Tz-GlcNAc (3) 2D NMR- HSQC (Acetone-D6)

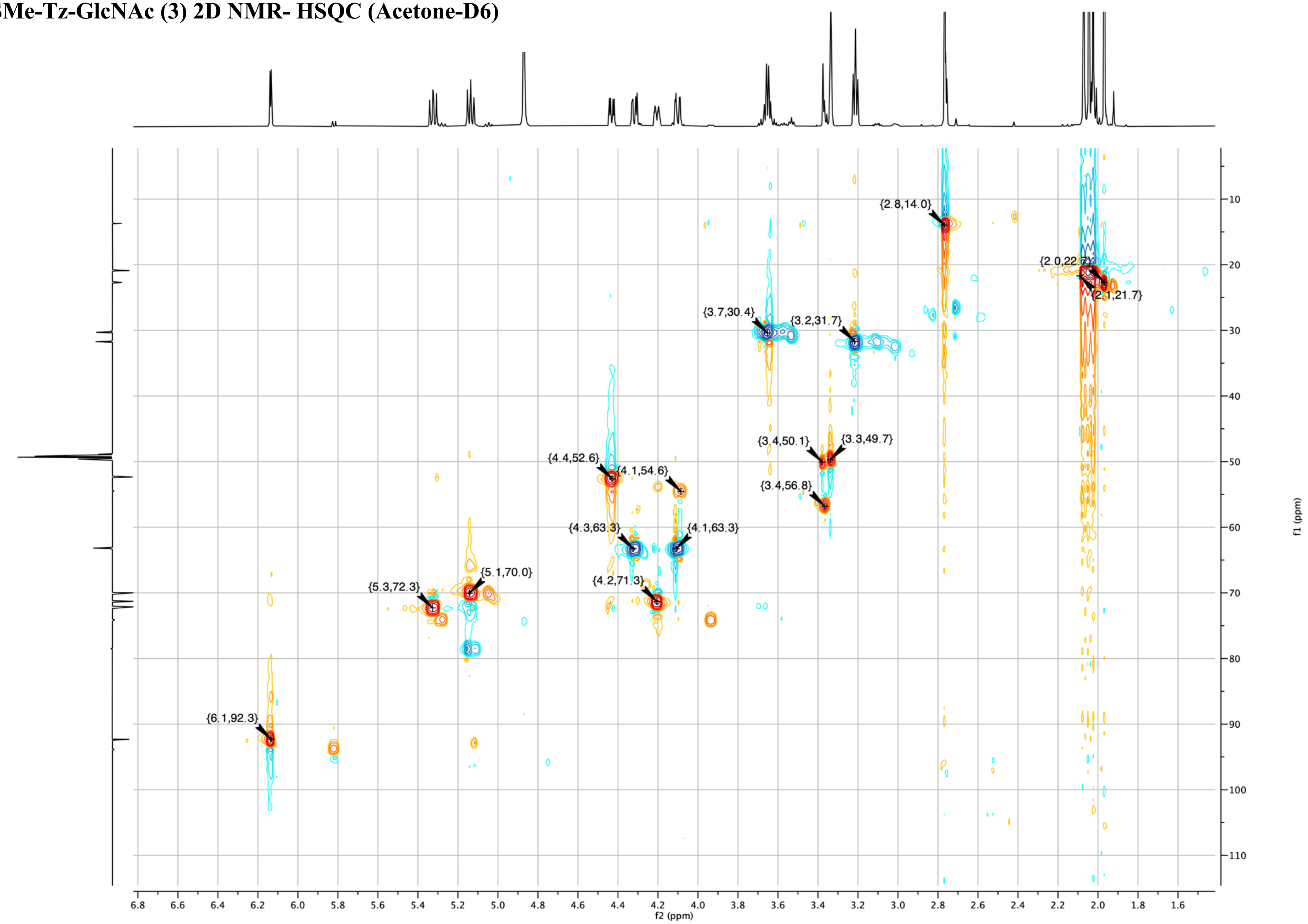

**SMe-Tz-GlcNAc (3)**
**2D NMR- HMBC**
**(Acetone-D6)**

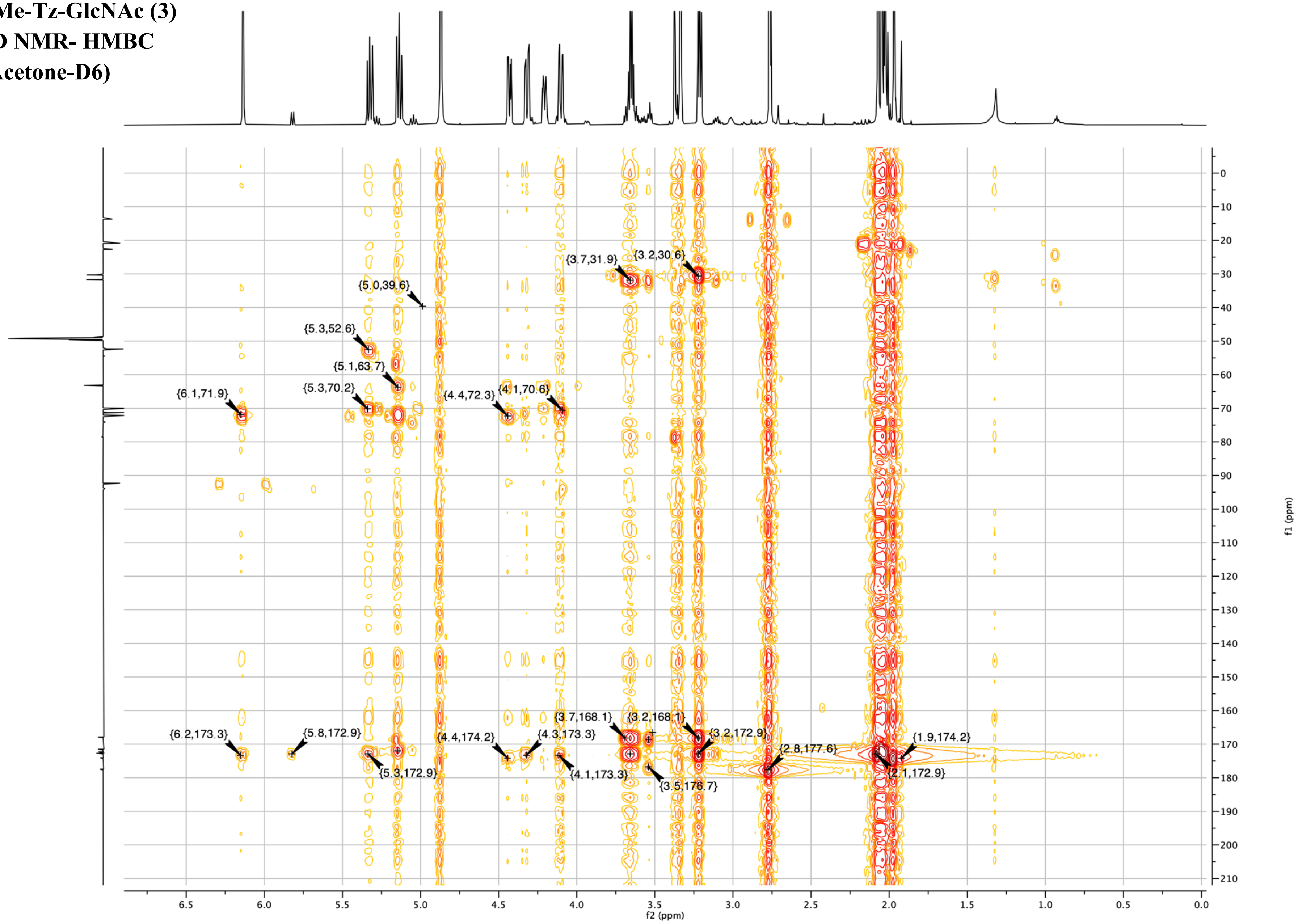

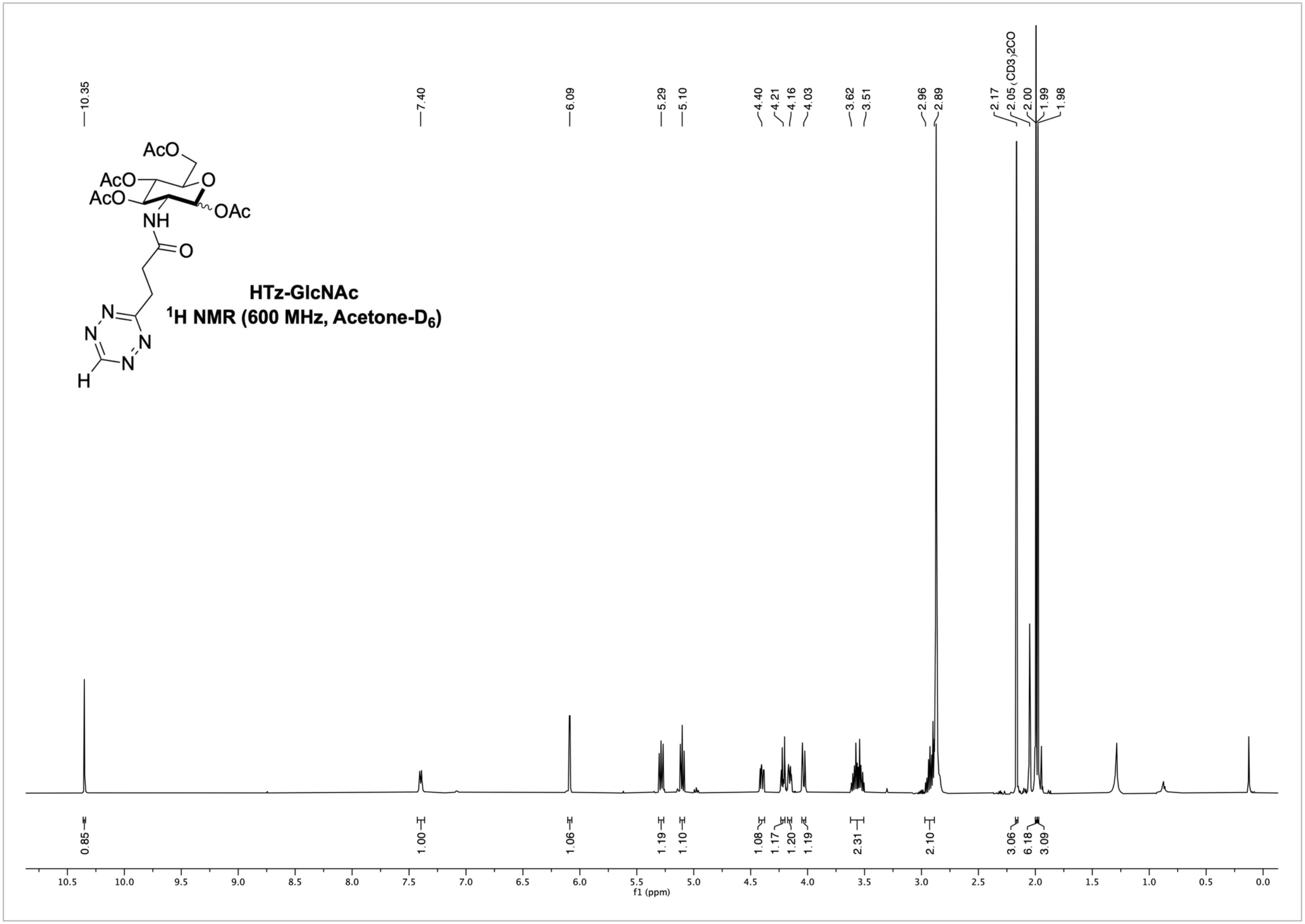
HTz-GlcNAc
$^1$H NMR (600 MHz, Acetone-D$_6$)
AcO
AcO
AcO
O
OAc
NH
O
N
N
N
N
H
10.35
7.40
6.09
5.29
5.10
4.40
4.21
4.16
4.03
3.62
3.51
2.96
2.89
2.17
2.05 (CD3)2CO
2.00
1.99
1.98
0.85
1.00
1.06
1.19
1.10
1.08
1.17
1.20
1.19
2.31
2.10
3.06
6.18
3.09
f1 (ppm)

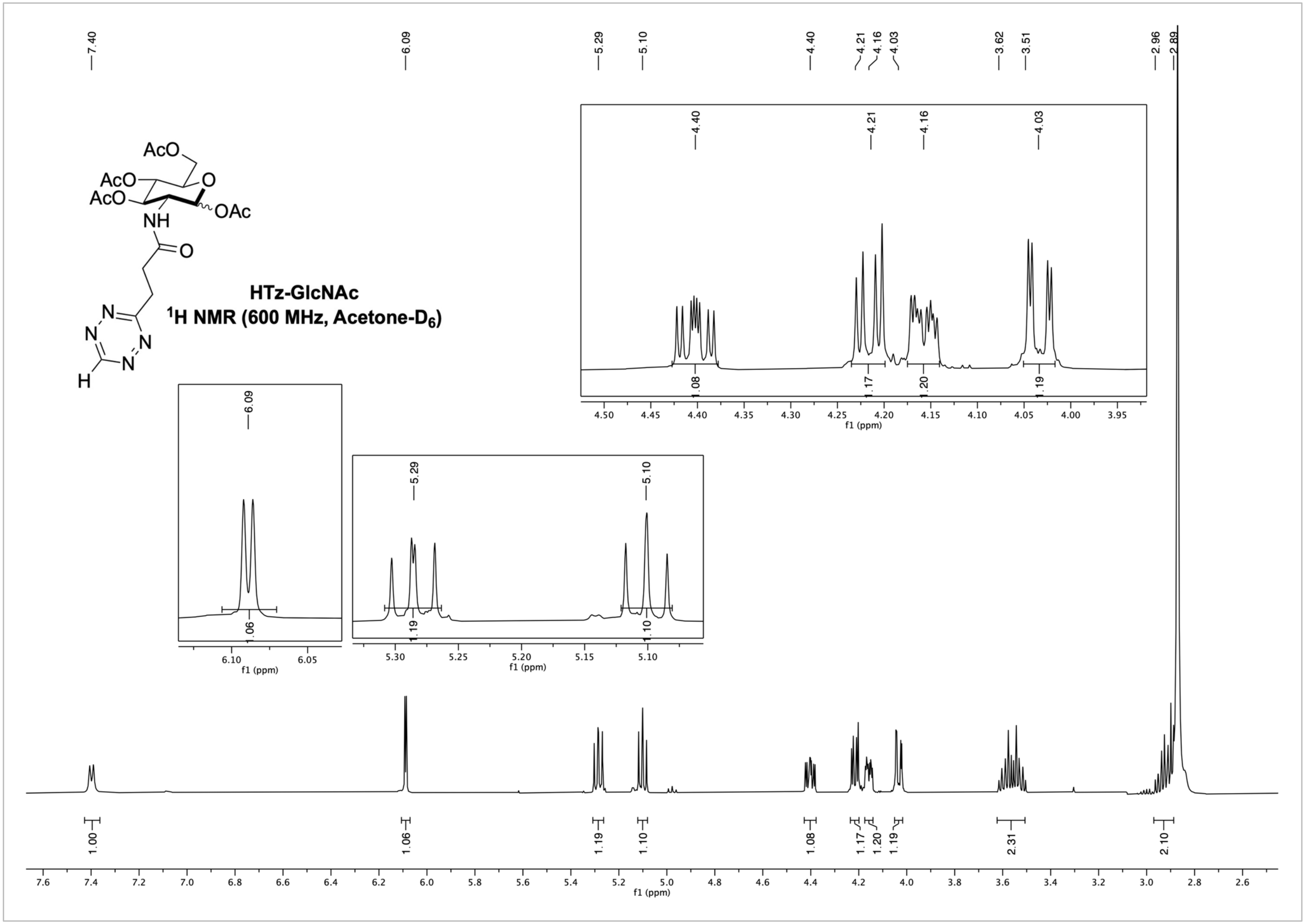
HTz-GlcNAc
1H NMR (600 MHz, Acetone-D6)
AcO
AcO
AcO
O
OAc
NH
O
N
N
N
N
H
7.40
6.09
5.29
5.10
4.40
4.21
4.16
4.03
3.62
3.51
2.96
2.89
1.00
1.06
1.19
1.10
1.08
1.17
1.20
1.19
2.31
2.10
f1 (ppm)

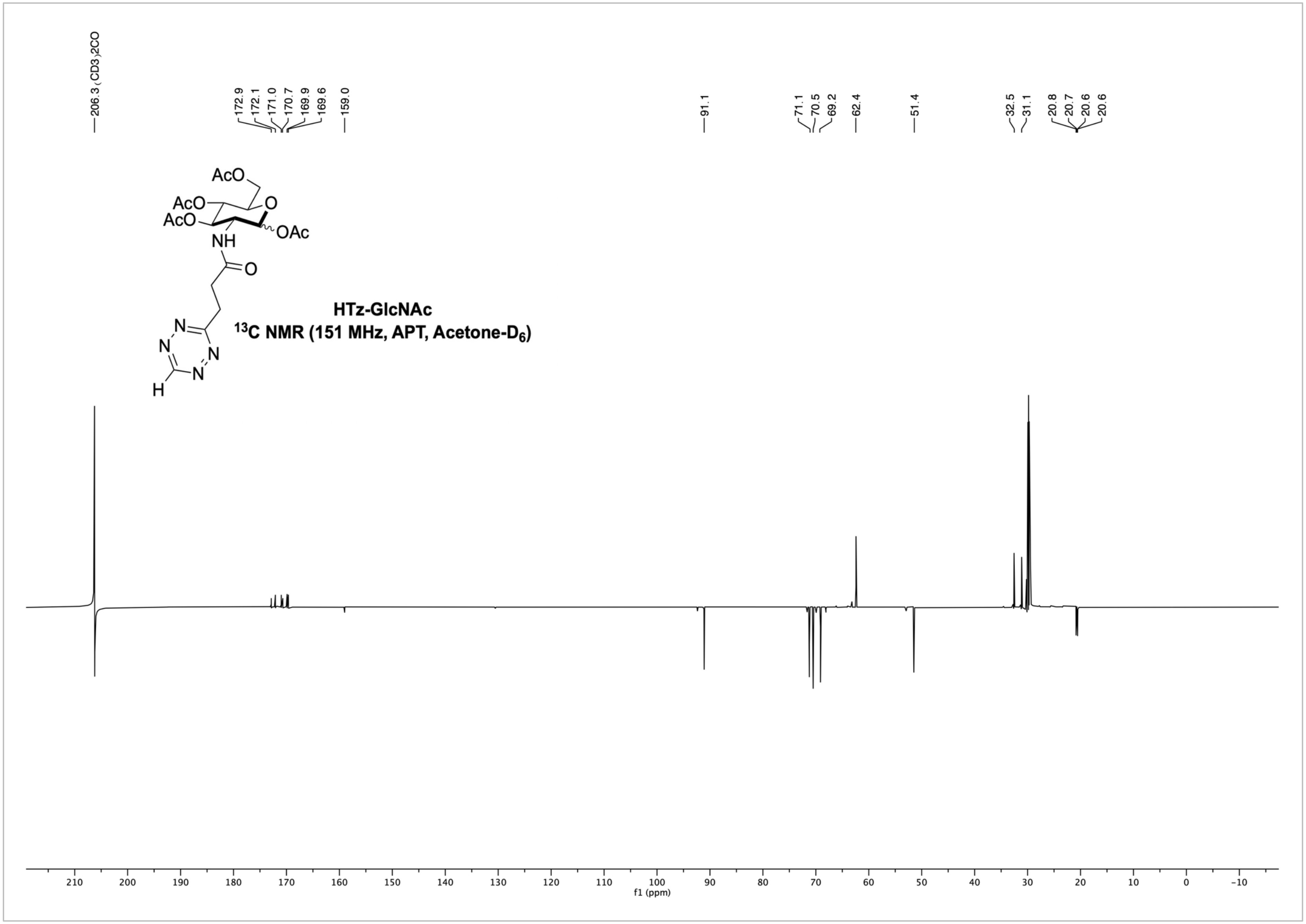
206.3 (CD3)2CO
172.9
172.1
171.0
170.7
169.9
169.6
159.0
91.1
71.1
70.5
69.2
62.4
51.4
32.5
31.1
20.8
20.7
20.6
20.6
AcO
O
OAc
NH
N
H
HTz-GlcNAc
13C NMR (151 MHz, APT, Acetone-D6)
210
200
190
180
170
160
150
140
130
120
110
100
90
80
70
60
50
40
30
20
10
0
-10
f1 (ppm)

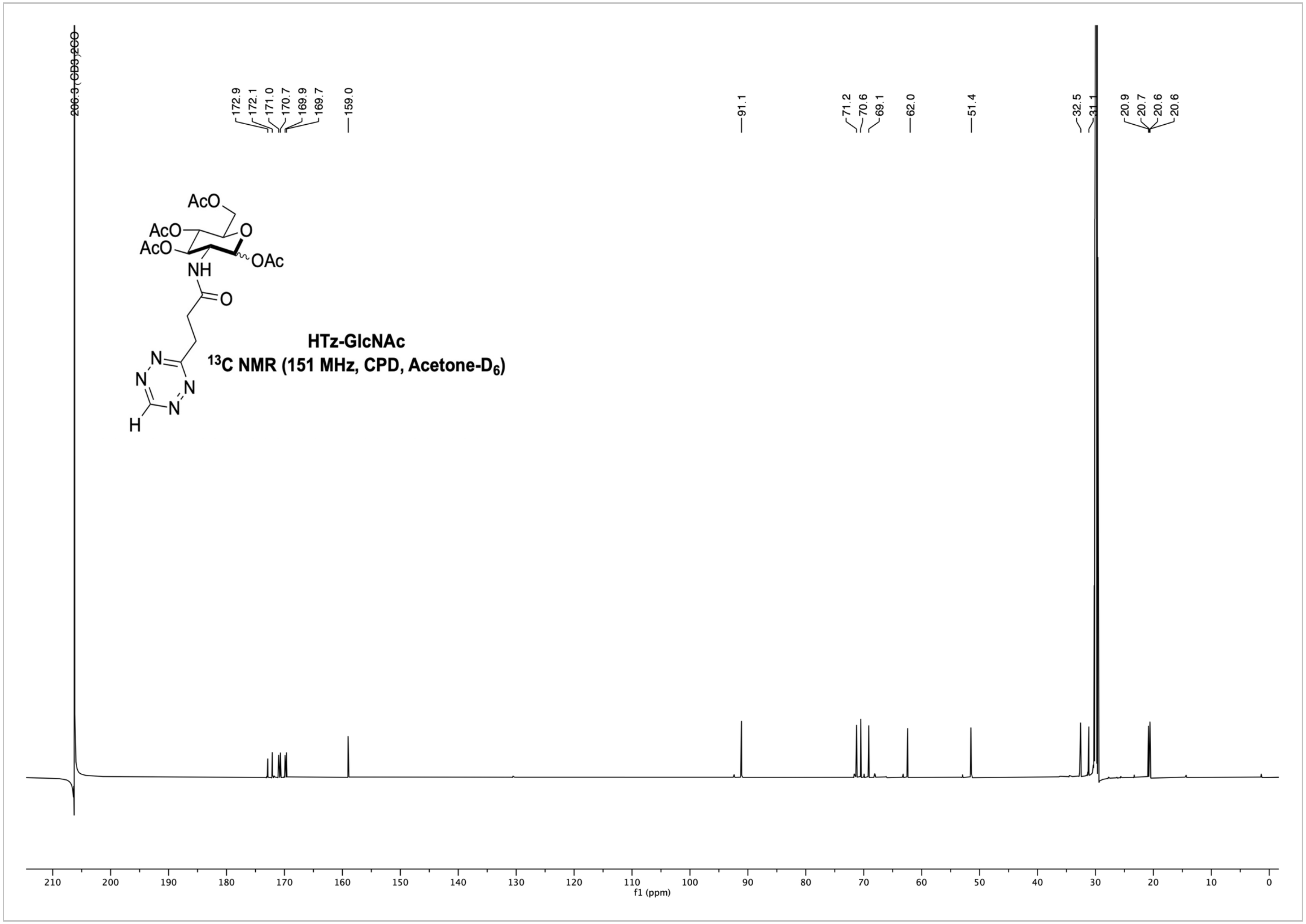
206.3 (CD3)2CO
172.9
172.1
171.0
170.7
169.9
169.7
159.0
91.1
71.2
70.6
69.1
62.0
51.4
32.5
31.1
20.9
20.7
20.6
20.6
AcO
AcO
AcO
O
OAc
NH
O
N
N
N
N
H
HTz-GlcNAc
13C NMR (151 MHz, CPD, Acetone-D6)
f1 (ppm)

HTz-GlcNAc 2D NMR- COSY (Acetone-D6)

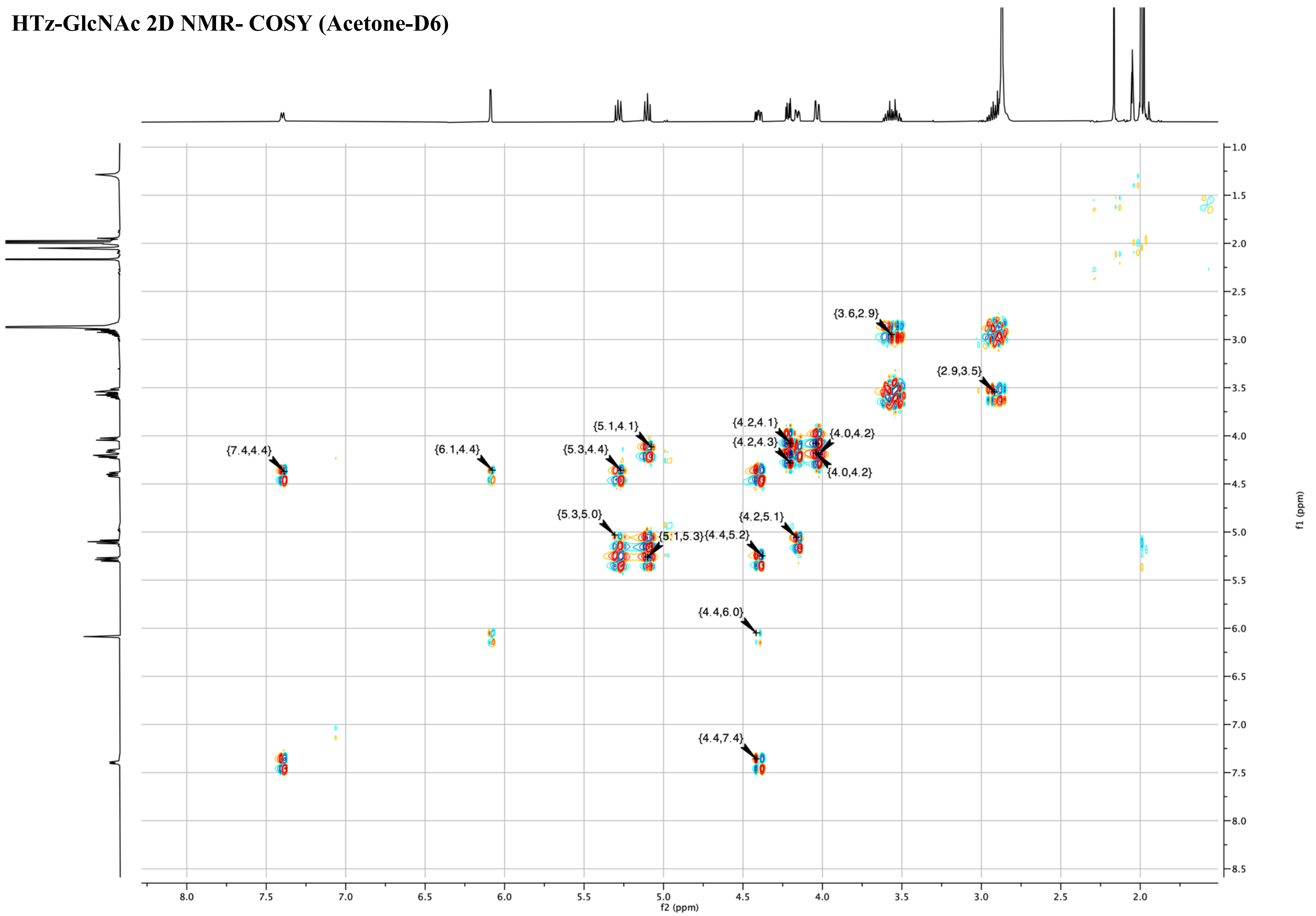

**HTz-GlcNAc 2D NMR- HSQC (Acetone-D6)**

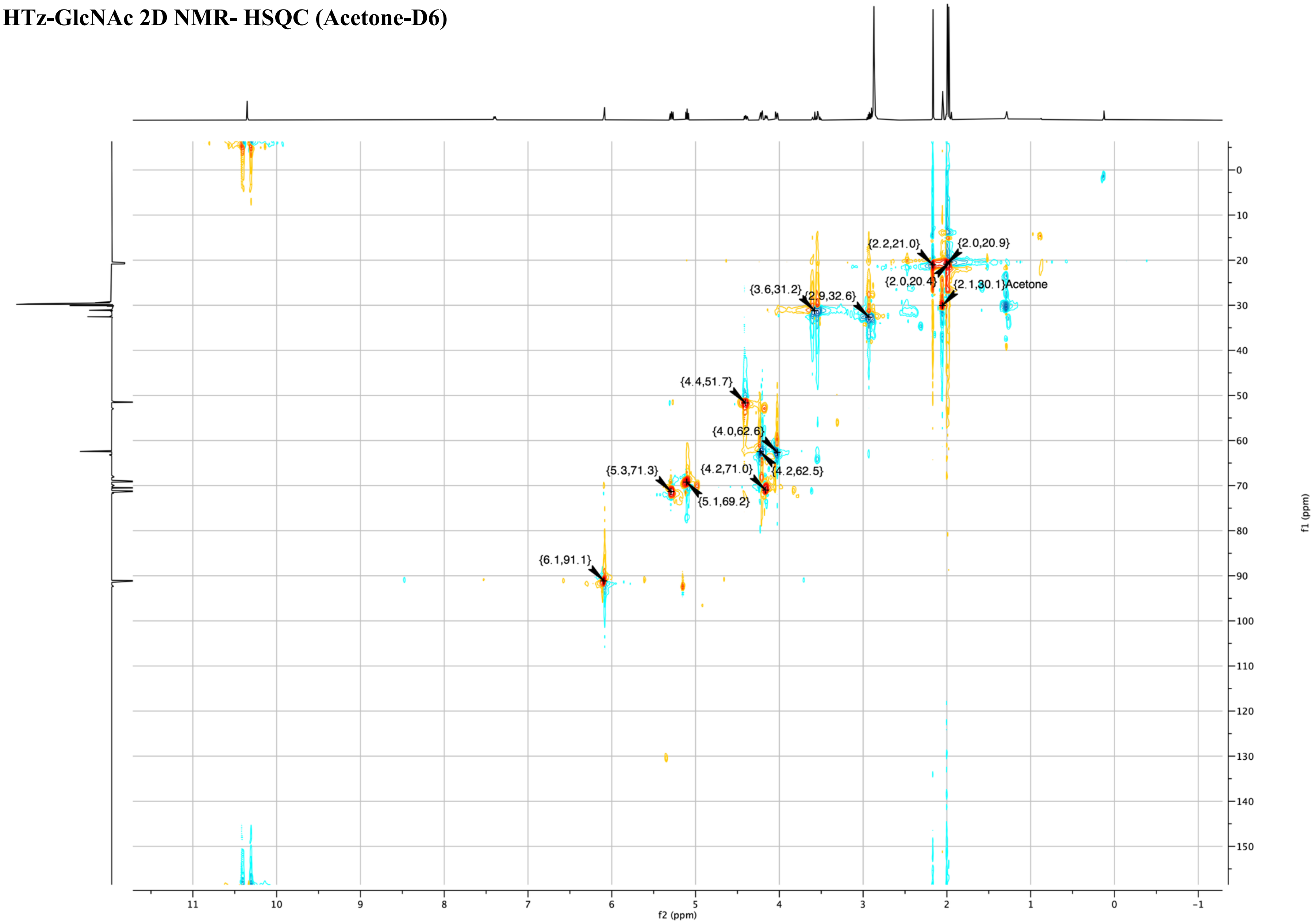

### HTz-GlcNAc 2D NMR- HMBC (Acetone-D6)

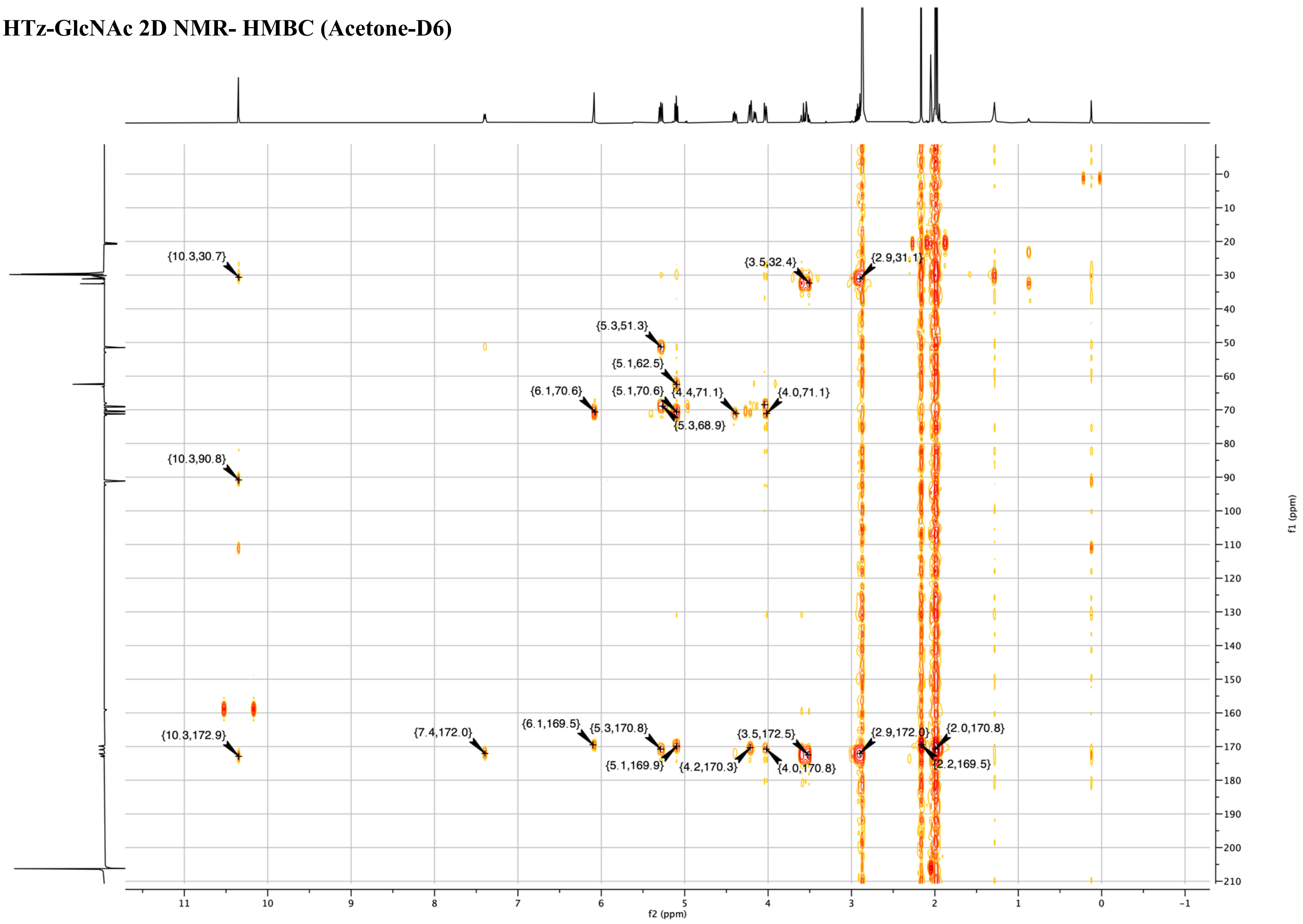

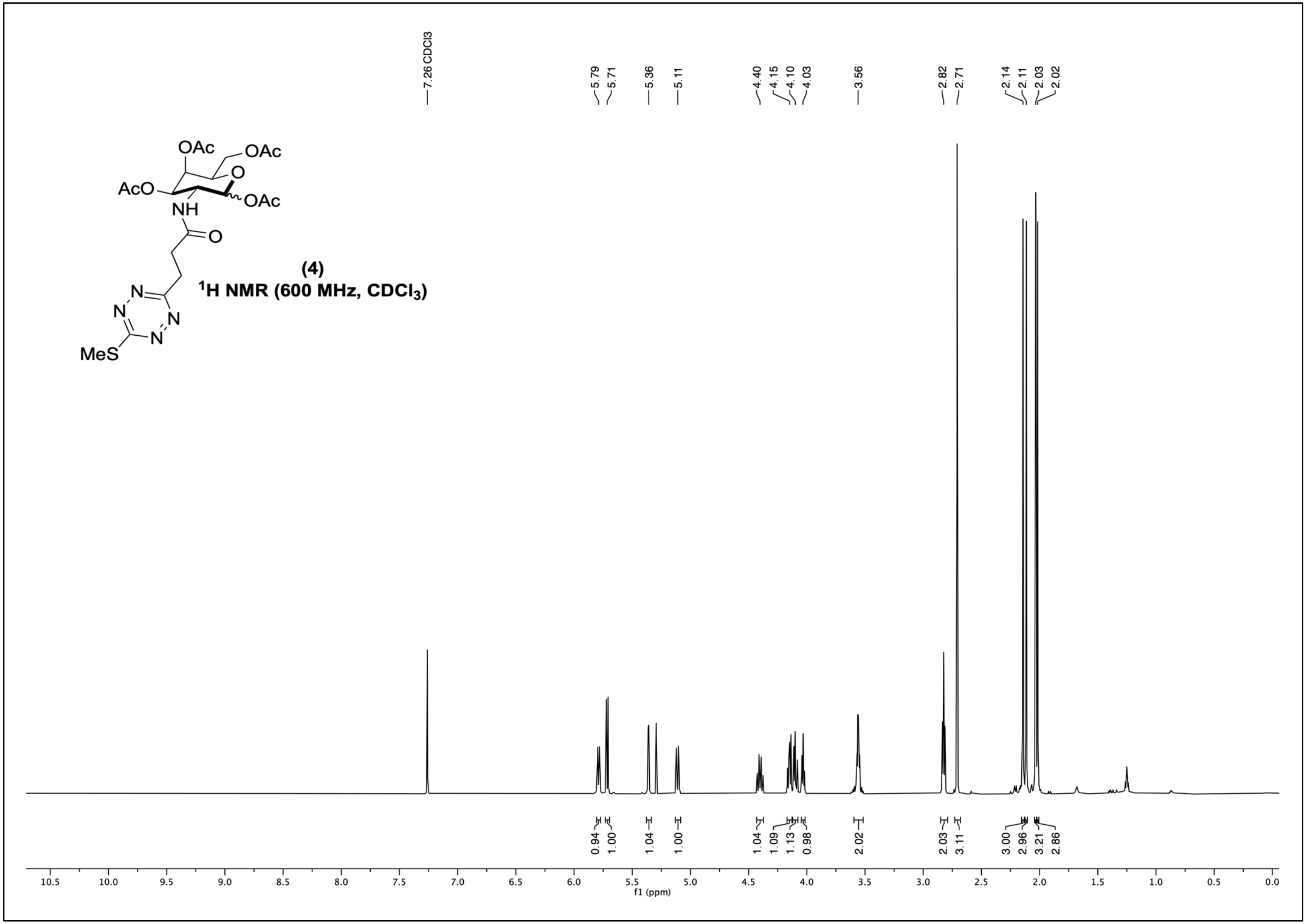

(4)
¹H NMR (600 MHz, CDCl₃)
7.26 CDCl3
5.79
5.71
5.36
5.11
4.40
4.15
4.10
4.03
3.56
2.82
2.71
2.14
2.11
2.03
2.02
0.94
1.00
1.04
1.00
1.04
1.09
1.13
0.98
2.02
2.03
3.11
3.00
2.96
3.21
2.86
f1 (ppm)

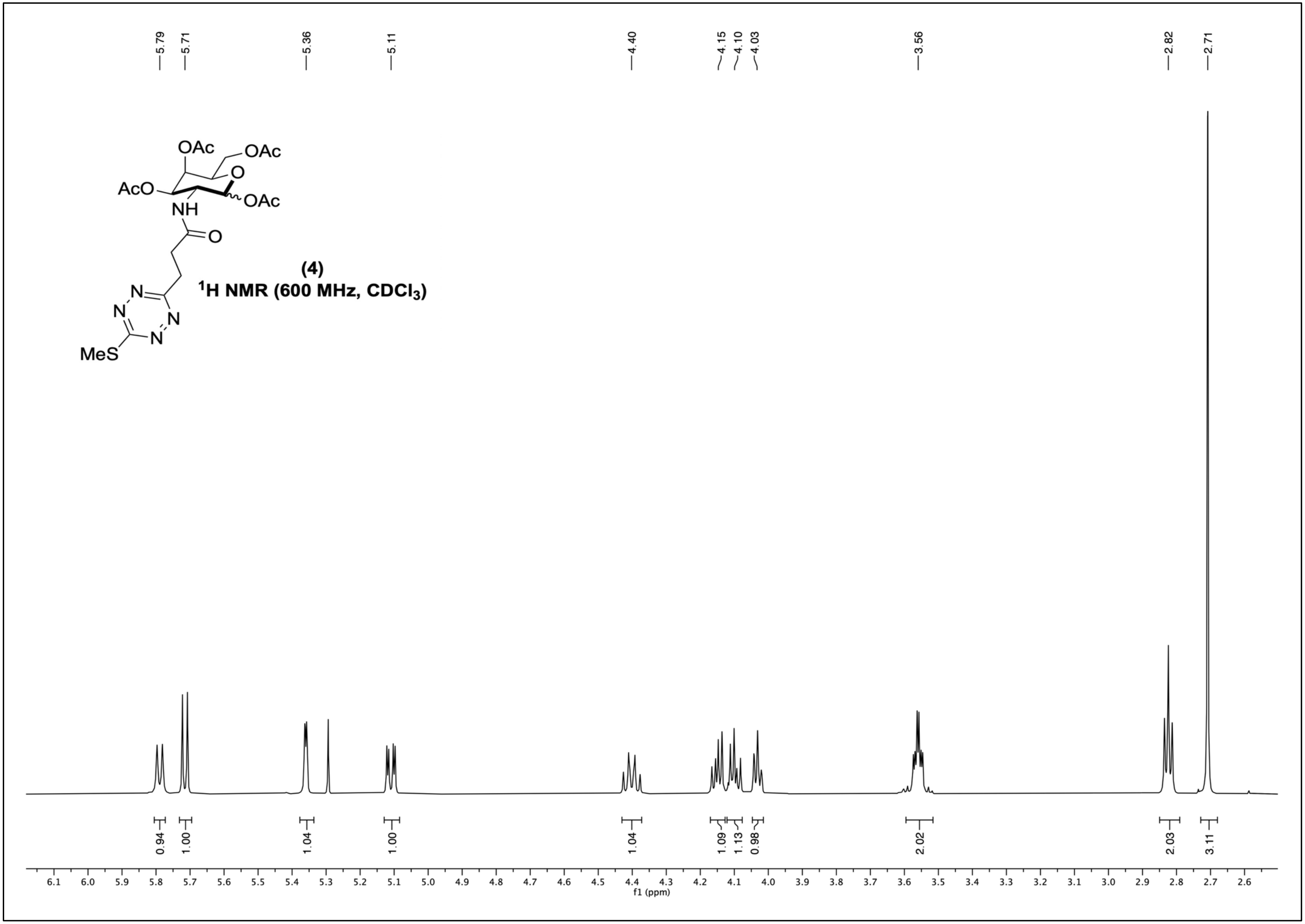

5.79
5.71
5.36
5.11
4.40
4.15
4.10
4.03
3.56
2.82
2.71
OAc
OAc
O
AcO
OAc
NH
O
N
N
N
N
MeS
(4)
1H NMR (600 MHz, CDCl3)
0.94
1.00
1.04
1.00
1.04
1.09
1.13
0.98
2.02
2.03
3.11
6.1 6.0 5.9 5.8 5.7 5.6 5.5 5.4 5.3 5.2 5.1 5.0 4.9 4.8 4.7 4.6 4.5 4.4 4.3 4.2 4.1 4.0 3.9 3.8 3.7 3.6 3.5 3.4 3.3 3.2 3.1 3.0 2.9 2.8 2.7 2.6
f1 (ppm)

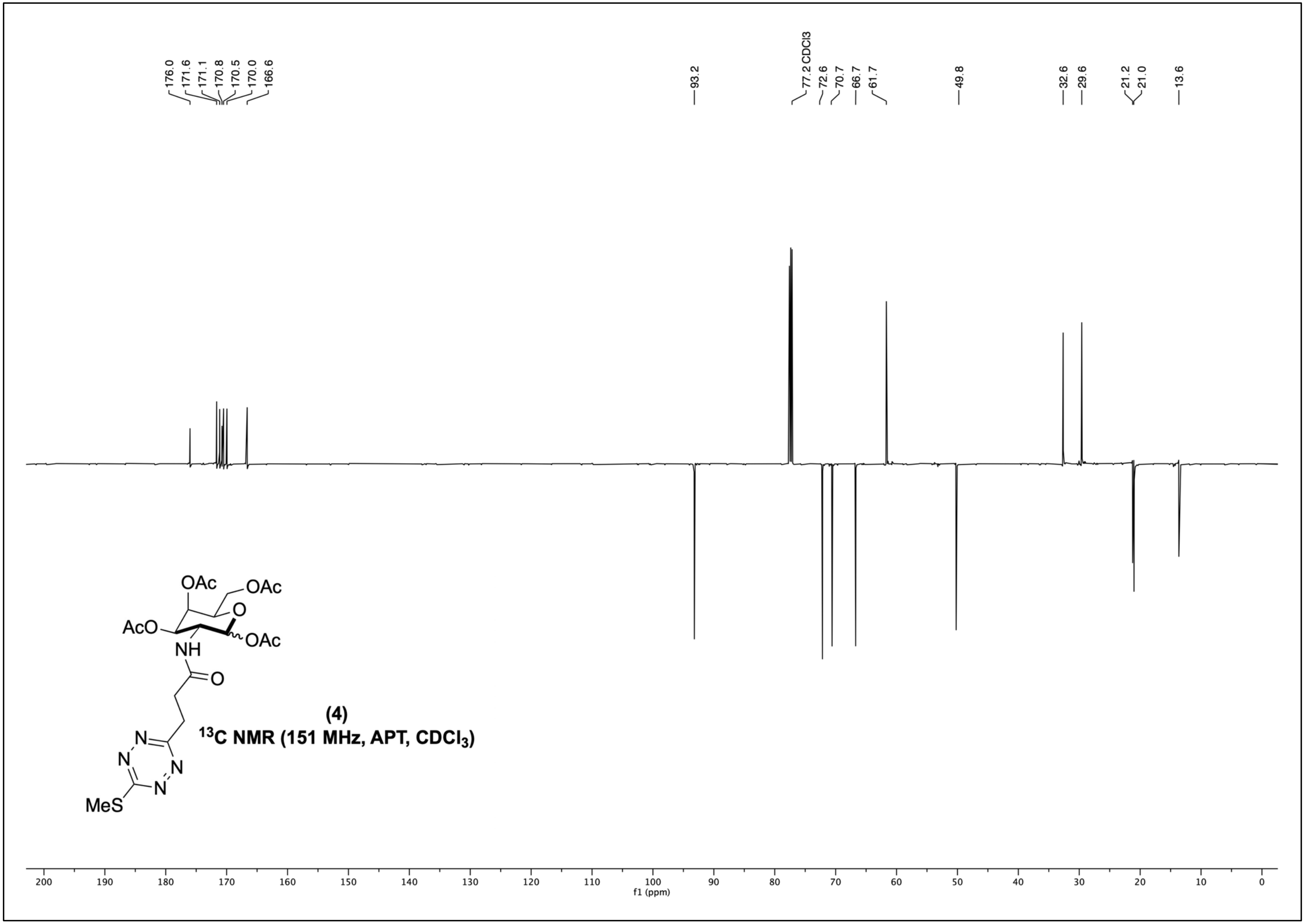
176.0
171.6
171.1
170.8
170.5
170.0
166.6
93.2
77.2 CDCl3
72.6
70.7
66.7
61.7
49.8
32.6
29.6
21.2
21.0
13.6
OAc
OAc
AcO
NH
OAc
O
O
N
N
N
N
MeS
(4)
$^{13}$C NMR (151 MHz, APT, $CDCl_3$)
200
190
180
170
160
150
140
130
120
110
100
f1 (ppm)
90
80
70
60
50
40
30
20
10
0

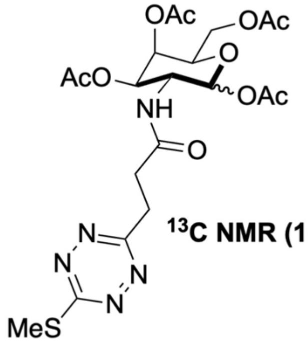

(4)

$^{13}$C NMR (151 MHz, CPD, $CDCl_3$)

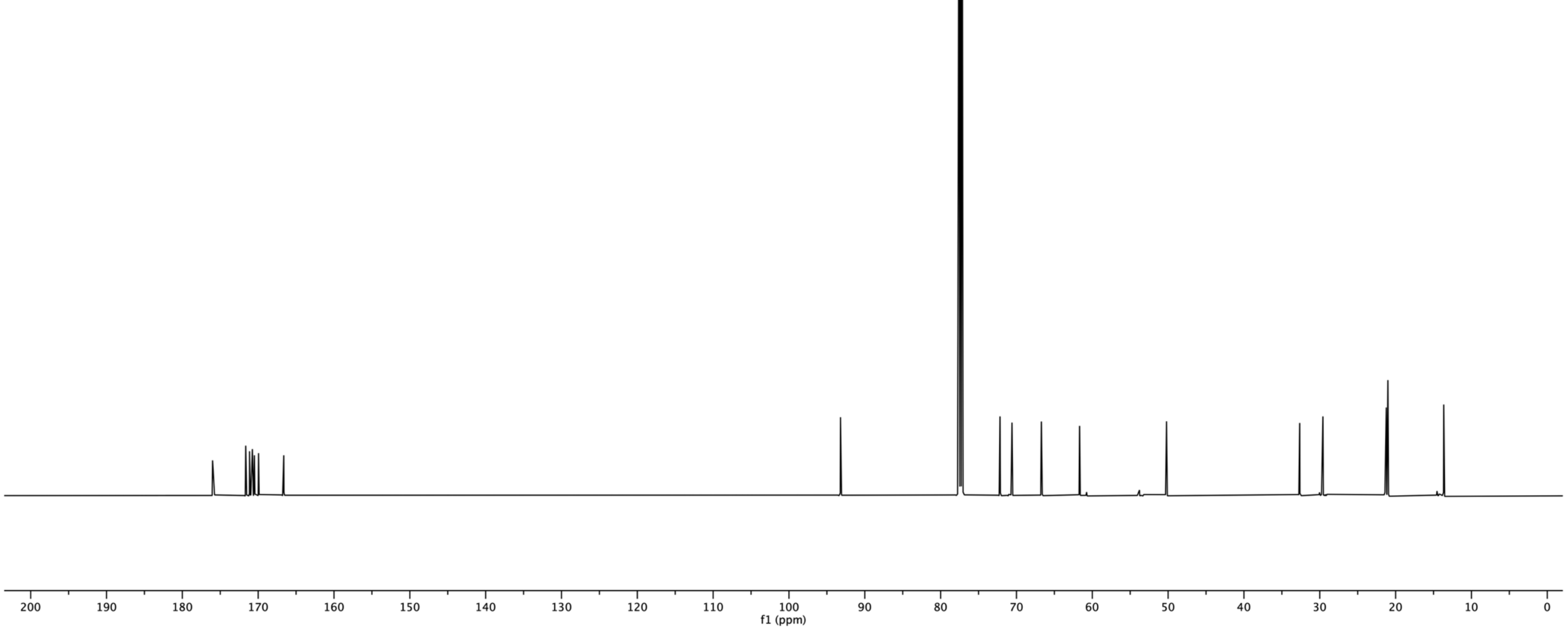

SMe-Tz-GalNAc (4) 2D NMR- COSY (Acetone-D6)

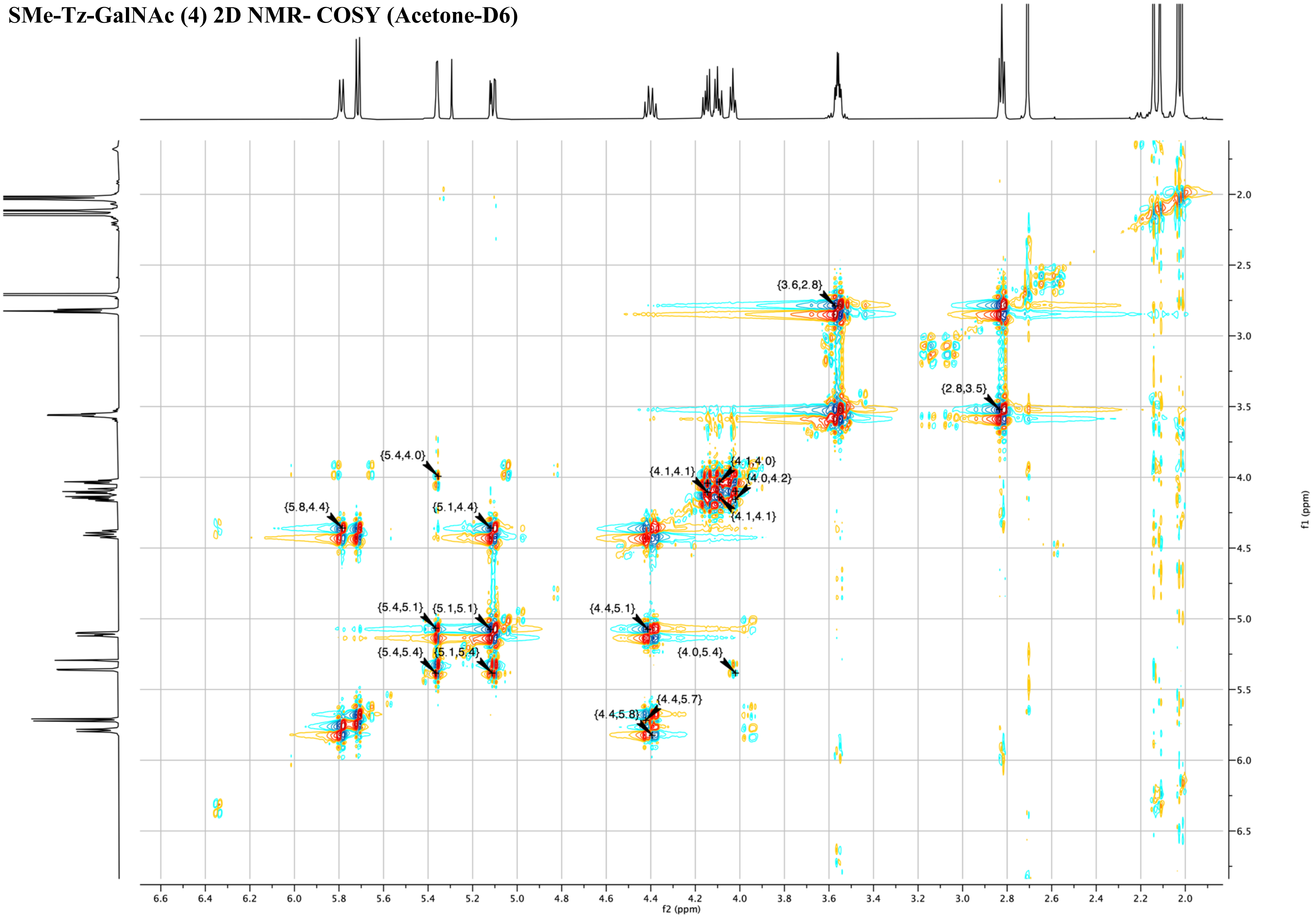

**SMe-Tz-GalNAc (4) 2D NMR- HSQC (Acetone-D6)**

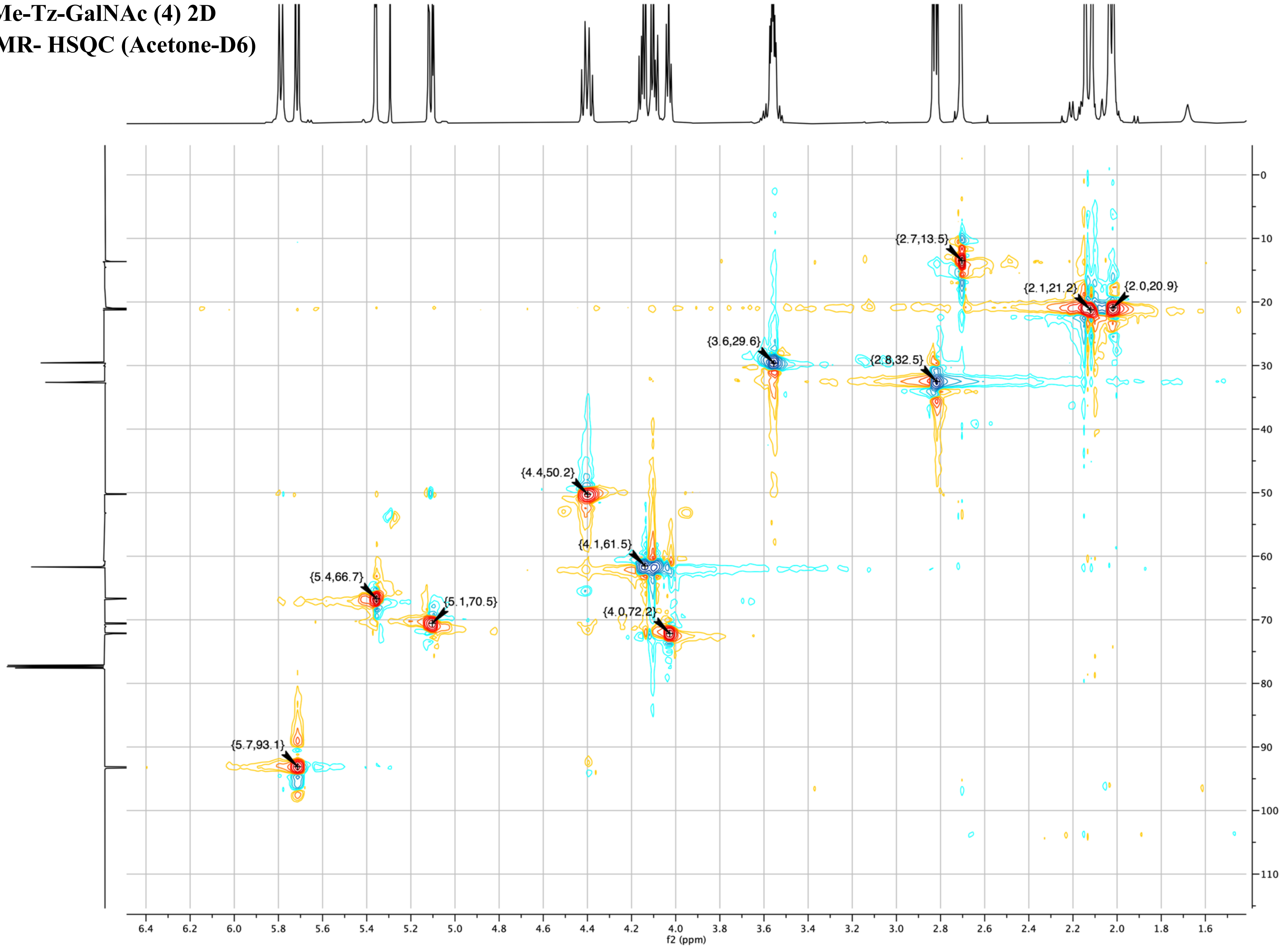

**SMe-Tz-GalNAc (4)**
**2D NMR- HMBC**
**(Acetone-D6)**

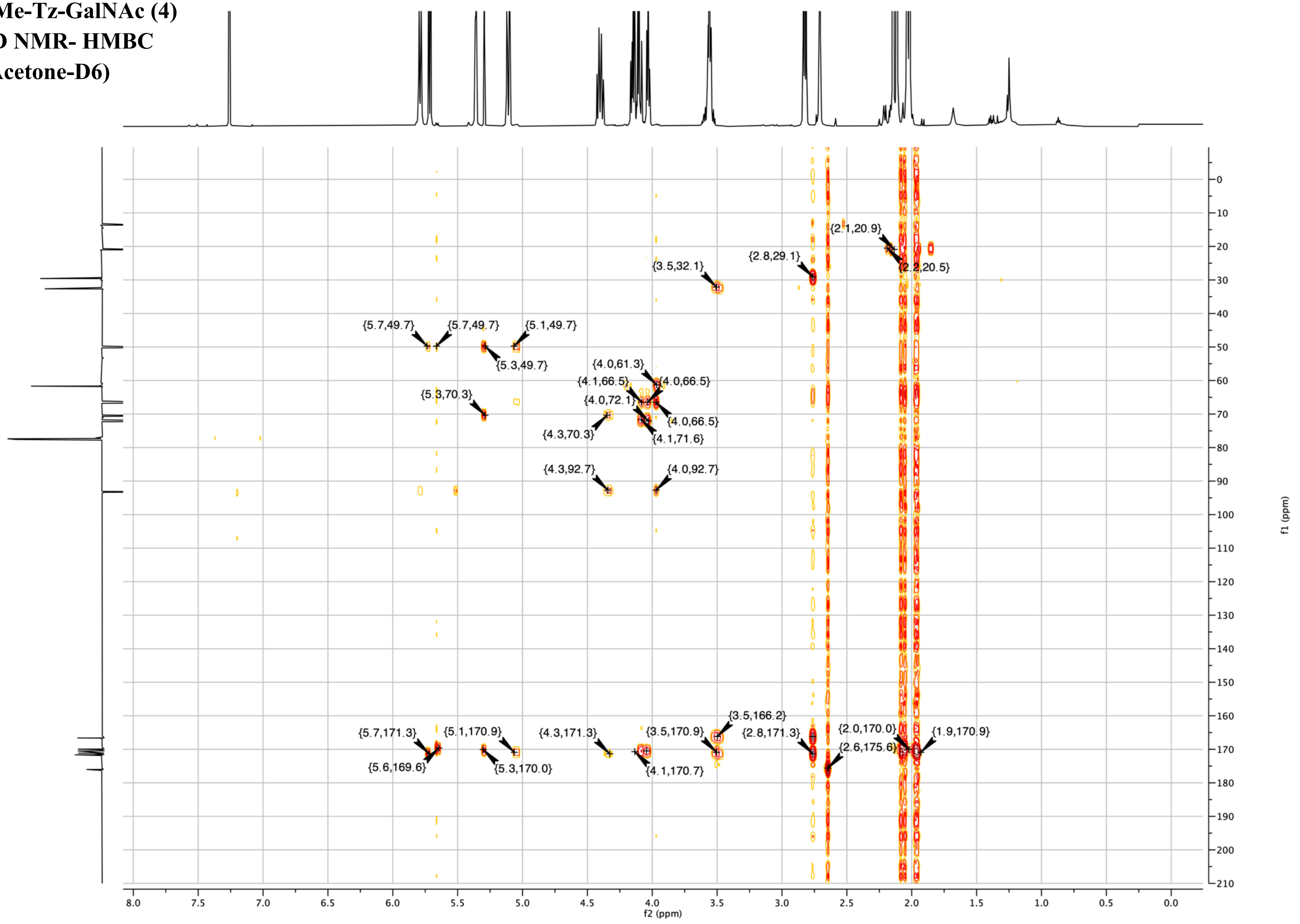

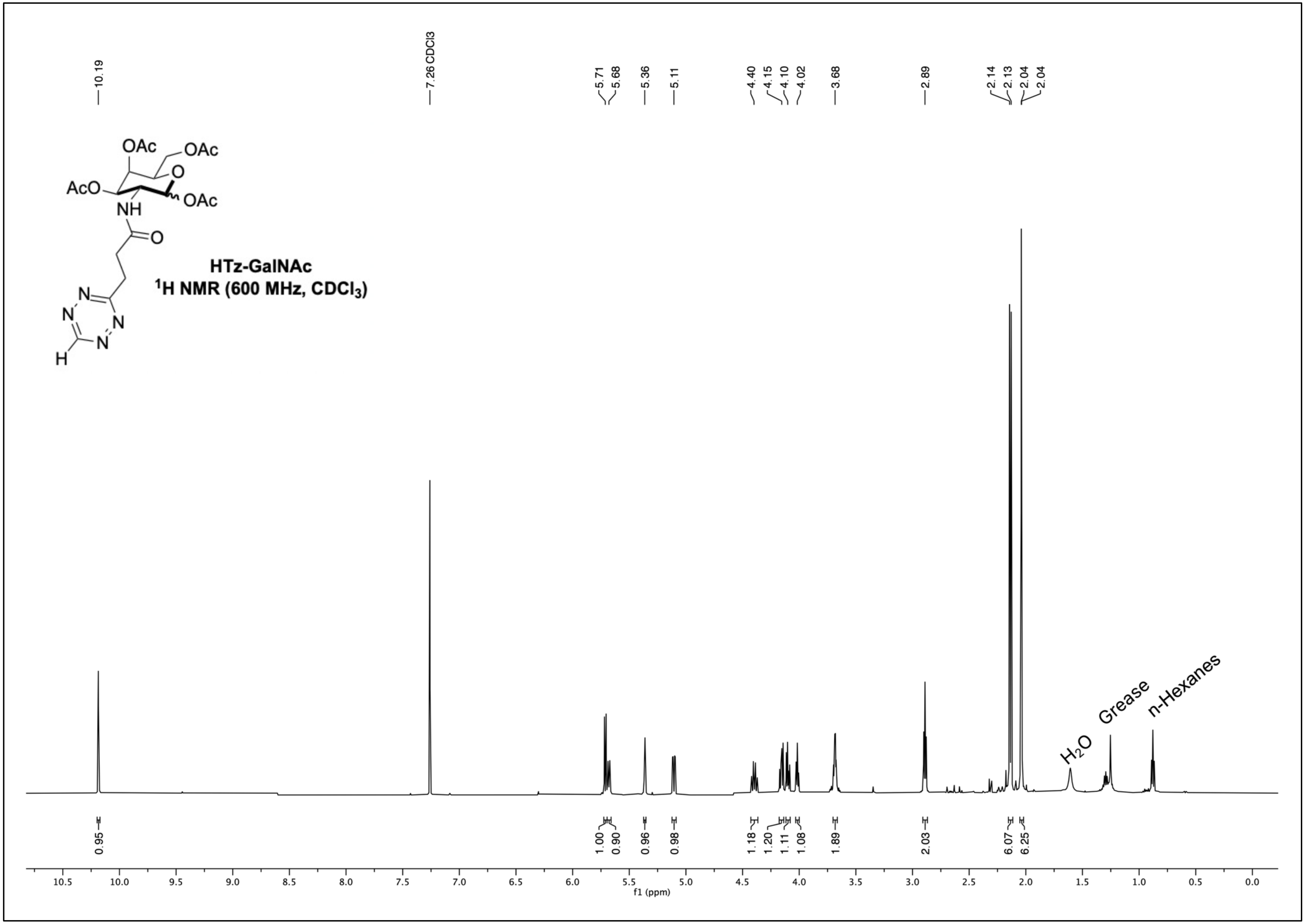
HTz-GalNAc
$^{1}H$ NMR (600 MHz, $CDCl_3$)
H₂O
Grease
n-Hexanes
f1 (ppm)

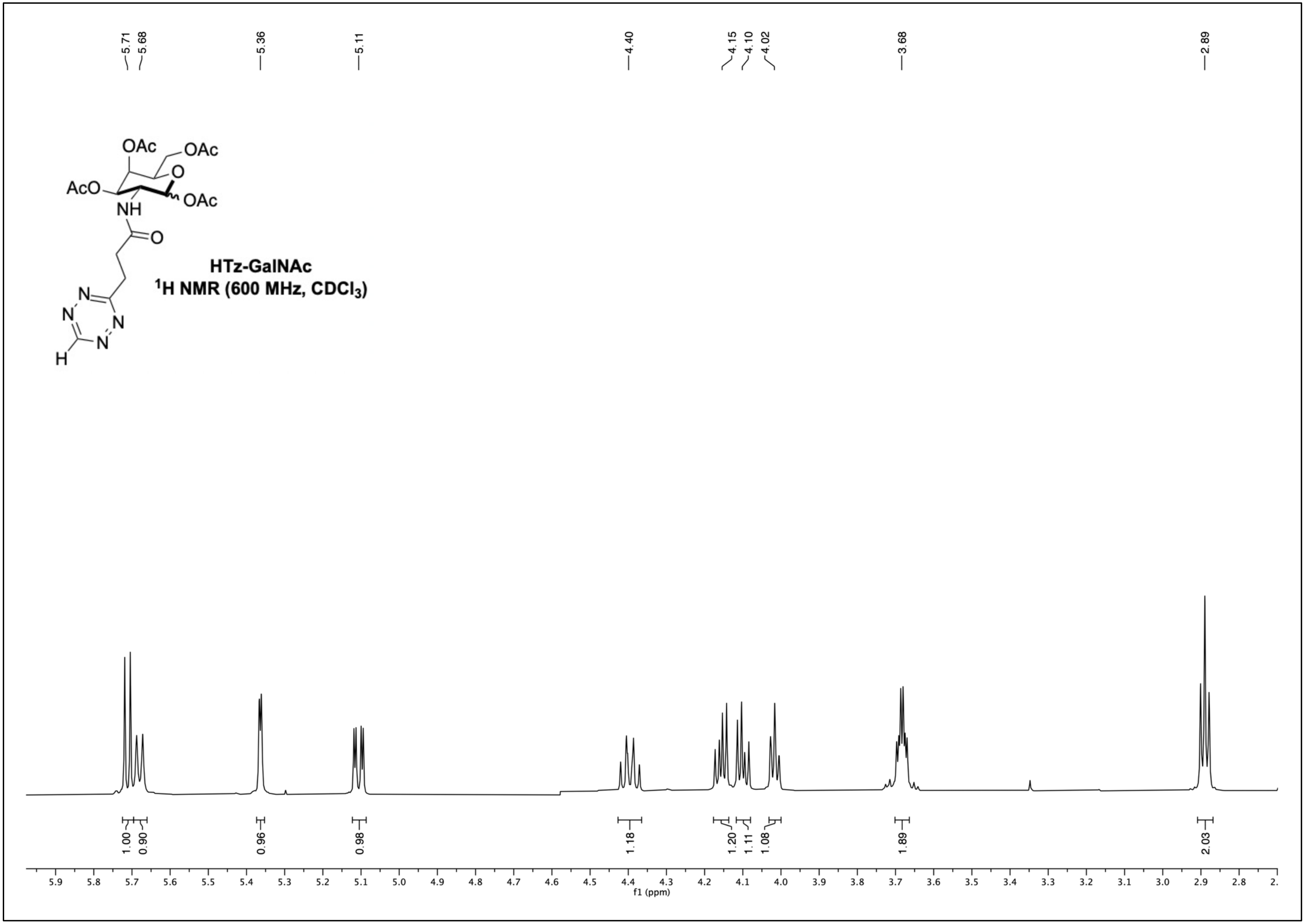
HTz-GalNAc
$^1$H NMR (600 MHz, CDCl$_3$)
5.71
5.68
5.36
5.11
4.40
4.15
4.10
4.02
3.68
2.89
1.00
0.90
0.96
0.98
1.18
1.20
1.11
1.08
1.89
2.03
f1 (ppm)

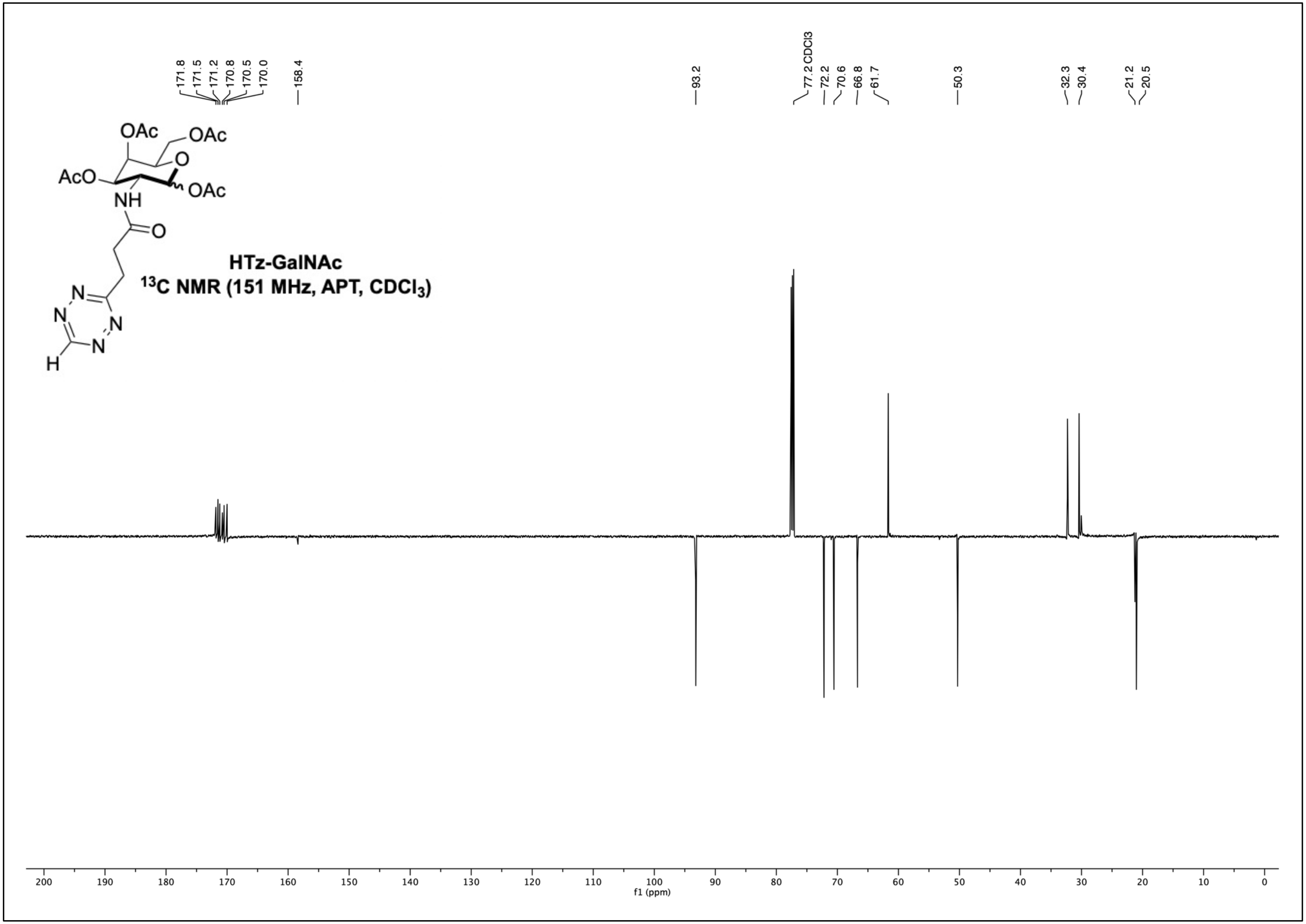
171.8
171.5
171.2
170.8
170.5
170.0
158.4
93.2
77.2 CDCl3
72.2
70.6
66.8
61.7
50.3
32.3
30.4
21.2
20.5
OAc
OAc
AcO
O
OAc
NH
O
N
N
N
N
H
HTz-GalNAc
13C NMR (151 MHz, APT, CDCl3)
200
190
180
170
160
150
140
130
120
110
100
90
80
70
60
50
40
30
20
10
0
f1 (ppm)

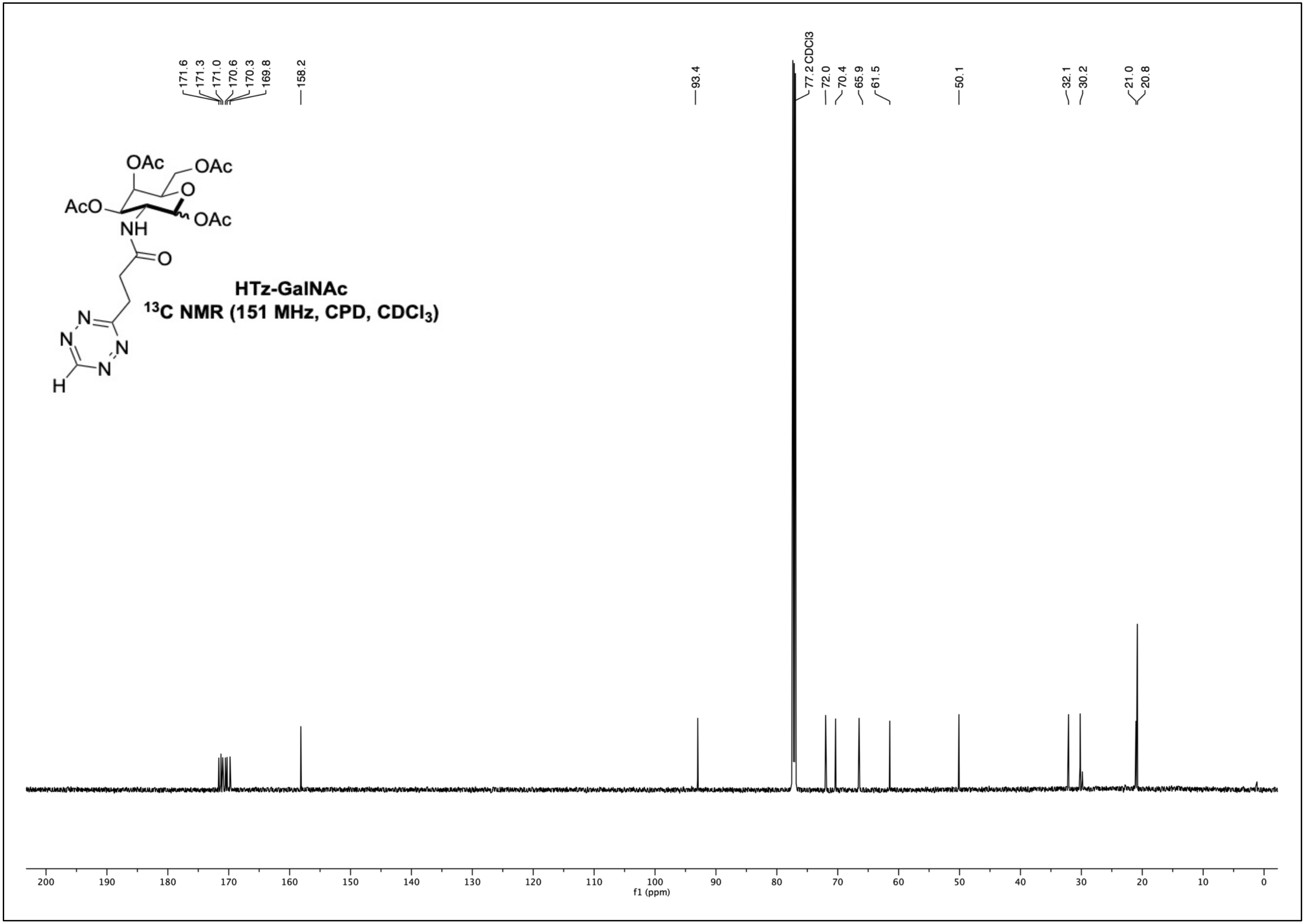
HTz-GalNAc
$^{13}$C NMR (151 MHz, CPD, $CDCl_3$)
171.6
171.3
171.0
170.6
170.3
169.8
158.2
93.4
77.2 CDCl3
72.0
70.4
65.9
61.5
50.1
32.1
30.2
21.0
20.8
f1 (ppm)

**HTz-GalNAc 2D NMR- COSY (Acetone-D6)**

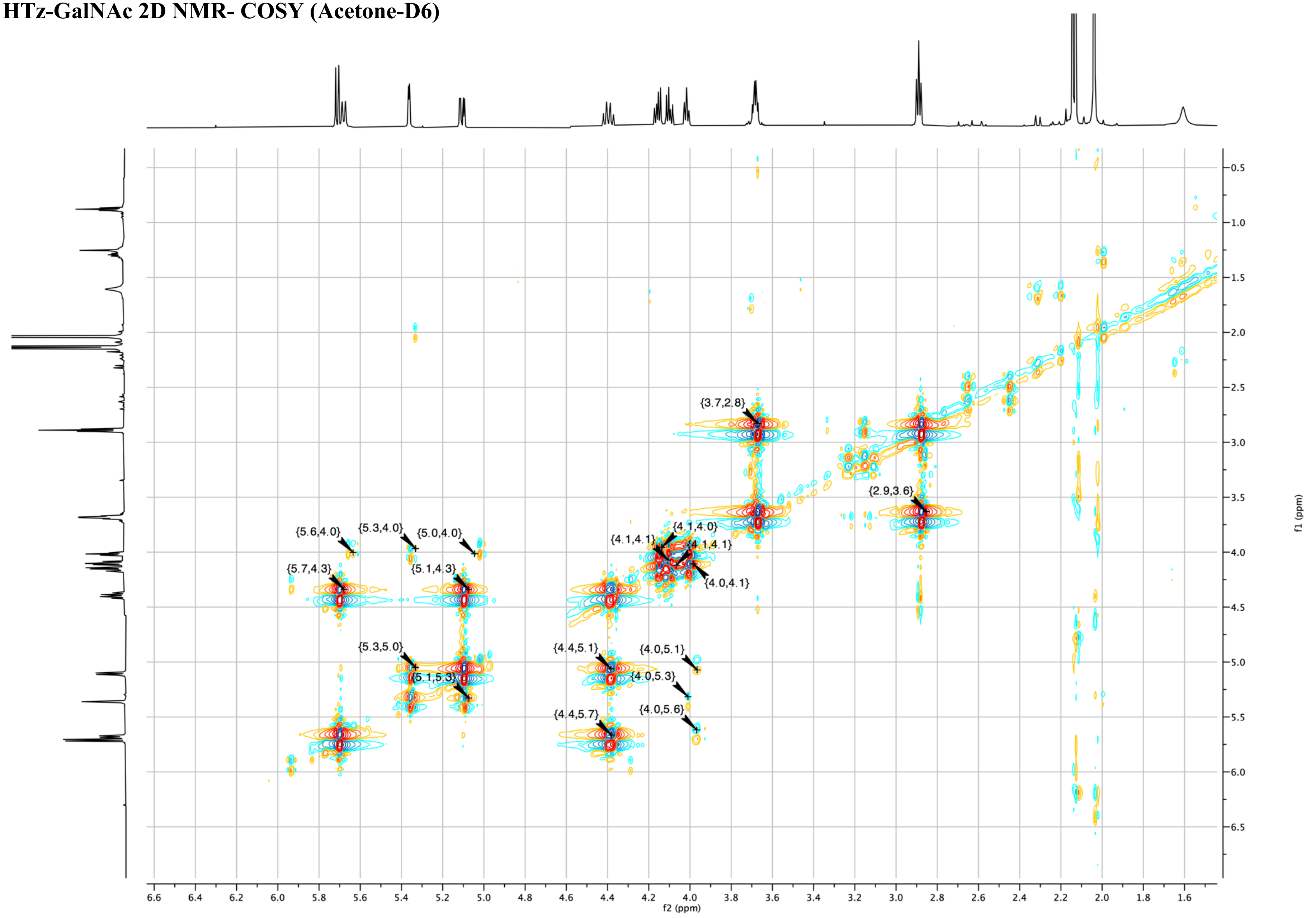

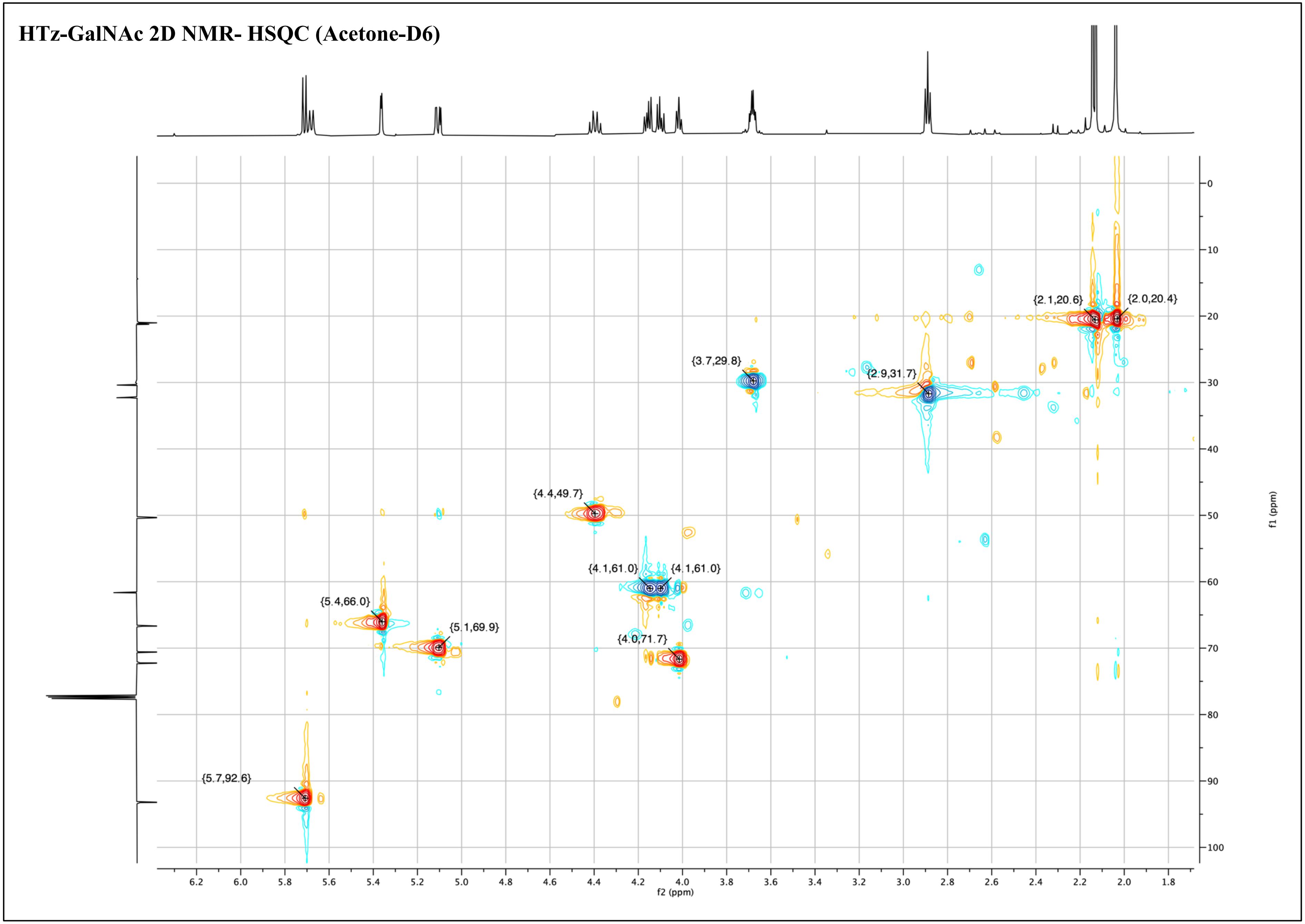
HTz-GalNAc 2D NMR- HSQC (Acetone-D6)
{2.1,20.6}
{2.0,20.4}
{3.7,29.8}
{2.9,31.7}
{4.4,49.7}
{4.1,61.0}
{4.1,61.0}
{5.4,66.0}
{5.1,69.9}
{4.0,71.7}
{5.7,92.6}
f2 (ppm)
f1 (ppm)

**HTz-GalNAc 2D NMR-HMBC (Acetone-D6)**

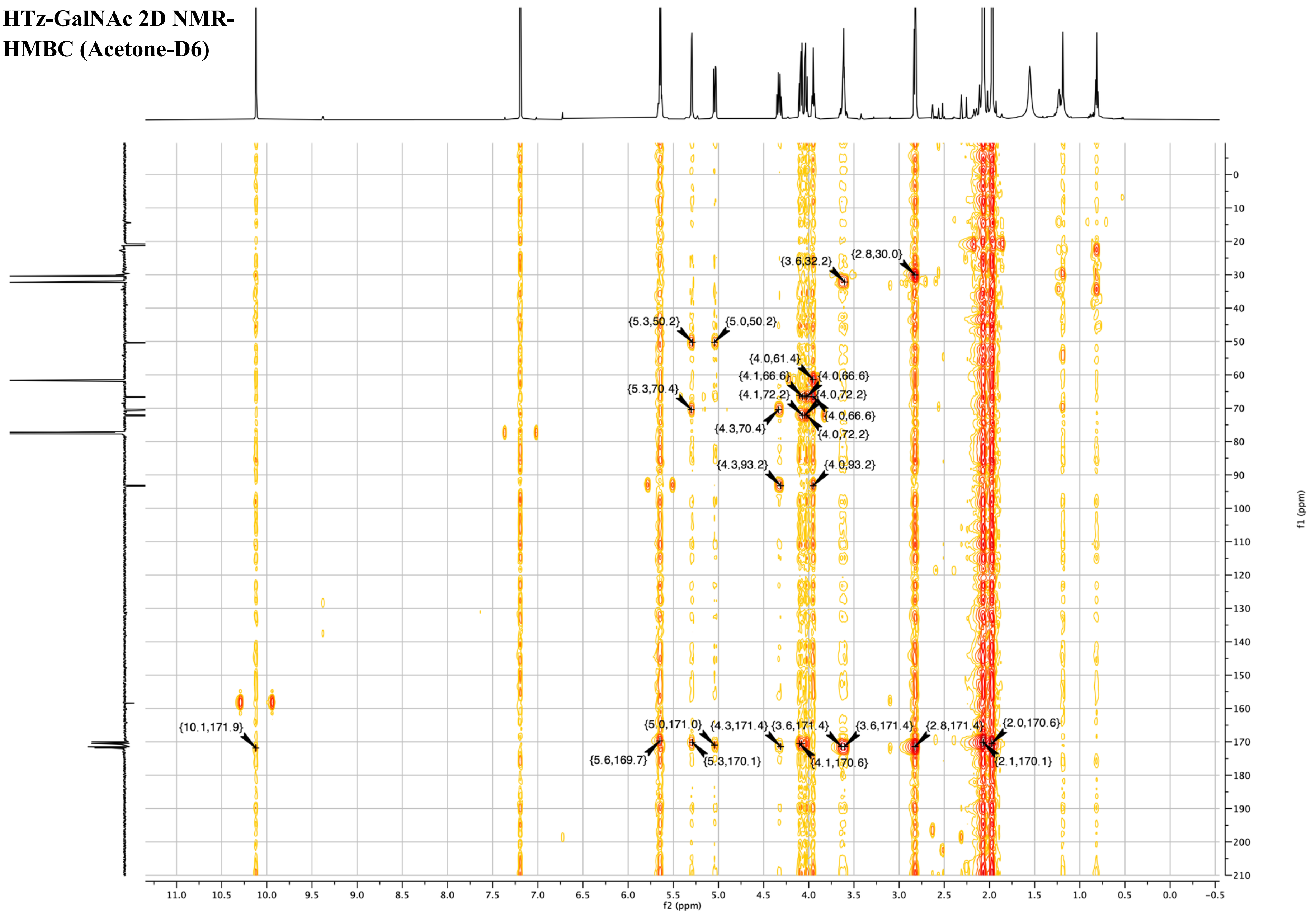
